# Effects of spatial heterogeneity on bacterial genetic circuits

**DOI:** 10.1101/2019.12.22.886473

**Authors:** Carlos Barajas, Domitilla Del Vecchio

**Affiliations:** Department of Mechanical Engineering, Massachusetts Institute of Technology, Cambridge, Massachusetts

## Abstract

Intracellular spatial heterogeneity is frequently observed in bacteria, where the chromosome occupies part of the cell’s volume and a circuit’s DNA often localizes within the cell. How this heterogeneity affects core processes and genetic circuits is still poorly understood. In fact, commonly used ordinary differential equation (ODE) models of genetic circuits assume a well-mixed ensemble of molecules and, as such, do not capture spatial aspects. Reaction-diffusion partial differential equation (PDE) models have been only occasionally used since they are difficult to integrate and do not provide mechanistic understanding of the effects of spatial heterogeneity. In this paper, we derive a reduced ODE model that captures spatial effects, yet has the same dimension as commonly used well-mixed models. In particular, the only difference with respect to a well-mixed ODE model is that the association rate constant of binding reactions is multiplied by a coefficient, which we refer to as the binding correction factor (BCF). The BCF depends on the size of interacting molecules and on their location when fixed in space and it is equal to unity in a well-mixed ODE model. The BCF can be used to investigate how spatial heterogeneity affects the behavior of core processes and genetic circuits. Specifically, our reduced model indicates that transcription and its regulation are more effective for genes located at the cell poles than for genes located on the chromosome. The extent of these effects depends on the value of the BCF, which we found to be close to unity. For translation, the value of the BCF is always greater than unity, it increases with mRNA size, and, with biologically relevant parameters, is substantially larger than unity. Our model has broad validity, has the same dimension as a well-mixed model, yet it incorporates spatial heterogeneity. This simple-to-use model can be used to both analyze and design genetic circuits while accounting for spatial intracellular effects.

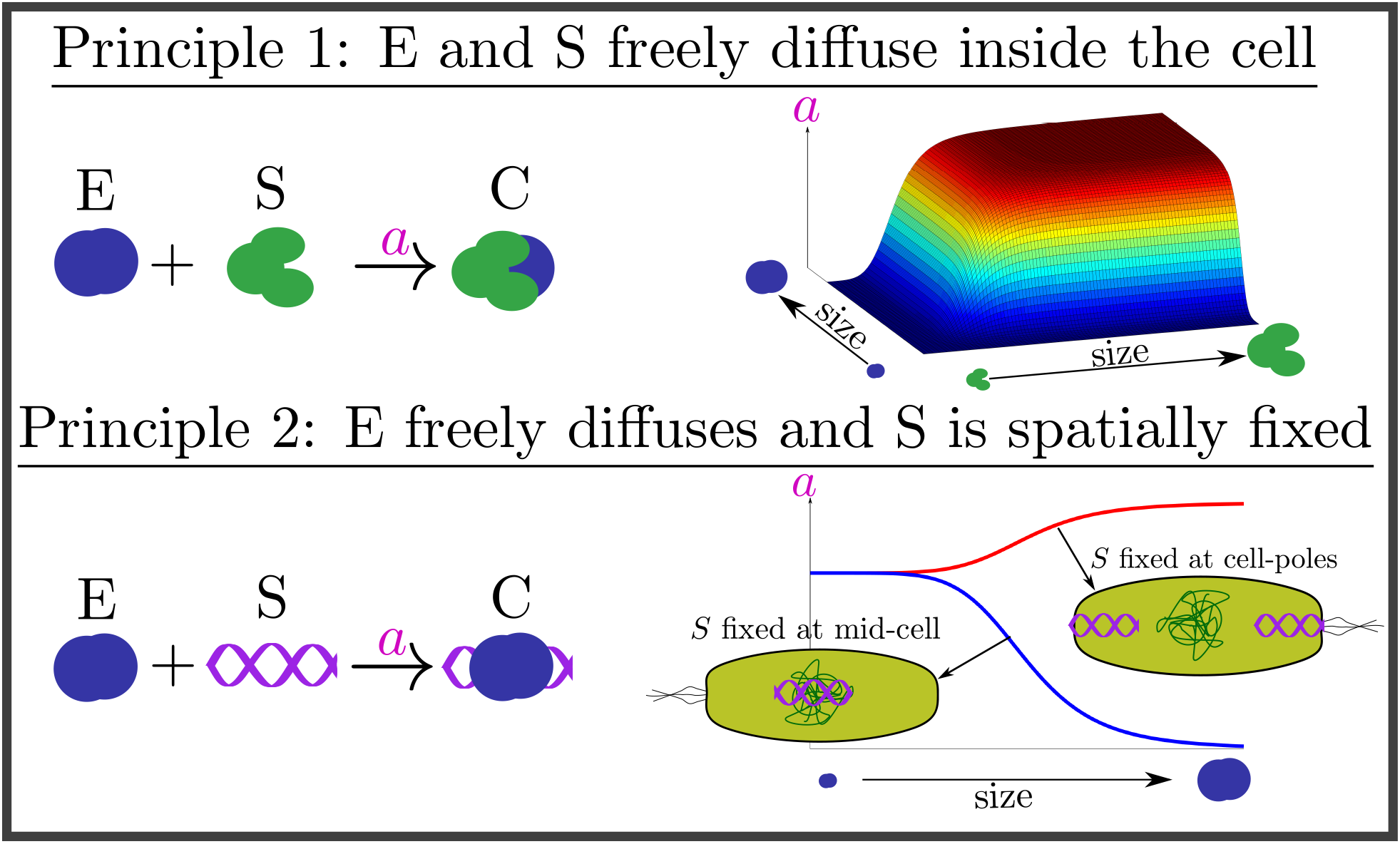

**Highlights:**

- Intracellular spatial heterogeneity modulates the effective association rate constant of binding reactions through a *binding correction factor* (BCF) that fully captures spatial effects
- The BCF depends on molecules size and location (if fixed) and can be determined experimentally
- Spatial heterogeneity may be detrimental or exploited for genetic circuit design
- Traditional well-mixed models can be appropriate despite spatial heterogeneity

**Statement of significance:** A general and simple modeling framework to determine how spatial heterogeneity modulates the dynamics of gene networks is currently lacking. To this end, this work provides a simple-to-use ordinary differential equation (ODE) model that can be used to both analyze and design genetic circuits while accounting for spatial intracellular effects. We apply our model to several core biological processes and determine that transcription and its regulation are more effective for genes located at the cell poles than for genes located on the chromosome and this difference increases with regulator size. For translation, we predict the effective binding between ribosomes and mRNA is higher than that predicted by a well-mixed model, and it increases with mRNA size. We provide examples where spatial effects are significant and should be considered but also where a traditional well-mixed model suffices despite severe spatial heterogeneity. Finally, we illustrate how the operation of well-known genetic circuits is impacted by spatial effects.

## 1 Introduction

Deterministic models of gene circuits typically assume a well-mixed ensemble of species inside the cell [1, 2]. This assumption allows one to describe genetic circuit dynamics through a set of ODEs, for which a number of established analysis tools are available [1]. However, it is well known that spatial heterogeneity is prevalent inside bacterial cells [3, 4, 5, 6, 7, 8]. Depending on the origin of replication, plasmids tend to localize within bacterial cells [9, 10, 11]. Furthermore, chromosome genes (endogenous and synthetically integrated ones [12]) are distributed in the cell according to the chromosome complex spatial structure. In bacterial cells, any molecule freely diffusing through the chromosome (e.g., mRNA, ribosome, and protease) experiences what are known as *excluded volume effects* which capture the tendency of species to be ejected from the nucleoid due to the space occupied by the dense DNA mesh [13]. These excluded volume effects for ribosomes and RNAP in bacteria have been observed experimentally [14].

Despite the strong evidence in support of spatial heterogeneity within bacterial cells, a convenient modeling framework that captures the spatio-temporal organization of molecules inside the cell is largely lacking. As a consequence, how spatial effects modulate genetic circuit dynamics remains also poorly understood. Partial differential equation (PDE) models have been employed on an *ad hoc* basis to numerically capture intracellular spatial dynamics for specific case studies [15, 16, 17]. Although a general PDE model of a gene regulatory network (GRN) can be constructed, it is difficult to analyze and impractical for design [18]. Recently, the method of matched asymptotic expansions was used to simplify the PDEs to a set of ODEs to analyze ribosome-mRNA interactions [19]. Similarly, [20] used a compartmentalized model to capture spatial heterogeneity in sRNA-mRNA interactions. However, these results have not been generalized, relied on simulation, and specific parameter values.

In this paper, we provide a general framework to model spatial heterogeneity through an ODE that has the same structure and hence dimensionality as a well-mixed ODE model. To this end, we first introduce a PDE model that captures spatial dynamics. Next, we exploit the time scale separation between molecule diffusion and biochemical reactions to derive a reduced order ODE model of the space averaged dynamics. This model accounts for spatial heterogeneity by multiplying the association rate constant of binding reactions by a factor that depends on the size of freely diffusing species and on the location of spatially fixed species. We call this factor the *binding correction factor* (BCF). Thus, this reduced model has the same dimensionality as traditional well-mixed models, yet it captures spatial effects.

We demonstrate the effects of spatial heterogeneity in genetic circuit behavior by modeling and analyzing several core biological processes. We show that the transcription rate of a gene and the affinity at which transcription factors bind to it, is lower (higher) when the gene is located near mid-cell (cell poles) with respect to the well-mixed model. We show that compared to a well-mixed model, translation rate is always higher and increases with mRNA size. Finally, we consider a genetic clock, a circuit that produces sustained oscillations. We show that for a parameter range where a well-mixed model predicts sustained oscillations, a model that accounts for spatial heterogeneity of DNA may not show oscillations. All of these phenomena can be recapitulated by our reduced ODE model.

## Methods

We use mathematical models to investigate the effects of spatial heterogeneity, specifically DNA localization and excluded volume effects, on genetic circuit behavior. The first part of this section introduces the mathematical model used, a set of nonlinear PDEs. Model reduction is performed on the resulting PDEs to obtain the reduced ODE model that we use to predict how molecule size and location affect genetic circuit’s behavior.

### 1.1 Reaction-Diffusion Model

A reaction-diffusion model describes the concentration of a species at a given time and location in the cell. We focus on enzymatic-like reactions since they can be used to capture most core processes in the cell. We specialize the model to the cases where the reacting species both freely diffuse or where one freely diffuses while the other one is fixed. For example, mRNA and ribosomes are both freely diffusing, while for RNA polymerase and DNA, one is freely diffusing and the other one is fixed.

#### 1.1.1 Enzymatic-like reactions that model core biological processes

Let S be a substrate being shared by *n* enzymes E_i_, to form product P_i_ where *i* = 1, …, *n*. The rate at which E_i_ and S are produced is given by *α_i_* and *α_s_* respectively. The decay rates of E_i_ and S are given by *γ_i_* and *γ_s_* respectively. Here, we assume that E_i_ and S can be degraded even in complex form, that is, the complex is not protecting them from degradation. Finally, all species are diluted as the cell divides at a rate *μ*. The biochemical reactions corresponding to this process are given by:

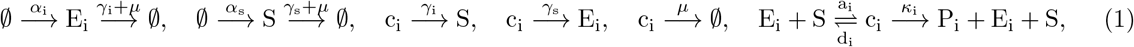

where c_i_ is the complex formed when E_i_ binds to S, *a_i_* is the association rate constant, *d_i_* is the dissociation rate constant, and *κ_i_* is the catalytic rate constant of product formation. These enzymatic-like reactions capture many core biological processes such as genes transcribed by RNA polymerase, mRNA translated by ribosomes, or proteins degraded by a common protease [1]. Notice that they differ from the classical enzymatic reactions since the substrate is not converted into product [1].

*E. coli* actively regulates its geometry to achieve a near-perfect cylindrical shape [21]. Thus, we model the cell as a cylinder of length 2*L* and radius *R_c_* This geometry is shown in Figure 1-A. We assume angular and radial homogeneity ((*R_c_/L*)^2^ ≪ 1) such that the concentration of a species varies only axially (the spatial *x* direction). Symmetry relative to mid-cell is assumed and hence only half of the cell is considered, that is, *x* ∈ [0, *L*], where *x* = 0 is at mid-cell and *x* = *L* is at the cell poles. Furthermore, we assume a constant cross-sectional area along the axial direction.

**Figure 1:**
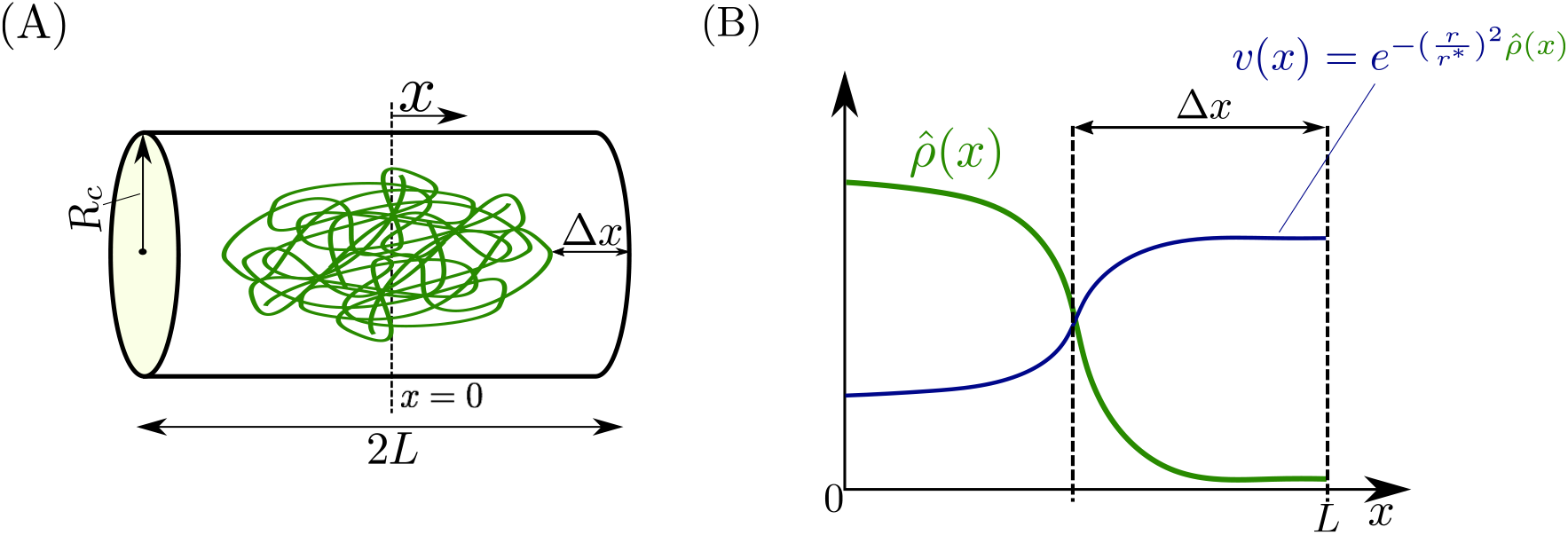
Intracellular spatial geometry. (A) We model the cell as a cylinder of radius *R_c_* and length 2*L* The distance between the end of the chromosome and the cell poles is Δ*x*. (B) The spatial profiles for the normalized local density of DNA length 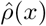 and the fraction of available volume *v*(*x*) of a freely diffusing species with radius of gyration *r* within the chromosomal mesh. These two quantities are related by 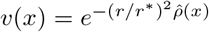, where *r*^*^ is a length scale dependent on the averaged chromosome density in the cell given by (2). The chromosome density is assumed to be monotonically decreasing from mid-cell to the cell poles (as in [13]), thus the available volume profile are monotonically increasing.

In [14] it was shown that polysomes were excluded from the dense chromosomal DNA mesh onto the cell poles. These phenomena is generalized for any species that freely diffuses within the DNA mesh and is referred to as “excluded volume effects”. Leveraging the diffusion modeling framework from [13], we now specify the model to capture excluded volume effects. Let *v*(*x*) ∈ (0, 1] be the volume fraction (dimensionless) available to a species to diffuse within the chromosome (Figure 1-B). As derived in [13] and discussed in SI Section 2.5, the available volume profile *v*(*x*) of a species with a radius of gyration *r*, takes the form

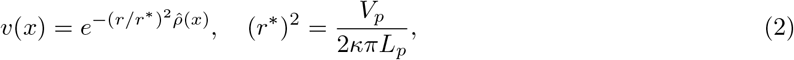

where 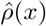 is the normalized local density of chromosome DNA length such that 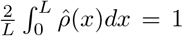, *L_p_* is the total length of chromosome DNA, *V_p_* the volume where the DNA polymer is confined, such that *L_p_*/*V_p_* is the total DNA length per volume and *κ* is an empirically determined correction factor (see [13] and SI Section 2.5). The quantity (*r*^*^)^2^ is inversely proportional to the total DNA length per volume. The local DNA density 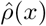 is assumed to be monotonically decreasing (i.e, the chromosome is more dense near mid-cell than at the cell poles as shown in Figure 1-B). Therefore by (2), the available volume profile is higher near mid-cell than at the cell poles (i.e., *v*(0) < *v*(*L*)) as shown in Figure 1-B and furthermore, the discrepancy between *v*(0) and *v*(*L*) increases with *r/r*^*^. For all simulations in this study, we model the normalized chromosome density as

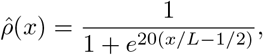

as experimentally determined in [13]. We note that the specific expressions of 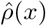 and *r/r*^*^ do not affect the model reduction result of this paper. The main results in this paper are presented for a constant cell length *L* and chromosome DNA density 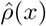, however in SI Section 2.13 we relax these assumptions and allow these quantities to vary in time as the cell divides.

For any given species with concentration per unit length given by *y*(*t, x*), free to diffuse, with available volume *v*(*x*), an expression for the flux term, derived in [13] is given by:

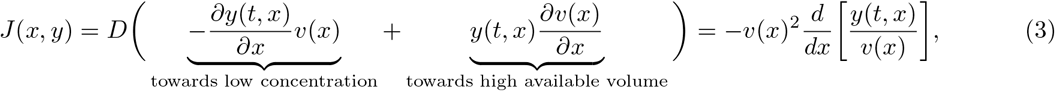

where *D* is the diffusion coefficient. The flux is driven by two mechanisms: the first is concentration gradient, which pushes molecules from high to low concentrations and the second drives molecules to regions with a higher volume fraction. This second term is referred to as the excluded volume effect [13]. From (3), if 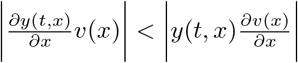 and 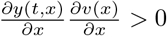, then the net flux is from low to high concentration, is the case when species are repelled from the which to high concentration areas in the cell poles. As we will show, this mechanism dictates intracellular heterogeneity in the limit of fast diffusion.

For species S, we denote by *S*(*t, x*) its concentration per unit length at time *t* at location *x* (similarly for E_i_ and c_i_). Assuming sufficently high molecular counts, the reaction-diffusion dynamics corresponding to (1) describing the rate of change of the species concentrations at position *x* are given by [22] :

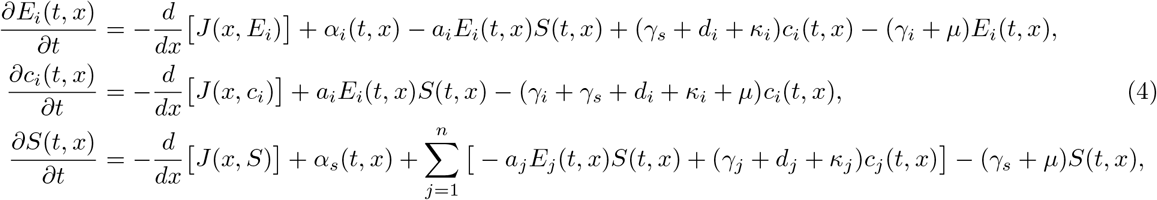

where *J* (*x*, ·) is the flux per unit area per unit time, within the cell. If the species is freely diffusing *J* (*x*, ·) is given by (3), otherwise if the species is spatially fixed, then *J* (*x*, ·) = 0 for all *x* ∈ [0, *L*]. The boundary conditions associated with freely diffusing species of (4) are zero flux at the cell poles and cell center due to the assumed left-right symmetry, which corresponds to:

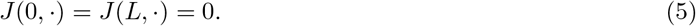

Notice that none of the parameters in (4) appearing in (1) depend explicitly on time and space except for the production terms *α_i_*(*t, x*) and *α_s_*(*t, x*). The explicit time dependence of the production terms allows us to capture how genes can be activated or repressed externally with a time varying signal [23]. The explicit dependence of the production terms on *x* allows us to capture where the species is produced within the cell (e.g., DNA in the chromosome or DNA in pole localized plasmid genes).

##### Dimensionless model

Depending on the parameter regimes, the dynamics of (4) can display time scale separation. For example, diffusion occurs in the order of mili-seconds compared to minutes for dilution due to cell-growth and mRNA degradation [2]. Therefore, we are interested in determining the behavior of (4) in the limit of fast diffusion. We thus rewrite (4) in dimensionless form to make time scale separation explicit. We nondimensionalize the system variables using dilution (1/*μ*) as the characteristic time scale, the length of the cell (*L*) as the characteristic length, and *μ/a*_1_ as the characteristic concentration per length scale: 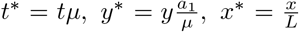, where *y* denotes concentration per unit length and the superscript “*” is used on the dimensionless variable. Concentrations are nondimensionalized through *a*_1_ because this parameter contains a concentration scale, it is fixed in time, and it is assumed to be nonzero. The dimensionless form of (4) is given by

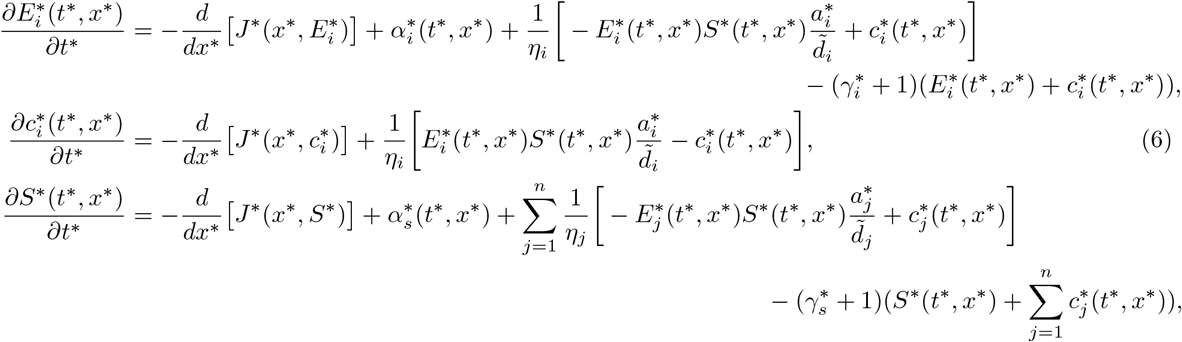

where 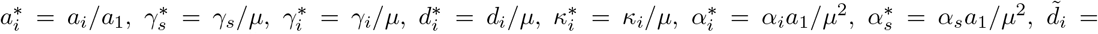 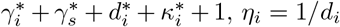, and *J*^*^ = *Ja*_1_(*μ*^2^*L*). For a freely diffusing species with diffusion coefficient *D*, the dimensionless parameter that determines the relative speed of diffusion is denoted by *ϵ* = *μL*^2^*/D* and fast diffusion corresponds to *ϵ* ≪ 1. Likewise, *η_i_* in (4) determines the relative speed of the binding dynamics, where *η_i_* ≪ 1 implies these reactions are fast. From hereon, unless otherwise specified, we work with variables in their dimensionless form and drop the star superscript for simplifying notation.

##### Space averaged concentrations

Concentrations per cell are usually the quantities measured experimentally [24] and are the primary quantities of interest. We now derive the space averaged dynamics corresponding to (6), which describe the dynamics of concentrations per half of the cell. We define 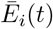, 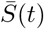, and 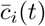 to be the *space averaged enzyme, substrate, and complex concentrations* respectively, and are given by

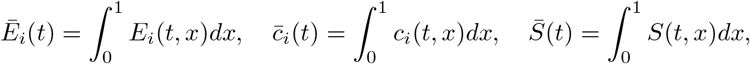

also giving the concentrations per half of the cell. The dynamics governing these space averaged variables are derived by integrating (6) in space and applying the boundary conditions (5) and are given by:

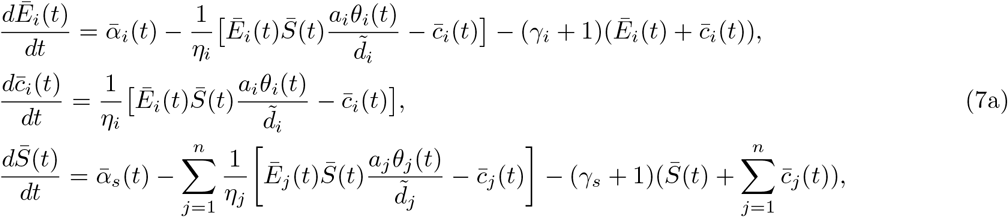

where overbars denote spatially averaged variables and

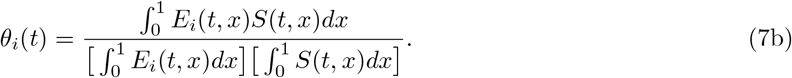

Therefore, to calculate the space averaged concentrations, one could integrate the outputs of the full PDE (6) directly or use (7) along with (7b), as illustrated in Figure 2. Notice that calculating *θ_i_* in (7b) requires solving the full PDE system (6) because of its dependence on the product *E_i_*(*t, x*)*S*(*t, x*). Therefore, in general, there is no obvious benefit in working with (7). In this paper, we provide a method to compute a guaranteed approximation of *θ_i_* without solving the PDEs (6).

**Figure 2:**
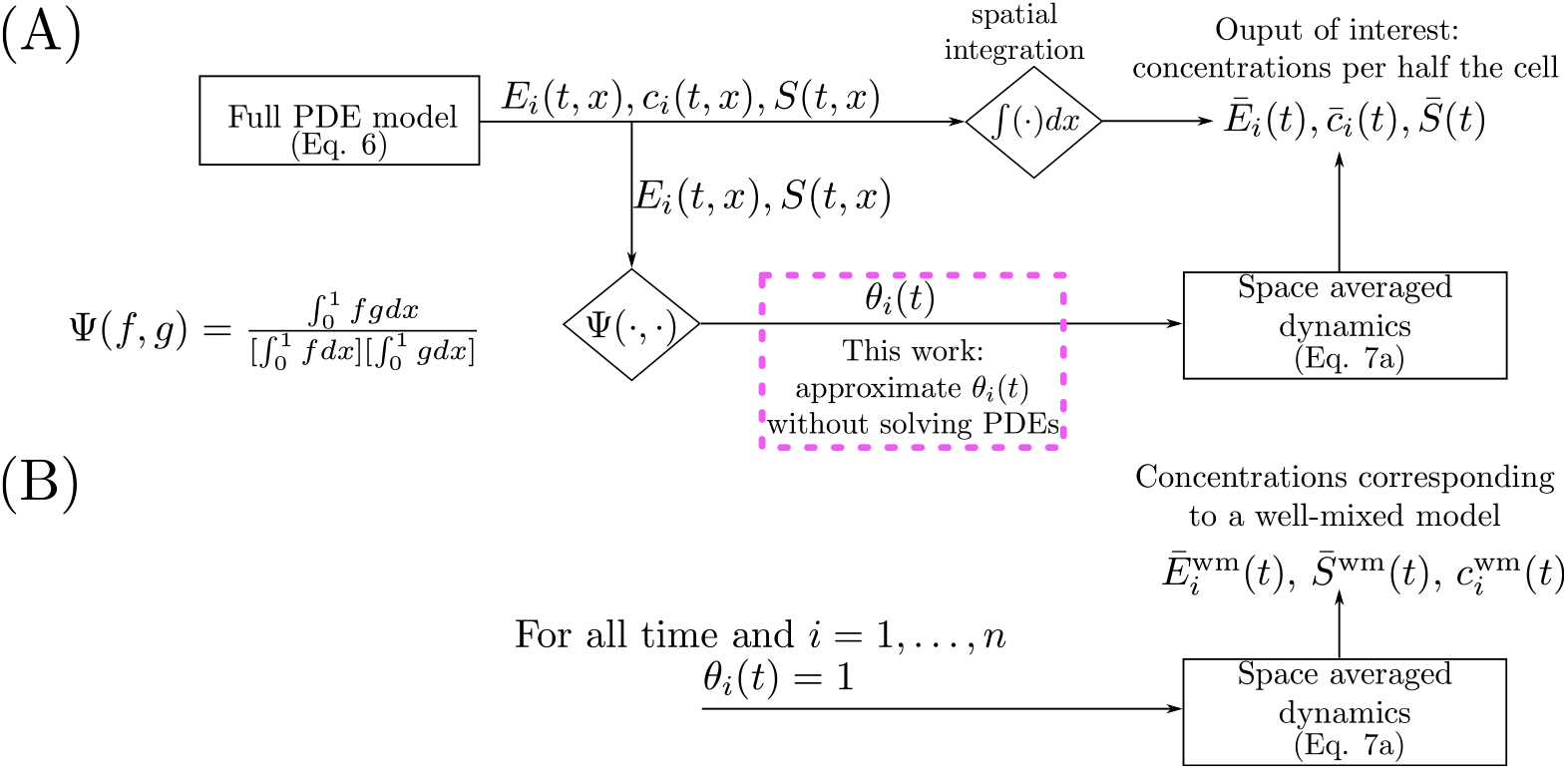
Methods to calculate space averaged concentrations. (A) The time and space dependent solutions of the full PDE (6) are integrated spatially to yield concentrations per half of the cell. Alternatively, space averaged concentrations can be calculated using the space average dynamics (7) with the BCF (*θ_i_*(*t*) (7b)) as a time varying parameter. (B) The dynamics of the space averaged concentration is given by the well mixed model (8) when *θ_i_*(*t*) = 1 for all time and for all *i* = 1, …, *n*.

##### Well-mixed model

Next, we define what we have been informally referring to as the “well-mixed” model [25]. A standard well-mixed model is derived starting from (1) assuming mass action kinetics, that molecular counts are sufficiently large, and that the intracellular environment is spatially homogeneous (well-mixed) [1]. We let 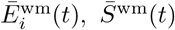 and 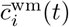, denote the well-mixed concentrations of E_i_, S, and c_i_, respectively, and their dynamics are given by

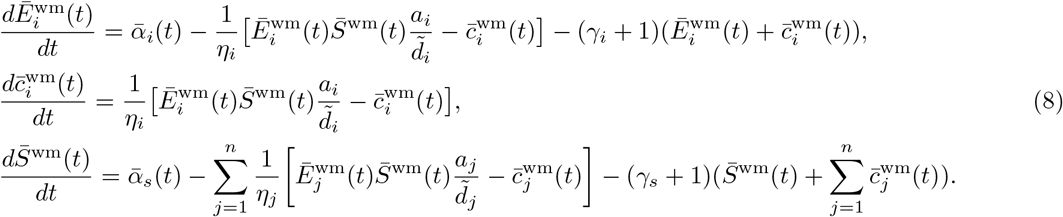

Comparing (7) and (8), motivates us to define 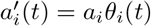, which can be regarded as the effective association rate constant between E_i_ and S in (7). We refer to *θ_i_*(*t*) as the *binding correction factor* (BCF). The dynamics of the space averaged concentrations (7) coincide with those of the well-mixed model (8) when *θ_i_*(*t*) = 1 (thus 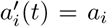) for all time and for all *i* = 1, …, *n*. From (7b) notice that *E_i_*(*t, x*) and *S*(*t, x*) being spatially constant for all time is not necessary for *θ_i_*(*t*) = 1 for all time. For example, if *S*(*t, x*) is spatially constant while *E_i_*(*t, x*) has an arbitrary spatial profile (or *vice-versa*), then *θ_i_*(*t*) = 1. Thus, the space averaged concentrations can coincide with those of a well-mixed model despite severe spatial heterogeneity. In this work, we provide a constant approximation of *θ_i_*(*t*) denoted by 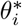, which depends on spatial variables such as molecule size and gene location. Under the fast diffusion approximation, we show that 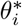 is close to *θ_i_*(*t*). The space averaged dynamics (7) with *θ_i_*(*t*) replaced by 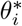, thus provides a reduced ODE model that captures spatial information without having to solve (6). We will compare how solutions to (7) with (7b) calculated from the full PDE (6) or with 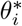 compare to each other and to the solutions of the well-mixed model (8).

#### 1.1.2 Three diffusion cases to capture core biological processes

To use model (6) to describe key biological processes, we consider three cases. In Case I, E_i_ for all *i* = 1, …, *n* and S are all freely diffusing within the cell. In Case II, E_i_ is spatially fixed for all *i* = 1, …, *n* (*J*(*x, E_i_*) = 0 for all *x* ∈ [0, 1] and for all *i* = 1, …, *n*) and S is freely diffusing. In Case III, E_i_ is freely diffusing for all *i* = 1, …, *n* and S is spatially fixed (*J*(*x, S*) = 0 for all *x* ∈ [0, 1]). Case I may represent mRNA molecules (E_i_) competing for ribosomes (S), all freely diffusing in the cell. Case II captures genes (E_i_), which are spatially fixed and are transcribed by RNA polymerase (S), which freely diffuses. Case III models transcription factors (E_i_), which freely diffuse regulating the same spatially fixed gene (S).

The flux dynamics, the boundary conditions, and a core biological process example for each case are summarized in Table. 1. When a species is spatially fixed, the flux is zero through the whole domain, that is, *J*(*x*, ·) = 0 for all *x* ∈ [0, 1]. The available volume profiles for the enzyme, complex, and substrate are denoted by 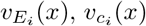, and *v_S_*(*x*), respectively. The available volume profile for the complex 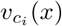, represents the probability that the complex has enough free volume to hop into the DNA mesh at position *x* and it equals the product of the probability of the two independent events of the enzyme and the substrate hopping into the DNA mesh [13], thus

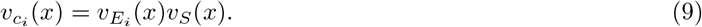

**Table 1:** The flux dynamics and the boundary conditions corresponding to (6) for each case of interest along with a core process example. Here *v_E,i_*(*x*), *v_S_*(*x*), and *v_c,i_*(*x*), are the available volume profiles of E_i_, S, and c_i_, respectively. The parameters 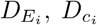, and *D_s_* are the enzyme, complex, and substrate diffusion coefficients, respectively, *ϵ* is a dimensionless parameter that captures the speed of diffusion (with respect to dilution). A species being spatially fixed translates to the flux being zero throughout the whole spatial domain. In Case II, for *i* = 1, …, *n*, 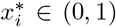 denotes the location of the fixed species E_i_. In Case III, 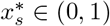 denotes the location of the fixed species S.

|  | **Case I**<br>All species diffuse | **Case II**<br>Substrate diffuse<br>and enzymes fixed | **Case III**<br>Enzymes diffuse<br>and substrate fixed |
| --- | --- | --- | --- |
| **Dimensionless Flux** | $J(x, E_i) = -\frac{1}{\epsilon} \chi_{E_i} v_{E_i}^2 \frac{d}{dx} \left[ \frac{E_i}{v_{E_i}} \right]$<br>$J(x, S) = -\frac{1}{\epsilon} v_S^2 \frac{d}{dx} \left[ \frac{S}{v_S} \right]$<br>$J(x, c_i) = -\frac{1}{\epsilon} \chi_{c_i} v_{c_i}^2 \frac{d}{dx} \left[ \frac{c_i}{v_{c_i}} \right]$ | $J(x, E_i) = 0$<br>$J(x, S) = -\frac{1}{\epsilon} v_S^2 \frac{d}{dx} \left[ \frac{S}{v_S} \right]$<br>$J(x, c_i) = 0$ | $J(x, E_i) = -\frac{1}{\epsilon} v_{E_i}^2 \frac{d}{dx} \left[ \frac{E_i}{v_{E_i}} \right]$<br>$J(x, S) = 0$<br>$J(x, c_i) = 0$ |
| **Boundary conditions** | $J(0, E_i) = J(1, E_i) = 0$<br>$J(0, c_i) = J(1, c_i) = 0$<br>$J(0, S) = J(1, S) = 0$ | $J(0, S) = J(1, S) = 0$ | $J(0, E_i) = J(1, E_i) = 0$ |
| $\epsilon$ | $(\mu L^2)/D_s$ | $(\mu L^2)/D_s$ | $(\mu L^2)/D_{E_1}$ |
| **Dimensionless diffusion** | $\chi_{E_i} = D_{E_i}/D_s, \chi_{c_i} = D_{c_i}/D_s$ | N/A | N/A |
| **Core process** | mRNAs binding<br>ribosomes | RNAP binding<br>several genes | Transcription factors<br>binding promoter |
| **Location of fixed species** | N/A | $x_i^*$ | $x_s^*$ |

Furthermore, we define the normalized available volume profiles as

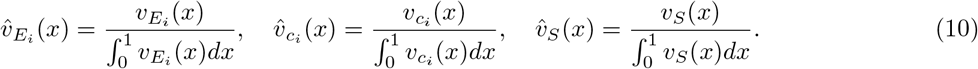

### 1.2 Numerical Method: Finite Difference

In general, a closed form solution to (6) is not available. Therefore, we rely on numerical solutions to directly integrate the PDE. In particular, we utilize a finite difference method that is widely used to simulate PDEs [26]. For a general diffusing species with its concentration given by *y*(*t, x*), and available volume profile *v*(*x*), we make the following coordinate transformation *u*(*t, x*) ≔ *y*(*t, x*)/*v*(*x*). In this coordinate, the boundary conditions (if applicable) are Neumann [27]: 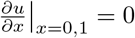 and thus are simpler to implement. Furthermore, with this transformation we observe less stiffness in the numerical simulations. We discretized the spatial domain into *N* + 1 equidistant points such that Δ = 1/*N*. Using a second order finite difference method, we approximate the derivatives as:

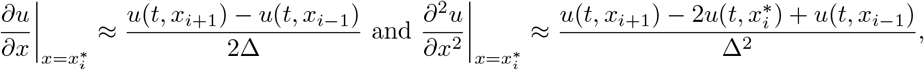

that appear in the flux terms (in the case where species freely diffuses). The new boundary conditions give rise to the following constraints:

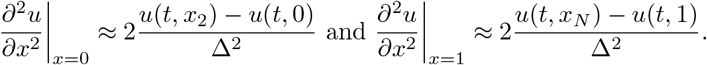

This discretization leads to a system of *N* + 1 ODEs. The resulting set of ODEs are then simulated with MATLAB, using the numerical ODE solvers ode23s. To calculate the space averaged concentrations, we implement a second oder trapezoidal integration scheme [27] given by:

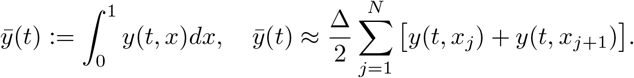

The convergence rate of our numerical method is demonstrated in Section 2.10. For all simulations in this paper *N* = 200.

## Results

### 1.3 Time Scale Separation

In this section, we provide a time independent approximation of the BCF (7b) in the limit of fast diffusion, which depends solely on the size of diffusing species, chromosome density profiles, and the spatial localization of non-diffusing species. With this approximation, we can compute space averaged solutions in (7) without solving the PDEs in (6).

#### 1.3.1 Reduced space averaged dynamics when diffusion is fast and fixed species are localized

For Case II and Case III of Table 1, in which one of the reacting species is fixed, we assume that the *fixed species is spatially localized* to a small space, that is, we have the situation depicted in Figure 3 (see SI Section 2.3, Assumption 3 for the mathematical definition). Practically, for Case II, spatial localization at 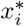 requires that the production rate *α_i_*(*t, x*) of the fixed species is smaller than some small threshold *δ* when *x* is outside the interval 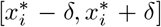 for all time and that the space averaged production rate is 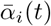 independent of *δ* (similarly for Case III, 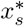, and *α_s_*(*t, x*)). From a biological perspective, having the space averaged production rate independent of *δ* is consistent with the fact that the total amount of DNA in the cell is independent of where the DNA is concentrated. Note that *δ* is a parameter that controls the amount of localization, such that *δ* ≪ 1 implies the production of spatially fixed species being localized to a small region. Let *ϵ* be as in Table 1 that appears in (6), the following definition will provide the candidate reduced model that approximates (7) well when *ϵ* ≪ 1 and *δ* ≪ 1. Recall that *ϵ* is a dimensionless parameter that captures the speed of diffusion (with respect to dilution).

**Figure 3:**
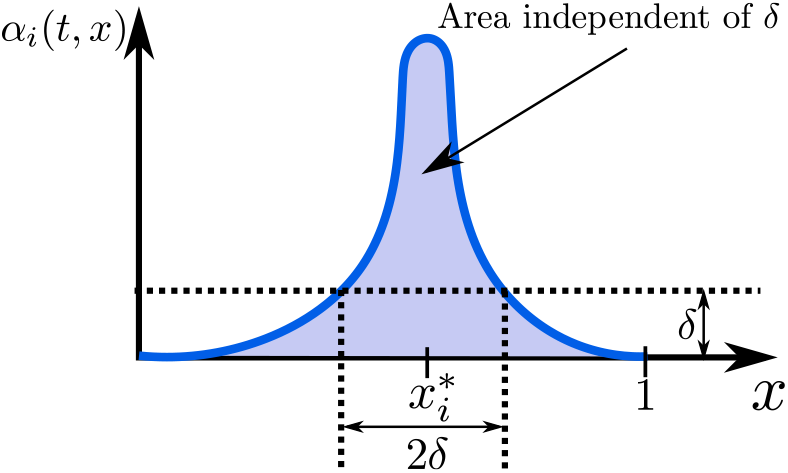
Graphical representation of localization of fixed species. The production rate *α_i_*(*t, x*) is assumed to be localized at 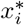 if 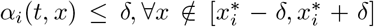. We assume that the space averaged production 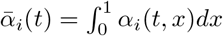 is independent of *δ*.

Let 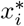 and 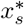 be the location of the fixed species for Case II and Case III, respectively (see Table 1). For *i* = 1, …, *n*, we define the *reduced space-averaged dynamics* as

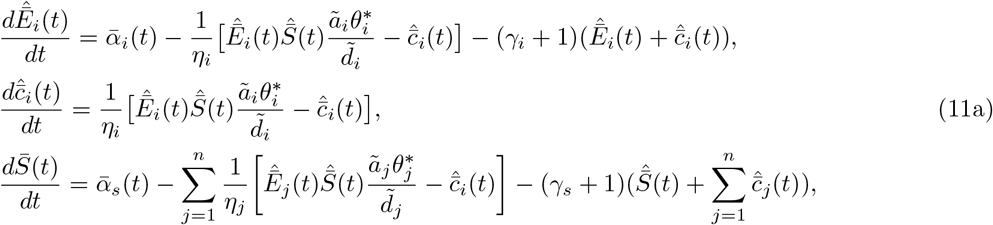

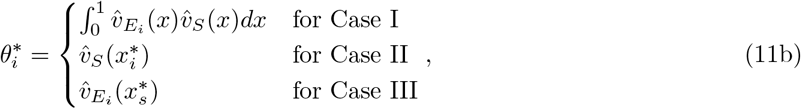

where 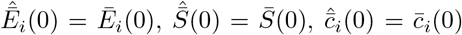, as given by (7), and 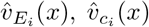, and 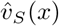 are given by (10). Then, we have the following main result of this paper (see SI Section 2.3, Theorem 3 for a formal statement with the proof).

##### Result 1

*Consider system* (6) *and let z*(*t, x*) = [*E*_1_(*t, x*),…, *E_n_*(*t, x*), *c*_1_(*t, x*), …, *c_n_*(*t, x*), *S*(*t, x*)]^*T*^ with 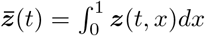. *Consider system* (11) *and let* 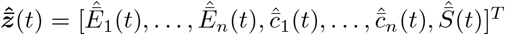. *Then, for all t* ≥ 0 *and ϵ, δ sufficiently small*

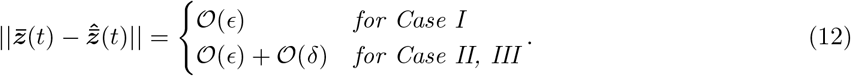

By virtue of this result, we can use the simple and convenient ODE model in equations (11) to describe the space-averaged dynamics of the PDE system (6). In particular, from (11) it appears that spatial effects are lumped into the BCF approximation 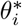. Therefore, in order to determine how spatial heterogeneity affects system dynamics, it is sufficient to analyze how dynamics is affected by parameter 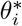 and how the expression of 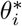 is, in turn, affected by spatial localization and molecule size (see (11b) and (2)).

##### Remark 1

As discussed in SI Section 2.3, as *ϵ* → 0^+^, the spatial profile of diffusing molecules approaches that of their available volume profile, that is,

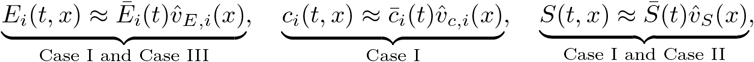

for the other spatially fixed species we have that their concentrations are localized in a manner as their production terms.

The consequence of Remark 1 is that knowledge of the space averaged dynamics from system (11) also leads to knowledge of the spacial profiles of the species within the cell. This information is used to propose a method to estimate the BCF from experimental data (See SI Section 2.11)

##### Remark 2

The approximation result holds for *ϵ* ≪ 1, that is, diffusion is much faster than any other time scales in (6). However, in SI Section 2.4 we motivate why the approximation should still hold (for which the relationship (9) is key) if 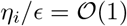 (binding and unbinding between E_i_ and S occurs at a similar timescale as diffusion), and confirmed via numerical simulations in Section 1.4.

The BCF 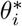 in (11b) is temporally constant and thus the reduced model has the same dimensionality as the well-mixed model (8), yet captures the role of spatial heterogeneity in the interactions between cellular species. Therefore, 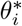 is a practical and accurate approximation of the BCF when *ϵ* ≪ 1 (sufficient for Case I) and *δ* ≪ 1 (needed for Cases II-III).

#### 1.3.2 Dependence of the BCF on species size and localization

When diffusion is fast and the expression of spatially fixed species is localized, the BCF is well approximated by 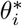 given in (11b). Substituting (2) into (11b) and denoting the radius of gyration of E_i_ and S by *r_e,i_* and *r_s_* respectively, we can rewrite 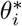 as

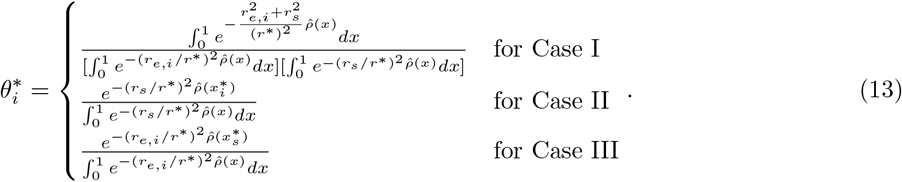

From (13), we observe that 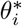, depends on the spatial localization of spatially fixed species (i.e,. 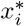 and 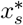), the radius of gyration of diffusing species, *r*^*^ (2), and the nominalized local density of DNA length 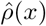.

Using (13), we graphically illustrate the dependence of 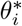 on 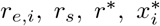 and 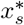 in Figure 4. By analyzing Figure 4, we observe the following:

**Figure 4:**
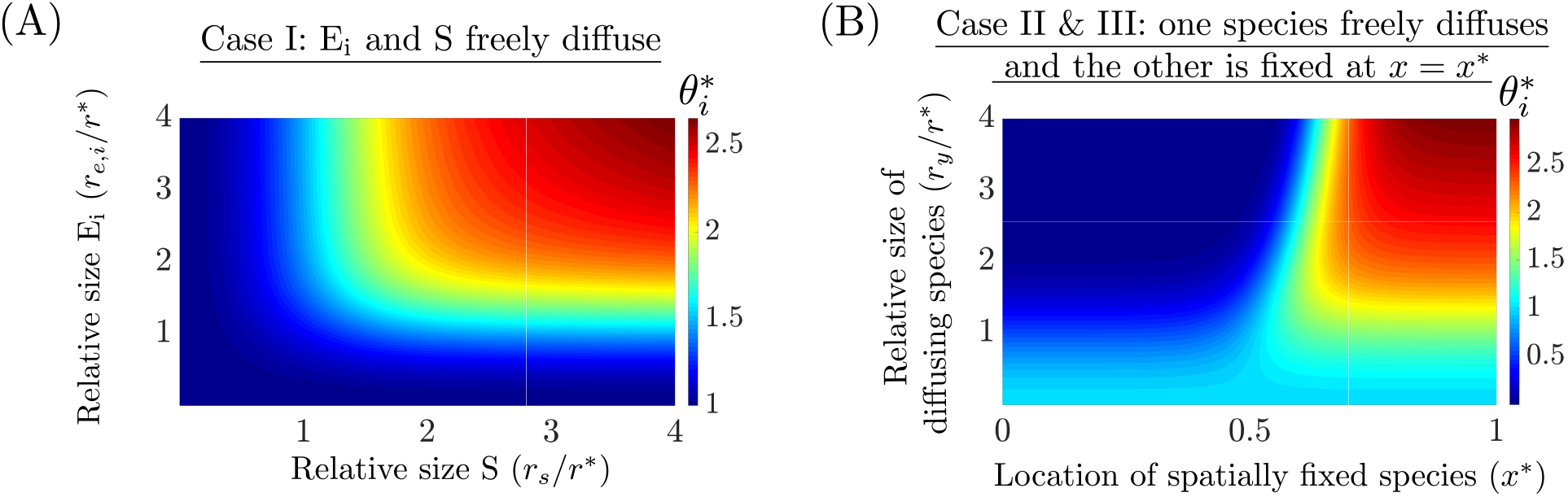
The BCF in the limit of fast diffusion and localization of spatially fixed species. Approximation of the BCF denoted by 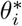 (13) is provided for Case I and for Case II/Case III. (A) For Case I, where E_i_ and S both freely diffuse, *θ_i_* ≥ 1 and increases when both the size of E_i_ (*r_e,i_*) and S (*r_s_*) are sufficiently large (with respect to *r*^*^). (B) For Case II and III, where one of the species diffuses (size *r_y_*) and the other is fixed at *x* = *x*\*, 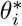 is different from unity when the radii of the diffusion species is sufficiently large. We observe that 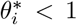 for *x*^*^ ≤ 0.4 and appears to approach zero near *x*^*^ = 0 for large *r_y_/r*^*^. Similarly, for *x*^*^ ≥ 0.65, 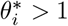. Between 0.4 ≤ *x*^*^ ≤ 0.65 there exists a region that 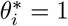 for all *r_y_/r*^*^.

**Case I**: the BCF is always greater than or equal to that of the well-mixed model (8) (where 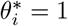 for all *i*) and this discrepancy increases with the size of E_i_ and S. Intuitively, as the size of E_i_ and S increases, they are pushed out of the chromosome and co-localize near the cell poles, thus they are confined to a smaller volume to interact and hence their effective binding strength increases. If only one of the species is large (with respect to *r*^*^), while the other one is small, then the large species will be ejected from the chromosome and thus will not be homogeneously distributed throughout the cell, however 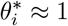, and thus a well-mixed model is valid despite this spatial heterogeneity.

**Case II and III**: where one of the species diffuses (size *r_y_*) and the other is fixed at *x* = *x*^*^, the BFC is different from unity when *r_y_* is sufficiently large. We observe that 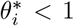 for *x*^*^ ≤ 0.4 and appears to approach zero near *x*^*^ = 0 for large *r_y_/r*^*^. Similarly, for *x*^*^ ≥ 0.65, 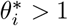. This occurs because as the size of the diffusing species increases, the species is ejected from the chromosome onto the cell poles and therefore it is more likely to interact with species fixed at the cell-poles than those near mid-cell. Between 0.4 ≤ *x*^*^ ≤ 0.65 there exists a region where 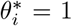 for all *r_y_/r*^*^. This provides additional evidence that a well-mixed model may be appropriate despite severe intracellular heterogeneity.

When 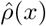 is assumed to be a step function, the upper bound for 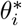 is 1/*Δx* for Case I-III, as derived in SI Section 2.5, where Δ*x* is the distance between the end of the chromosome and the cell poles as shown in Figure 1. Furthermore, the lower bound for Case I was unity and for Case II-III it was zero.

The value of the BCF provides a measure to determine the extent to which spatial effects modulate the biomolecular dynamics. Therefore, an experimental method to estimate the BCF is desirable. In SI Section 2.11, we propose such a method that only requires knowledge of Δ*x* and of the value of concentration of freely diffusing species inside and outside the nucleoid.

In SI Section 2.13, we consider how the BCF can vary temporally as the cell divides and the chromosome density shifts from being concentrated primarily near mid-cell to quarter-cell. We demonstrate that the BCF can vary by over 50% in time for the case where one species is stationary and localized near mid-cell. Furthermore, in SI Section 2.14 we show how the BCF is affected when we consider exclusion effects from the DNA of a pole localized high copy plasmid. We show that for the case where both reactant freely diffuse, the BCF decreases as the amount of plasmid DNA increases. For the case where one reactant is spatially fixed and the other freely diffuses, we show that the BCF decreases for a species localized at the cell poles and increases for a species localized near quarter-cell, as the amount of plasmid DNA increases.

### 1.4 Application to Core Processes and Genetic Circuits

In this section we apply the results of the time scale separation analysis from Section 1.3 in order to both determine and modulate the effects of intracellular heterogeneity on core processes, such as transcription and translation, and on genetic circuit behavior.

#### 1.4.1 Application of the reduced ODE model to transcription and translation

In this section, we investigate how and the extent to which intracellular heterogeneity affects the core biological processes of transcription and translation, which are responsible for protein production. We model a gene (D) being transcribed by RNAP (S) to form a DNA-RNAP complex (c_s_) to produce mRNA (m). The mRNA is then translated by ribosomes (R) to form mRNA-ribosome complex (c_m_) which produces protein P. The chemical reactions are given by

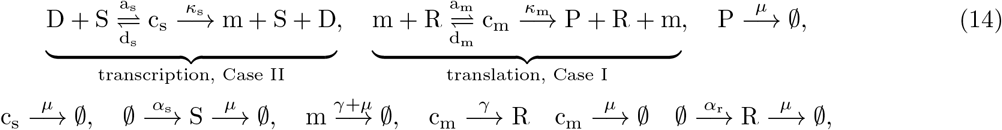

where *a_s_* and *d_s_* are the association and dissociation rate constants, respectively, between RNAP and the gene D, *κ_s_* is the catalytic rate constant of formation of mRNA m, *a_m_* and *d_m_* are the association and dissociation rate constants, respectively, between ribosomes and mRNA, *κ_m_* is the catalytic rate constant of formation of protein P, *α_s_* is the production rate of RNAP, *α_r_* is the ribosome production rate, *μ* is the cell growth rate constant (set to unity in our nondimensionalization), and *γ* is the mRNA degradation rate constant. The transcription reaction is in the form of Case II (Table 1) since the gene does not freely diffuse and the RNAP freely diffuses. The translation process falls under Case I, since both mRNA and ribosomes freely diffuse. We assume that the total concentration of D is conserved, so that *D_T_* (*x*) = *D*(*t, x*) + *c_s_*(*t, x*) and that *D_T_* (*x*) is localized at *x* = *x*^*^. From (11), the dimensionless reduced space averaged dynamics corresponding to (14) are given by

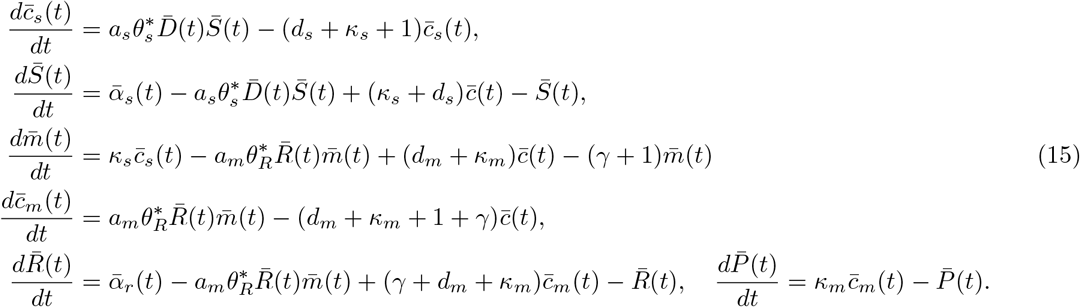

Concentration variables are nondimensionalized with respect to the total steady state space averaged RNAP 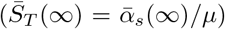, since this quantity is a readily available in the literature. Letting *r_s_*, *r_m_* and *r_R_* be the radius of gyration of RNAP, mRNA, and ribosomes, respectively, we compute the BCF’s via (11b) and (2),

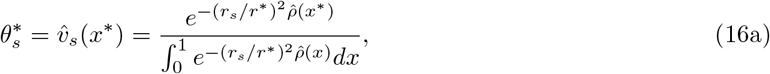

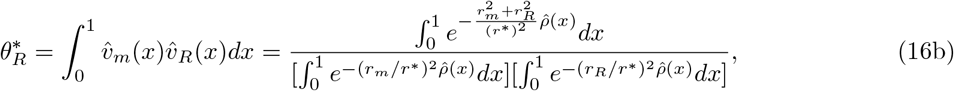

where 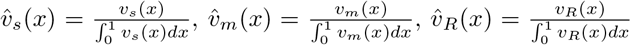, and 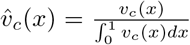 are the normalized available volume profiles of RNAP, mRNA, ribosomes, and of the mRNA-ribosome complex, respectively. Recall that the quantity (*r*^*^)^2^ is inversely proportional to the total DNA length per volume. We now consider the steady state behavior of system (15) by equating the time derivatives to zero. Specifically, we are interested in how the steady state levels of produced mRNA and protein are affected by 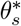 and 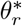 and, hence, how they depend on spatial quantities such as *r_s_/r*^*^ *r_m_/r*^*^, *r_R_/r*^*^, and *x*^*^.

##### Total mRNA steady state level

We are interested in investigating the role of spatial effects on the binding between RNAP and the DNA and thus on mRNA production. Here we analyze the steady state total mRNA levels 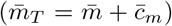 of (15) rather than the free amount of mRNA (*m*), since 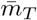 is independent of 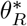 as shown by

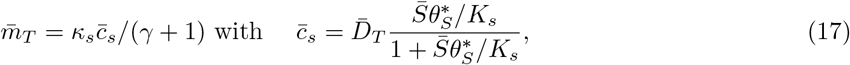

where 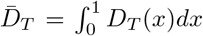, *K_s_* = *d_s_/a_s_*, *K_R_* = *d_m_/a_m_*. If 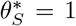 in (17), then the predicted total mRNA steady state level will be identical to that of a well-mixed model (as in (8)). From (16) and Figure 4-B, if the RNAP radius of gyration is sufficiently large with respect to *r*^*^ then 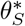 may be different from unity (depending on *x*^*^), in which case spatial effects arise. If the DNA is localized near mid-cell (*x*^*^ ≈ 0), then it implies that 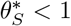 from Figure 4-B and, as a consequence, a decreased steady state total mRNA level will result. Furthermore, for very large values of *r_s_/r*^*^ and *x*^*^ ≈ 0 we have that 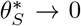 and the total mRNA steady state levels will approach zero. Similarly, if the DNA is localized near the cell-poles (*x*^*^ ≈ 1), then it implies that 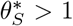 from Figure 4-B and, as a consequence, an increased steady state total mRNA level. This phenomenon occurs because as the excluded volume effects of RNAP are amplified (large *r_s_/r*^*^), RNAP will localize primarily in the cell poles and hence transcribe pole-localized DNA more efficiently than DNA near mid-cell (or any region where the local chromosome density is high). When designing genetic circuits, a plasmid backbone is chosen to provide a certain DNA copy number, however the backbone also determines where in the cell the plasmid localizes [9, 10, 11]. Therefore, based on our results, localization also affects steady state total mRNA level. If instead of introducing the DNA via a plasmid, the DNA is integrated directly into the chromosome, then the location of integration site should be a parameter to consider.

Figure 5-A shows the behavior of the steady state total mRNA level as a function of *r_s_/r*^*^ and of the location of the transcribed gene, when compared to the level predicted by the well mixed model. Simulations confirm that total mRNA levels are higher for pole localized genes than those near mid-cell and that the discrepancy increases with the size of RNAP relative to *r*^*^. The agreement between the full PDE model ((62) in SI Section 2.6) and the reduced ODE model (15) provides numerical validation of the model reduction results (explicitly shown in SI Figure 11). In the SI Section 2.6, we show in Figure 9 the transient response corresponding to Figure 5, for which the full-PDE and reduced models agree. Furthermore, in SI Figure 9, we also verify that as the size of RNAP increases, it is indeed ejected from the chromosome and adopts its available volume profile (Remark 1). Furthermore, in SI-Figure 10, we demonstrate that these results hold independent of the binding and unbinding speed between RNAP and DNA (Remark 2). In SI Section 2.12 we propose an experimental method to test the hypothesis that mid-cell genes are transcribed less effectively than pole localized genes.

**Figure 5:**
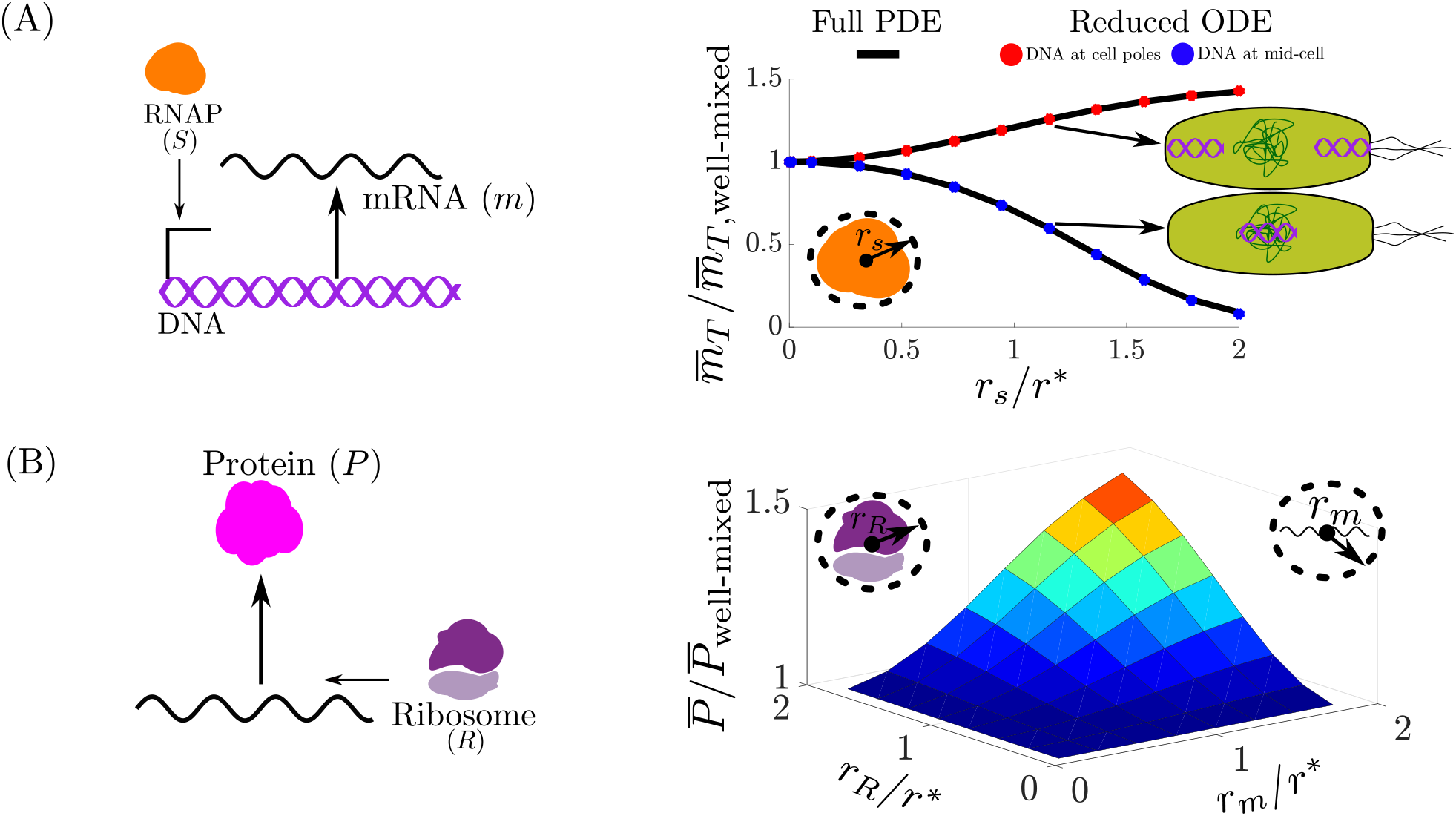
Spatial heterogeneity effects on steady state total mRNA and protein levels. (A) The space averaged total mRNA 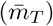 concentration predicted by the full PDE model ((62) in SI Section 2.6) and the reduced ODE model (15) normalized by that of the well-mixed model 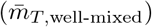 as the size of the RNAP (*r_s_*) varies with respect to *r*^*^. With respect to the well-mixed model, the amount of mRNA decreases (increases) when the DNA is localized near mid-cell (cell poles). (B) The space averaged protein concentration 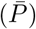 predicted by the full PDE model ((62) in SI Section 2.6) normalized by that of the well-mixed model 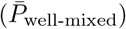 as the size of the of mRNA (*r_m_*) and ribosome (*r_R_*) varies with respect to *r*^*^. The amount of protein increases when both the mRNA and ribosome size increases. We set *r_s_/r*^*^ = 1 × 10^−3^, such that *θ_S_* ≈ 1 and thus the result is independent of the spatial location where the gene is expressed. We refer to the well-mixed model as (15) with (16) given by 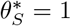 and 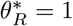. The parameter values and full simulation details are provided in SI Section 2.6

In [28] it was estimated that *r_s_* = 6.5 ± 0.1 nm, which implies that *r_s_/r*^*^ ≈ 0.3. From Figure 4-B, this implies that 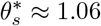 when the DNA is at the cell poles and 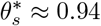 when the DNA is near mid-cell, thus the we expect the binding strength between RNAP and the DNA to deviate by 6% from that of a well-mixed model. From (17), if 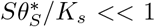, then 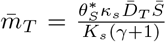; thus in this regime the mRNA concentration is proportional to 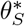. So we expect at most a 6% difference in steady state mRNA concentration with respect to what is predicted by a well-mixed model.

##### Protein steady state level

The steady state protein levels of (15) are given by

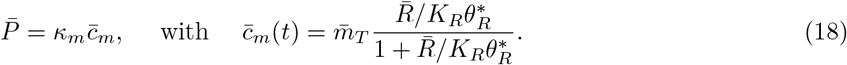

From (17) and (18), if 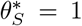 and 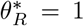, then protein steady state level will be identical to that of a well-mixed model. From (16) and Figure 4-A, we conclude that 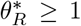 and increases with *r_s_/r*^*^ and *r_m_/r*^*^. Increasing *r_s_/r*^*^ and *r_m_/r*^*^ implies that the ribosomes and mRNA are further excluded from the chromosome onto the cell poles. Once localized at the cell-poles, the mRNA and ribosomes are more likely to bind. Figure 5-B shows the behavior of the steady state protein levels as a function of *r_m_* and *r_R_* when compared to the level predicted by the well mixed-model for the full PDE model ((62) in SI Section 2.6). Simulations confirm that protein levels with respect to a well-mixed model increases when both the mRNA and ribosome size are sufficiently large. In SI-Figure 13, we show that the reduced ODE model (15) is within 2% of the full PDE model (SI (62)) for the result in Figure 5-B. In the SI Section 2.6 we show in Figure 12 the transient response corresponding to Figure 5, for which the full-PDE and reduced model agree. Furthermore, in SI Figure 12 we verify that as the size of ribosome and mRNA increase, they are ejected from the chromosome and and become distributed according to their available volume profile (Remark 1). Furthermore, in SI-Figure 14–15, we demonstrate that these results hold independent of the binding and unbinding speed between ribosomes and mRNA (Remark 2).

It is well known that most mRNA-ribosome complexes exists in configurations with multiple ribosomes bounded (polysomes) [29, 30]. To capture the prevalence of these polysomes, we model the translation process accounting for the fact that one mRNA can be bound to multiple ribosomes. We first model the mRNA binding simultaneously to *N_r_* − 1 ribosomes to form the c_l_ complex, to which another ribosome binds to to form the fully loaded c_t_ complex. The leading ribosome with a complete peptide is released from c_t_ at a rate *κ_t_* to yield protein P. This is described by the following set of biochemical reactions:

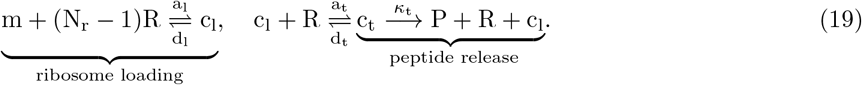

While the ribosome loading reaction in (19) is not in the form of the chemical reactions (1), which assume bimolecular reactions, we can nevertheless apply our results as follows (see SI Section 2.7 for details). Specifically, ribosome and mRNA profiles will still approach their available volume profiles (Remark 1), that is, 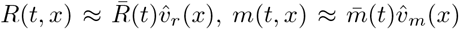, and 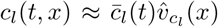 where 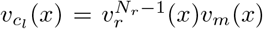 (recall (9)) and 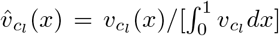. This is verified through simulations in SI Section 2.7, Figure 16. By virtue of the reactants in (19) mirroring their available volume profiles and (7b), we can approximate the BCF of the loading 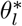 and translation 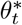 reactions in (19), given as

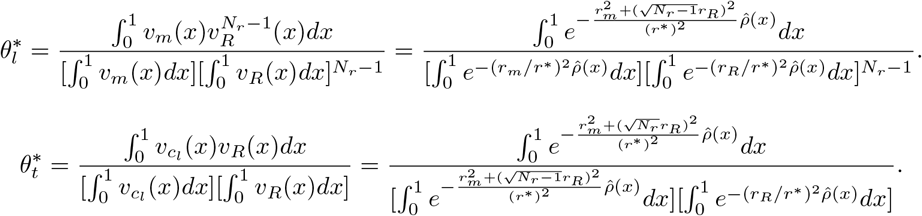

In SI Section 2.7, Figure 16-D, we show computationally that 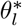 and 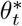 are good approximations to the BCF. At this point, we can write the ODE corresponding to this system of reactions and just modify the association rate constants by 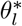 and 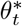, as shown in SI Section 2.7, Equation (66). Let 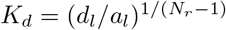, *K_t_* = (*d_t_* + *κ_t_*)/*a_t_*, *β_l_* = (*γ* + 1)/*d_l_* and *β_t_* = (*γ* + 1)/(*κ_t_* + *d_t_*), if 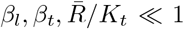 (dilution and mRNA degradation is much slower than the rate of ribosome unbinding and *K_t_* is sufficiently large comparer to 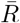 [52]), then a simple expression for the steady state protein concentration is given by

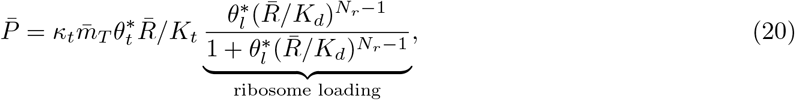

where 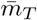 is given by (17).

In [13] it was estimated that *r_m_* = 20 nm and *r_R_* = 10 nm, which implies that *r_m_/r*^*^ ≈ 0.88 and *r_R_/r*^*^ ≈ 0.44. Assuming the average distance between ribosomes on an mRNA to be 70 nucleotides [31], for a 700 nucleotide mRNA (e.g., GFP or RFP), then we have *N_r_* = 10. Thus, for these values, 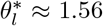 and 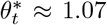. This implies that the forward rate in the reaction of 9 ribosomes binding to an mRNA (*a_l_*) is amplified by 56% and the rate at which an additional ribosome binds to this complex (*a_t_*) increases by 7% with respect to a well-mixed model. From (20), this would imply up to 67% increase in protein production with respect to a well-mixed model.

Taken together, these results suggest that while a well-mixed ODE model may be sufficient to describe transcription, it is not sufficiently descriptive to capture spatial effects on translation. In this case,the BCF should be incorporated in the ODE. Additionally, these results are indicative that for other processes in the cell where complexes of similar size as polysomes are formed, then spatial effects will likely be substantial.

#### 1.4.2 Gene expression regulation by transcription factors

Regulation of gene expression is often performed by transcription factors (TFs) [1]. A transcription factor can either enhance (for activators) or repress (for repressors) transcription. Spatial affects play an identical role in gene regulation via activators as they do in gene regulation via RNAP (Figure 5), thus we focus on transcriptional repressors. In this section, we model transcription regulation where a repressor P_r_ dimerizes to form dimer c_1_ (e.g., TetR dimerizes before binding to a gene [32]) and then blocks transcription of gene D that produces protein P. The biochemical reactions corresponding to this process are:

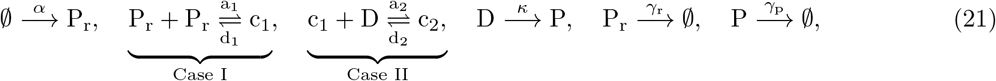

where *α* is the production rate of P_r_, *a*_1_ (*d*_1_) is the association (dissociation) rate constant to form the c_1_ complex, *a*_2_ (*d*_2_) is the association (dissociation) rate constant to form the c_2_ complex, *κ* is the catalytic rate constant to produce protein P, and *γ_r_* and *γ_p_* are the degradation rate constant of *P_r_* and *P*, respectively. Notice that we have lumped the transcription and translation process to produce P_r_ into one production reaction and similarly for P. From the results of Section 1.4.1, we know that 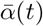 depends on the location where P_r_ is expressed (higher if its coding DNA is near the cell poles than mid-cell) and the size of it’s mRNA (higher for longer mRNAs). Similarly, *κ* depends on the location where P_r_ is expressed and it’s mRNA size. Using the results from Section 1.4.1 we can explicitly model these dependences, however, we opt not to do so to solely investigate the role of spatial effects on transcriptional repression. Since the repressor P_r_, freely diffuses, the dimerization reaction belongs to Case I. The gene D is spatially fixed and it is repressed by the freely diffusing c_1_, thus this interaction falls under Case II. We assume that the total concentration of D is conserved, so that *D_T_*(*x*) = *D*(*t, x*) + *c*_2_(*t, x*) and that *D_T_*(*x*) is localized at *x* = *x*^*^. The reduced ODE model corresponding to (21) obeys

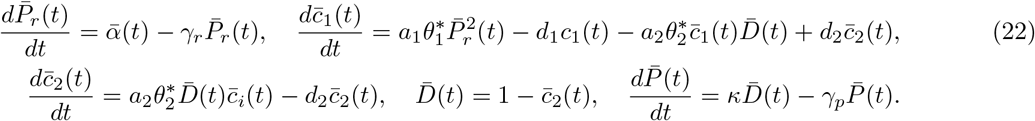

Concentration variables were nondimensionalized with respect to the space averaged total DNA 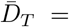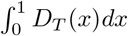. From our main result, the BCF’s are given by

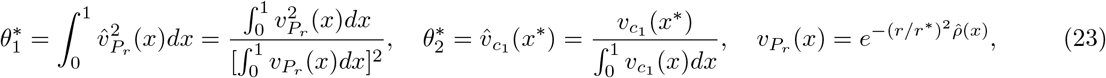

where *r* is the radius of gyration of P_r_, 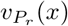 and 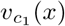 are the available volume profiles of P_r_ and c_1_, respectively, and from (9), 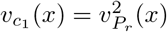.

We now consider the steady state behavior of system (22) by equating the time derivatives to zero. Specifically, we are interested in how the steady state levels of P is affected by the spatial quantities *r/r*^*^ and *x*^*^. From setting (22) to steady state, we obtain

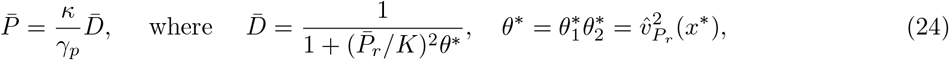

where 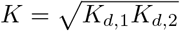 and *K_d,i_* = *d_i_*/*a_i_* for *i* = 1, 2. From (24) and (23), we observe that *θ*^*^ contains all the spatial information, which includes the size of P_r_ and the location of the target gene D. If *θ*^*^ = 1, then the protein concentration would be the same as the well-mixed model. The ratio 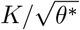 can be thought of as an effective disassociation rate constant of the repressor. If D is located near mid-cell (*x*^*^ ≈ 0 in (23)), then for *r/r*^*^ ≪ 1 we have *θ*^*^ ≈ 1 (see Figure 4), but as *r/r*^*^ increases, we have that *θ*^*^ < 1 and asymptotically approaches zero as *r/r*^*^ → ∞. Similarly, if D is located near the cell poles (*x*^*^ ≈ 1 in (23)), then for *r/r*^*^ ≪ 1 we have *θ*^*^ ≈ 1 (see Figure 4), but as *r/r*^*^ increases, we have that *θ*^*^ > 1. Thus, the efficacy of a transcriptional repressor regulating genes in the chromosome (cell-poles) decreases (increases) with TF size. Intuitively, this occurs because as the TF size increases, excluded volume effects will push it out of the chromosome onto the cell-poles (see Remark 1), thus interacting with DNA near the cell-poles more frequently than with DNA near mid-cell. Numerical simulations validate our predictions as shown in Figure 6, where increasing the transcription factor size leads to higher (lower) repression when the target DNA is localized at the cell poles (mid-cell) with respect to a well-mixed model. The simulation results also show agreement between the predictions of the full PDE ((70) in SI Section 2.8) and reduced ODE model (22) (as shown explicitly in SI Figure 19). Figure 17 in SI Section 2.8, further shows the temporal trajectories corresponding to Figure 6, also showing agreement between the full PDE model and the reduced ODE model. Finally, all our results hold independent of the binding and unbinding speeds of the transcription factor dimerizing and of the dimer binding to the DNA (Figure 18 in SI Section 2.8). In SI Section 2.12 we propose an experimental method to test the hypothesis that mid-cell genes are regulated less effectively than pole localized genes.

**Figure 6:**
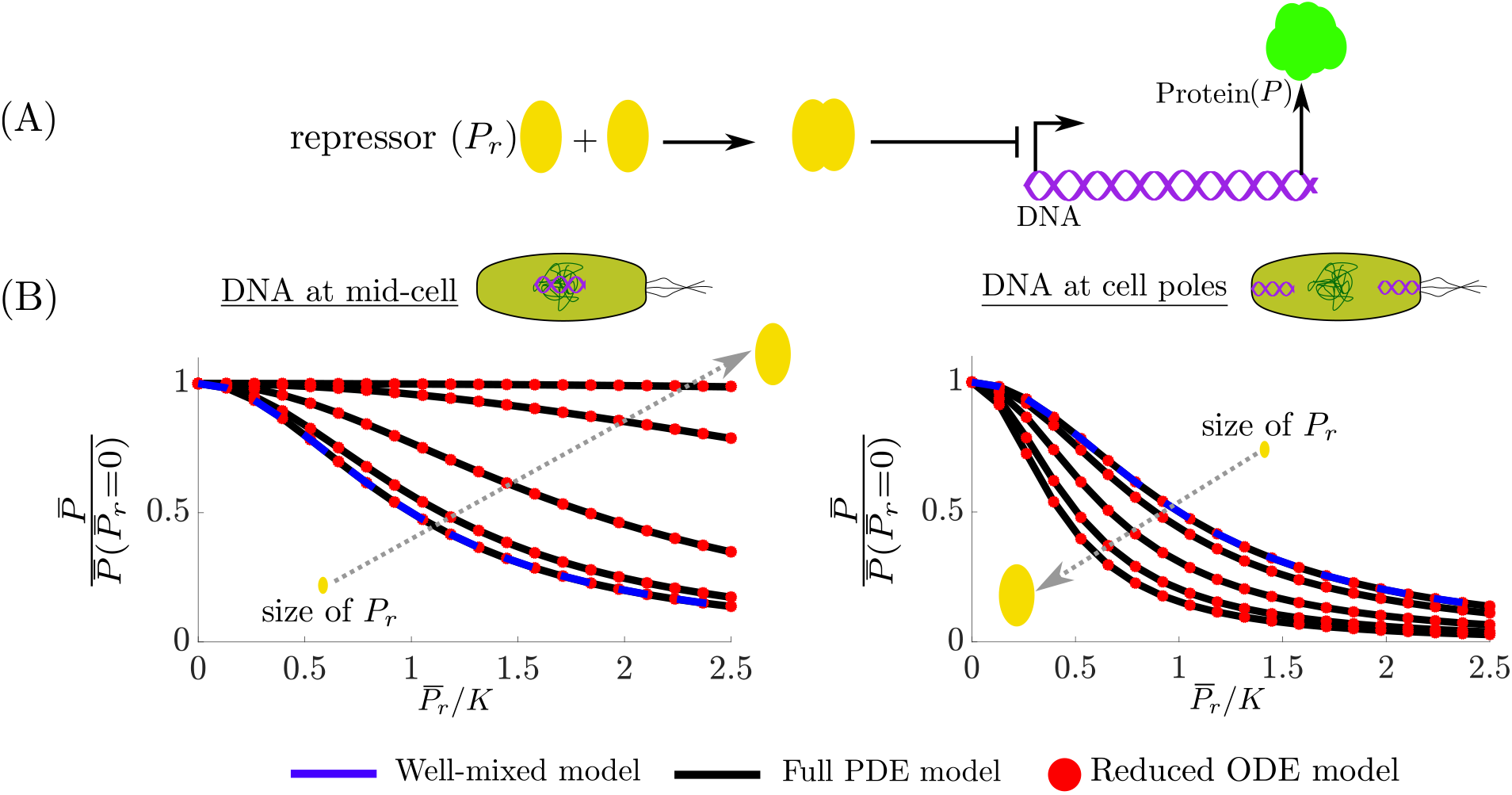
Spatial heterogeneity effects in transcriptional regulation. (A) The repressor P_r_ dimerizes and regulates the production of protein P. (B) The steady state space-averaged concentration per-cell of 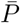 normalized by its value when 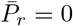 (24) for the PDE model ((70) in SI Section 2.8), the well-mixed model ((22) with 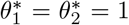), and the reduced ODE model (22) when the DNA is located near mid-cell (*x*^*^ ≈ 0 in (24)) and when the DNA is located at the cell-poles (*x*^*^ ≈ 1) for several sizes of P_r_. The parameter values and full simulation details are provided in SI Section 2.8.

The reactions in (21) can be easily extended to CRISPRi/dCas9 repression systems [33], where instead of two identical species dimerizing, we have two distinct freely diffusing species bind (dCas9 and guide RNA) to form the complex gRNA-dCas9, which targets a desired DNA sequence. Exploiting the insight gained from analyzing (21), we expect that due to the large size of dCas9 [34] (which is further augmented as it forms a complex with the gRNA), it will regulate pole localized DNA (e.g,. ColE1 plasmid DNA [9]) more efficiently than genes in mid-cell (e.g,. chromosomally integrated) and thus spatial effects are expected to be more significant when using CRISPRi/dCas9 in genetic circuit design. Specifically, based on approximate values found in the literature, we estimated that *θ*^*^ ≈ 1 for a transcription factor, while *θ*^*^ can range between 0.9 and 1.1 for dCas9-enabled repression. This indicates that a well-mixed model is appropriate for modeling transcription factor-enabled repression of gene expression but may not be sufficient to capture effects of spatial heterogeneity arising with larger repressing complexes such as with dCas9/gRNA (see SI Section 2.8 for details).

#### 1.4.3 Genetic Oscillator

As a final example, we consider the repressor-activator clock genetic circuit designed in [35] and shown in Figure 7-A. This circuit produces sustained oscillations if tuned within an appropriate parameter range [36, 1]. The circuit consists of two proteins P_a_ and P_r_. Protein P_a_, is an activator which dimerizes to form P_a,2_ and then binds to its own gene D_a_ to form complex c_a,1_ to initiate transcription. The dimer P_a,2_ also binds to the gene D_r_, which transcribes P_r_ to form complex c_a,2_ and initiates transcription. Protein P_r_, dimerizes to form P_r,2_ and then represses P_a_ by binding to D_a_ to form complex c_r_. The biochemical reactions corresponding to this circuit are:

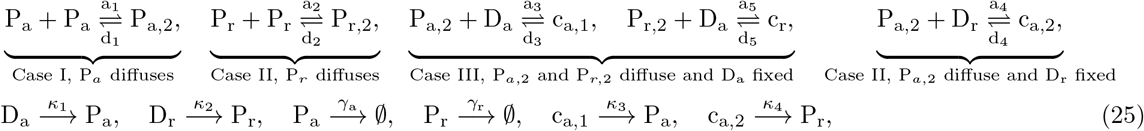

where *a_i_* (*d_i_*) for *i* = 1, …, 5 are association (dissociation) rate constants, *γ_a_* (*γ_r_*) is the degradation rate constant of P_a_ (P_r_) *κ*_1_ (*κ*_2_) is the basal rate at which gene D_a_ (D_r_) is transcribed, and *κ*_3_ (*κ*_4_) is the rate at which the DNA-transcription-factor complexes are transcribed for D_a_ (D_r_). We assume that the total concentration of D_a_ is conserved, so that *D_a,T_*(*x*) = *D_a_*(*t, x*) + *c*_*a*,1_(*t, x*) + *c_r_*(*t, x*) and that D_a,T_ is localized at 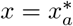. Similarly, we assume that the total concentration of D_r_ is conserved, so that *D_r,T_*(*x*) = *D_r_*(*t, x*) + *c*_*a*,2_(*t, x*) and that D_r,T_ is localized at 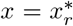. The reduced ODE model corresponding to (25) is given by:

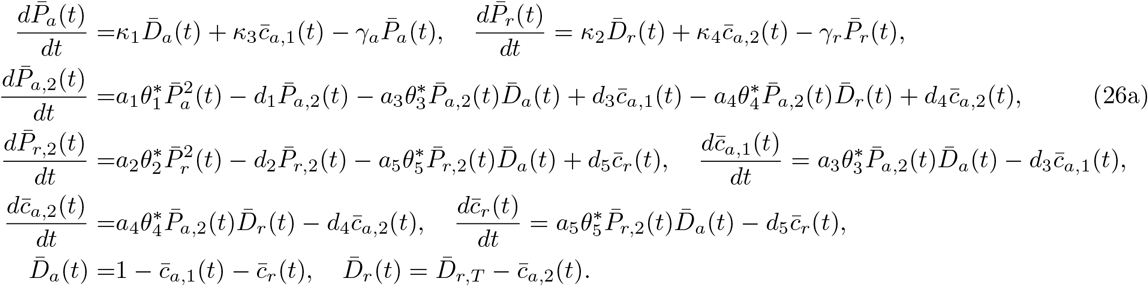

**Figure 7:**
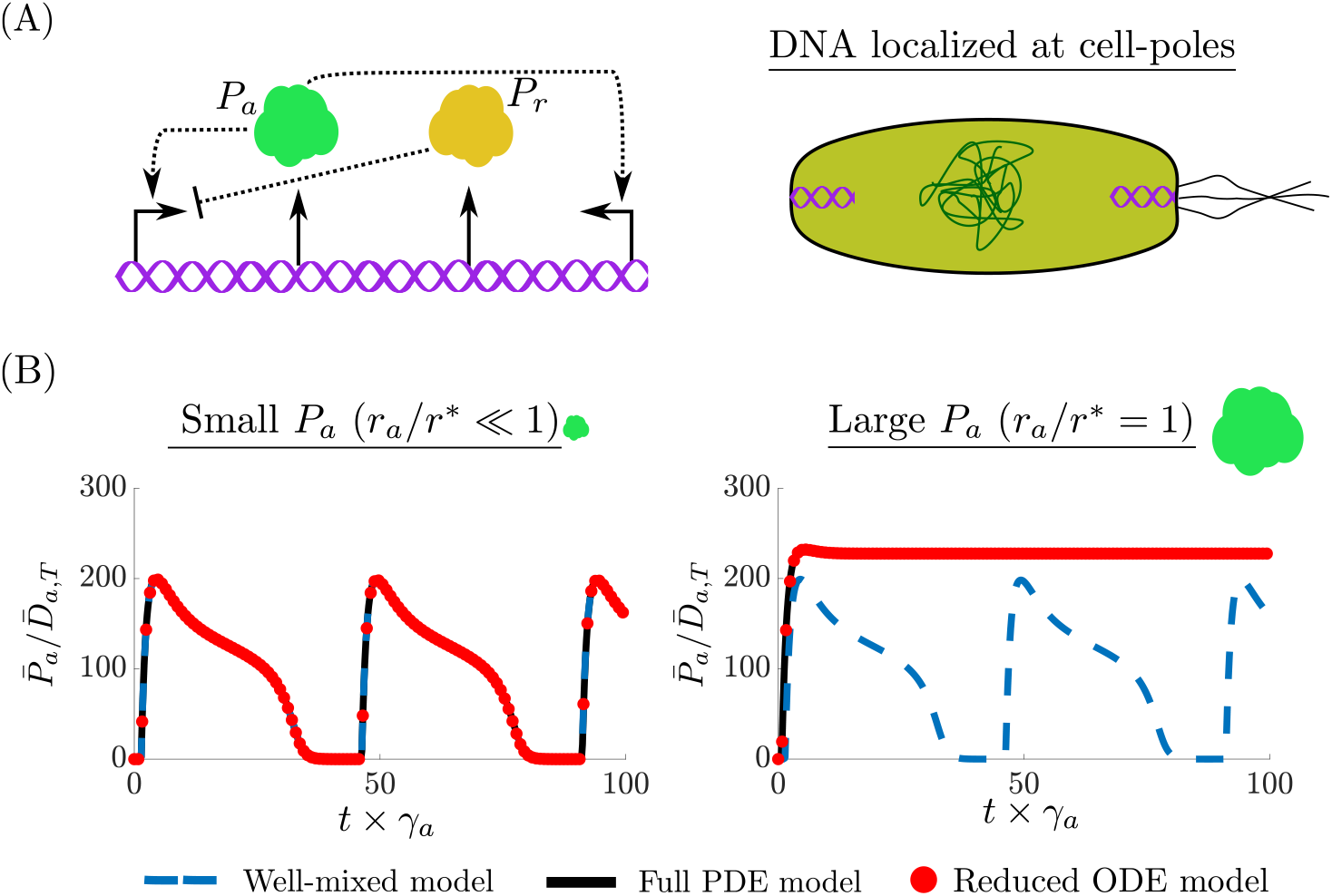
Spatial effects on the dynamics of genetic circuits. (A) The activator-repressor clock where P_r_ represses P_a_ and P_a_ activates itself and P_r_. Both proteins are expressed from the same cell-pole localized plasmid. (B) The temporal evolution of P_a_ is given for the full-PDE model ((72) in SI Section 2.9), the reduced ODE model (26), and the well-mixed model (same as (26) with 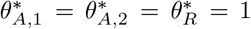). When P_a_ is small (*r_a_/r*^*^ ≪ 1), all three models predict sustained oscillations. When P_a_ is large (*r_a_/r*^*^ = 1), the full-PDE model and the reduced ODE model predict the oscillations will cease. For both simulations 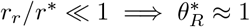. The full simulation details and parameter values are given in SI Section 2.9.

Concentration variables were nondimensionalized with respect to the space averaged total DNA 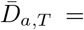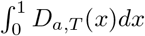. Applying our main result, the BCF’s are given by

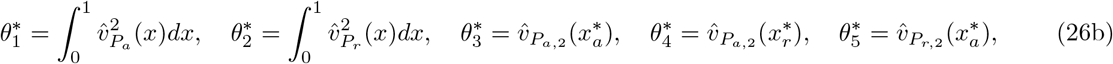

where 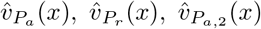, and 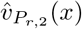 are the normalized available volume profiles (i.e,. 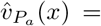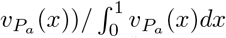 of P_a_, P_r_, P_a,2_, and P_r,2_, respectively. From (9), notice that 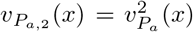 and 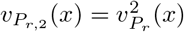. The available volume profiles are given by

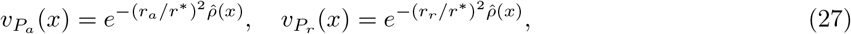

where *r_a_* and *r_r_* are the radius of gyration of P_a_ and P_r_, respectively. Approximating 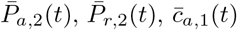, 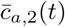 and 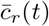 at their quasi-steady state (since *d_i_* ⨠ *γ_a_*, *γ_r_* for *i* = 1, …, 5, [1]), we obtain

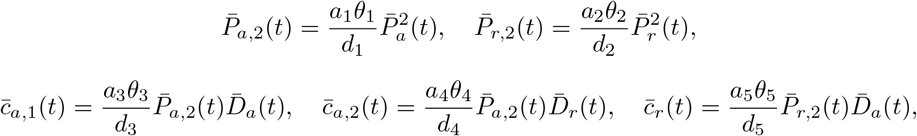

and, therefore, we can further reduce (26) to

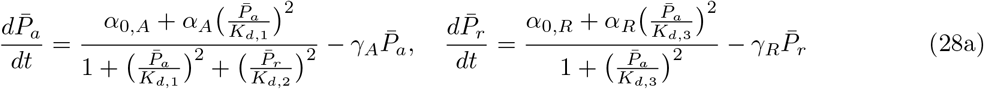

where 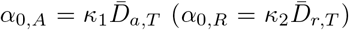 is the basal production rate of 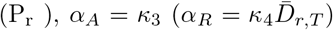 is the additional production rate of P_a_ (P_r_) due to activation from P_a_, and

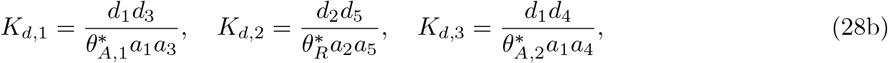

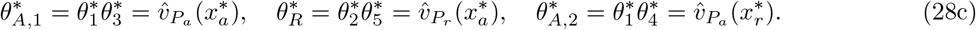

The form of the dynamics given by (28) was theoretically analyzed in [36, 1], and it was shown that the values of *K_d,i_* for *i* = 1, 2, 3, were critical in determining whether sustained oscillations occur. From (28b), these parameters depend on (28c) and thus on the size of P_a_ and P_r_ through the available volume profiles (27) and the location of D_a_ and D_r_ (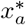 and 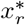). Numerical simulations demonstrate how these spatial parameters affect circuit behavior. In our simulation setup, the parameters are chosen such that the well-mixed model ((26) with 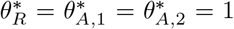) oscillates, the DNA of P_a_ and P_r_ are localized at the cell poles and have the same copy number (i.e., 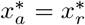 and 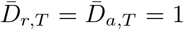), the size of P_r_ is chosen to be small *r_r_/r*^*^ ≪ 1 (thus 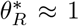), and the size of P_a_ is varied (thus varying 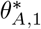 and 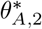). Since D_a_ is localized at the cell poles, it implies 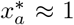 and from (28c), we observe that if 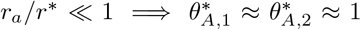 and 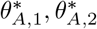 increase as *r_a_/r*^*^ increases. The results of these simulations are shown in Figure 7. When *r_a_/r*^*^ ≪ 1 the full PDE model ((72) in SI Section 2.9), the reduced ODE model (26), and the well-mixed model are all in agreement and sustained oscillations are observed. By contrast, when *r_a_/r*^*^ = 1, the PDE and reduced model (which are in agreement with each other as explicitly shown in SI Figure 21) predict that sustained oscillations will no longer occur. Furthermore, in SI Section 2.9 we demonstrate in Figure 22 that indeed as the size of P_a_ increases it is excluded from the chromosome onto the cell poles while the spatial profile of P_r_ is homogeneously distributed throughout the cell since *r_r_/r*^*^ ≪ 1 (Remark 1).

## Discussion

We derived a reduced order ODE model of genetic circuits with the same dimension as traditional ODE well-mixed models; yet, it captures effects of spatial heterogeneity within bacterial cells (11). In particular, our reduced model is the same as a well-mixed model where all the association rate constants are multiplied by the binding correction factor (BCF). This factor depends on the size and location (if fixed in space) of the reacting species, according to an analytical formula that we derived from first principles (11b) and its value can be estimated experimentally through simple procedures (SI Section 2.11). We have mathematically demonstrated that this reduced order model is a good approximation of the space-averaged dynamics resulting from a reaction-diffusion PDE model under the assumption of fast diffusion. It can therefore be used in place of PDE models, providing substantial advantages for both simulation and mathematical analysis.

We applied this model to analyze the effects of spatial heterogeneity on core biological processes and genetic circuits. Specifically, motivated by the fact that DNA, ribosomes, and mRNA have been shown to localize within the cell [4, 14, 5], we analyzed the transcription and translation processes. We determined that mRNA levels are lower (higher) when the gene is localized near the mid-cell (cell poles). We also showed that when the target gene of a transcriptional repressor is near mid-cell (cell poles) the effective repression is lower (higher) with respect to that of the well-mixed model. This discrepancy is amplified as the size of the transcription factor increases. The extent of these spatial effects depends on how different the value of the BCF is from unity. Based on parameters found in the literature, we determined that for the processes of transcription and its regulation the BCF should be close to unity and hence a well-mixed ODE model should be sufficient. However, in situations where the nucleoid is highly compacted (from overexpressing mRNA [13] or translational inhibition [37]), we expect that the available volume profile (2) approaches small values and, as a consequence, the value of the BCF can substantially deviate from unity (11b).

Our results provide additional interpretations of well-known biological phenomena. For example, it has been shown that the expression rate of chromosomal genes depends on the locus where the gene is inserted [38]; that the nucleoid dynamically changes shape to control gene expression and transcription regulation [4, 39] (e.g., see SI Section 2.13 for how a time varying chromosome density modulates the BCF); and that coregulation and coexpression among genes depends on their spatial distance [40]. For a fixed amount of mRNA, we showed that spatial heterogeneity leads to higher translation rates since both mRNA and ribosome are pushed out of the chromosome into a smaller region near the cell poles, which results in larger effective binding affinity. How larger, it depends on the value of the BCF. For a polysome with 10 translating ribosomes, the value of the BCF can deviate from unity by 56% in the ribosome loading step and by 7% in the peptide release step. These estimates are believed to be conservative since we did not account for the exclusion effects from the peptide chains attached to the translating ribosome, which will result in even more pronounced spatial effects. Therefore, a well-mixed model may not be sufficient to capture the effects of spatial heterogeneity on translation.

Our modeling framework can be easily extended to other aspects of gene expression. For example, we may consider co-transcriptional translation [41]. In this case, as a result of translation being localized at the gene location, the effective ribosome binding site strength will also depend on gene location through the BCF. We may also consider the role of spatial heterogeneity on orthogonal translational machinery [42]. From our models, we predict that one can tune the rate at which orthogonal ribosomes are formed by creating larger synthetic 16S rRNA. Furthermore, once the production of orthogonal ribosomes is placed in a feedback form to decouple genetic circuits [42], our framework suggests that the feedback efficiency may depend on the spatial location of the synthetic 16S rRNA gene. The value of the parameter *r*^*^, whose squared value is inversely proportional to the average chromosome density (2), is critical in determining the extent of spatial effects. In this study we indirectly estimated a value of *r*^*^ based on [13]. However, a more comprehensive study should be conducted to estimate *r*^*^ for several contexts (SI Section 2.11), or equivalently to estimate extent of excluded volume effects, which may easily be performed via superresolution imaging [14].

In summary, this paper provides a general and convenient modeling framework to account for DNA localization and excluded volume effects on intracellular species dynamics. While other phenomena contributing to intracellular spatial heterogeneity, such as crowding [43], sliding, hopping, and dimensionality [17], exist, this is a first step towards creating a general framework to modify current models to capture spatial information. Our model can be used both as an analysis and a design tool for genetic circuits, in which variables such as gene location and regulator size may be considered as additional design parameters.

## Author Contributions

C.B. performed the research, developed the mathematical analysis, and wrote the article. D.D.V. designed the research, assisted with the mathematical analysis, and edited the article.

## Acknowledgments

This work was supported in part by NSF Expeditions, Grant Number 1521925, AFOSR grant FA9550-14-1-0060, the NSF Graduate Research Fellowships Program, and the Ford Foundation Predoctoral Fellowship. We thank Jean-Jacques Slotine for the technical discussions on contraction theory. We thank Theodore Grunberg and Yili Qian for reviewing the article and helpful discussions

## Supplementary Material

### 2 Time Scale Separation Proofs

#### 2.1 Preliminaries

##### Notation

Let *z* = [*z*_1_, …, *z_n_*]^*T*^ ∈ ℝ^*n*^ (where superscript *T* denotes the transpose operation) and the *j*-th component of *z* is denoted by *z^j^*. A vector of zeros is denoted as 0_*n*_ = [0, …, 0]^*T*^ ∈ ℝ^*n*^ and we use *A* = diag(*v*) ∈ ℝ^*n*×*n*^ to refer to a square matrix with all zeros in the off-diagonals and diagonal elements specified by the vector *v*. The (*j, k*)-th element of matrix *A* is denoted by *A^j,k^* and *I_n,n_* is the identity matrix acting on ℝ^*n*^. Let 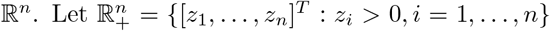 denote th positive orthant of ℝ^*n*^. For a Hilbert space *H* with inner product (·, ·)_*H*_, we denote the norm as 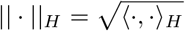. Finally, let overbars denote spatial integration 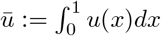.

###### Definition 1

(Linear Differential Operator) Let *v*(*x*) : [0, 1] → ℝ_+_ be a smooth function, and consider the following linear differential operator:

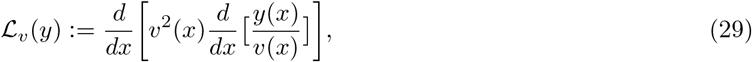

with domain 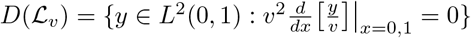:

Next, we introduce the Hilbert space 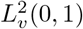 that we use for our analysis. This space is isomorphic to *L*^2^(0, 1) however, the operator (29) is self-adjoint with respect to the inner product in 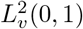.

###### Definition 2

(Weighted *L*^2^(0, 1) space) For smooth *v* : [0, 1] → ℝ_+_ we denote 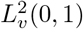 as a weighted space of the square integrable functions such that 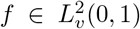 if and only if : 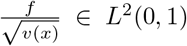. The inner-product is defined as 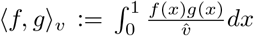, for 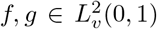 and 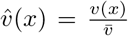. Furthermore, let *v* = [*v*_1_(*x*),…, *v_n_*(*x*)]^*T*^, where *v_i_*(*x*) : [0, 1] → ℝ_+_ is smooth, then 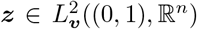 if 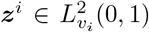 for *i* = 1,…, *n*, and the inner and the inner product in this space is defined as 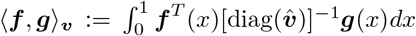, for 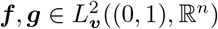, where 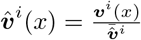.

###### Remark 3

(Norm equivalence between *L*^2^ and 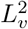) For smooth *v* : [0, 1] → ℝ_+_, let 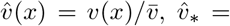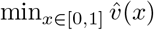 and 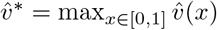, the norms defined on 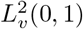 and *L*^2^(0; 1) are related by

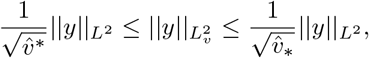

for any 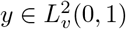. Thus, when performing convergence analysis we may use the norms defined in 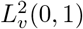 and *L*^2^(0, 1) interchangeably.

###### Lemma 1

*(Negative semi-definite and self-adjoint Operator) For smooth v* : [0, 1] → ℝ_+_ *let* 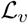 *be as in Definition 1. This operator has the following properties:*

I. *Has countably many, real, and distinct eigenvalues such that λ*_1_ > … > … *λ_n_* > *and* lim_*n*→∞_ *λ_n_* = −∞
II. *The set of corresponding eigenfunctions* {*ψ_i_*(*x*)} *form a complete orthonormal basis for* 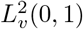 *(Definition 2)*
III. *λ*_1_ = 0 *and* 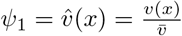

*Proof:* The proof of (I) and (II) follow from Sturm-Liouville theory [44]. To prove (III), we take the weighted inner product of both side of 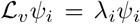 with *ψ_i_* we use orthonormality, use integration by parts, and apply the boundary conditions:

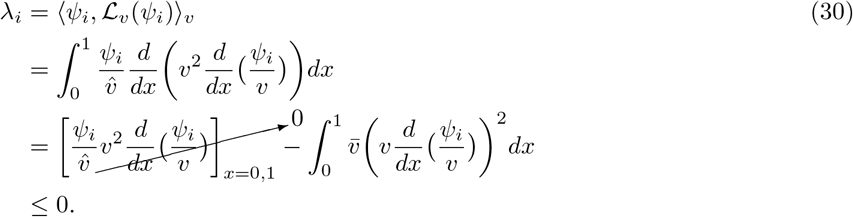

The maximum of (30) is achieved (*λ*_1_ = 0) for 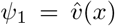, since substituting 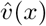 directly into 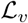, one observes that 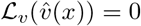 and we have that 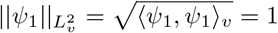, therefore *λ_i_* < 0, ∀*i* > 1.

The following will introduce the notion of contracting dynamical systems. A dynamical system is said to be contracting within an open and connected subspace of the state space, if all trajectories starting within this region converge exponentially to each other. We provide sufficient conditions to guarantee that a dynamical system is contracting, and finally show that contracting systems have a particular robustness property. The robustness property will be exploited several times in our analysis to perform our model reduction.

###### Theorem 1

*Let* 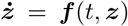 *be a dynamical system in the Hilbert space H* = ℝ^*n*^ *where f is a smooth nonlinear function. A dynamical system is said to be contracting within an open and connected subspace of the state space χ* ⊆ *H, if all trajectories starting within this region converge exponentially to each other. A sufficient condition for a system to be contracting in H is the existence a uniformly positive definite matrix P*(*t, z*) ∈ ℝ^*n*×*n*^ *and constant λ* > 0 *such that*

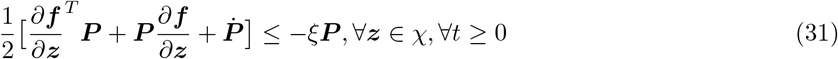

*where ξ is the contraction rate of the system and* 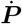 *is the total time derivative of P*.

*Proof:* See Theorem 2 in [45].

###### Lemma 2

*(Hierarchies of contracting systems) Let*

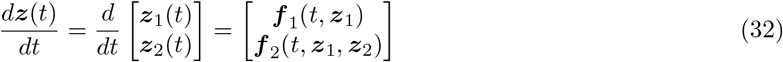

*be dynamical systems in the Hilbert space H* = ℝ^*n*+*m*^ *where f*_1_ : [0, ∞) × ℝ^*n*^ → ℝ^*n*^ *and f*_2_ : [0, ∞) × ℝ^*n*^ × ℝ^*m*^ → ℝ^*m*^ *are smooth nonlinear functions. Then, sufficient conditions for* (32) *to be contracting in χ* = *χ*_1_ ⊕ *χ*_2_, *where χ*_1_ *and χ*_2_ *are open connected subspaces of* ℝ^*n*^ *and* ℝ^*m*^, *respectively, are*

I. 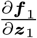 *satisfies* (31) *in χ*_1_ *for some P*_1_ ∈ ℝ^*n*×*n*^ *such that* 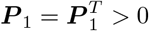
II. 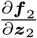*satisfies* (31) *in χ*_2_ *for some P*_2_ ∈ ℝ^*m*×*m*^ such that 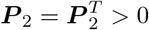
III. 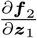 *is uniformly bounded for all t* ≥ 0, *z*_1_ ∈ *χ*_1_, *and z*_2_ ∈ *χ*_2_

*Proof:* See hierarchal structures in [45] and applied in [46]

###### Lemma 3

*(Robustness property of contracting systems) Assume that* 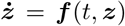 *satisfies the conditions of Theorem 1, for some P*(*t, z*) = *P*(*t, z*)^*T*^ > 0 *in a region χ* ⊆ *H, and thus it is contracting with some contraction rate ξ. Furthermore, assume that there exists a positive constant λ*^*^ (*λ*_*_) *that upper (lower) bounds the maximum (minimum) eigenvalue of P for all t* ≥ 0. *Consider the “perturbed” system* 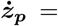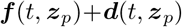 *and suppose there exists L*_1_, *L*_2_, *ζ, ϵ* > 0 *such that* ζ/ϵ > ξ *and* ‖*d*(*t, z_p_*)‖_*H*_ ≤ *L*_1_*e*^−*ζ*/*ϵt*^+*L*_2_*ヵ for all t* ≥ 0 *and z_p_* ∈ χ. *Then, there exists L*^*^, *ϵ*^*^ > 0 *such that for all z*(0), *z_p_*(0) ∈ *χ and* 0 < *ϵ* < *ϵ*^*^, *the solutions z_p_*(*t*) *and z*(*t*) *satisfy*

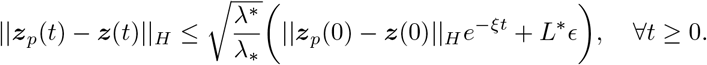

*Proof:* This follows from the result of Seciton 3.7 Result vii in [45]. Let *R_z_*(0) be the length of the straight path connecting *z_p_*(0) and *z*(0), that is, 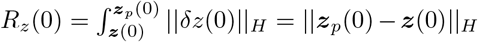, where *δz* is the virtual displacement (see [45] for details. Each point in the straight path connecting *z_p_*(0) and *z*(0) evolves in time and we denote the length of the path connecting these points be given by *R_z_*(*t*). Precisely, 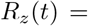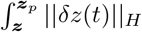. Notice that

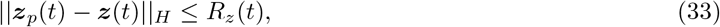

since the straight line path is the shortest path between the trajectories (they would be equal if the initial straight segment remained a line for all time, but this is only the case for constant vector fields). Let 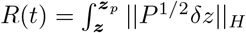. Notice that

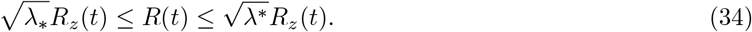

From Equation 15 in [45], we have that 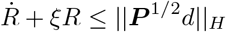 and thus

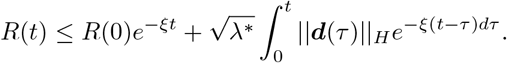

From (34), we have that

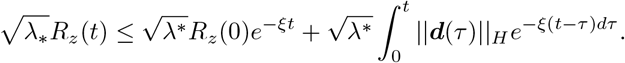

Finally, by (33), this implies that

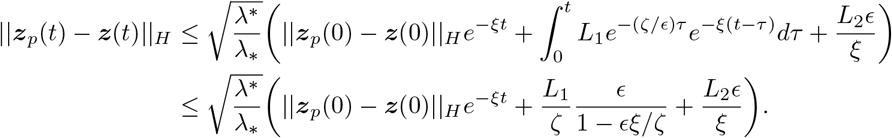

The desired result holds for *ϵ*\* = *ζ/*(2*ξ*) and *L*\* = 2*L*_1_/*ζ* + *L*_2_/*ξ*.

### 2.2 Solutions of diffusing states converge to the null space of the differential operator 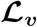

Let 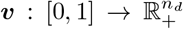 be a smooth vector-valued function and *V*(*x*) = diag(*v*(*x*)). For the state vectors 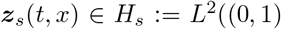 ∀*t* ≥ 0 and 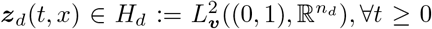, consider the following reaction-diffusion system:

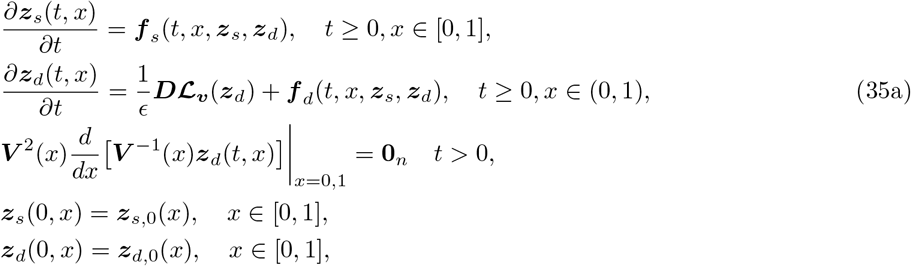

where 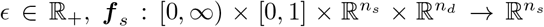 and 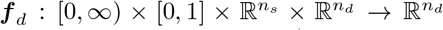 are smooth functions, 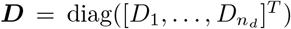 such that *D_i_* > 0 for all 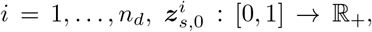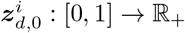, and

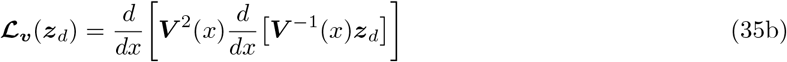

is such that 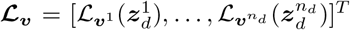 where 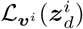 is an in Definition 1. The elements of *z_d_* may be thought of as “freely diffusing” and those of *z_s_* as “spatially fixed”. For the general system (35), we show in this section that the spatial profiles of the diffusing species *z_d_* approach those of *v*(*x*), that is, the null space of 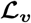 when *ϵ* ≪ 1.

#### Assumption 1

Consider the system given by (35)

I. There exists a positively invariant set 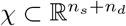 such that if 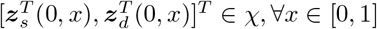, then 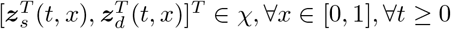.
II. There exists a positive constant *M* > 0 such that

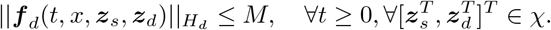

Let 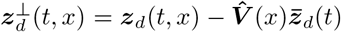 where 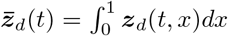 and 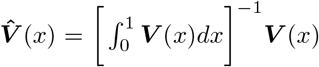. The dynamics of 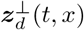 are given by

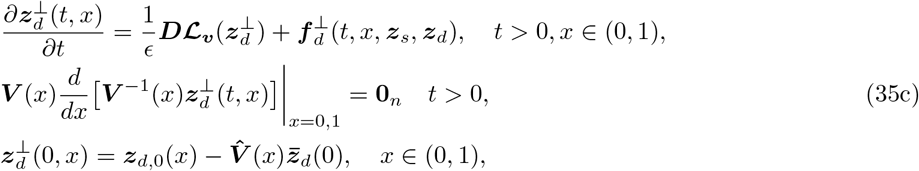

where

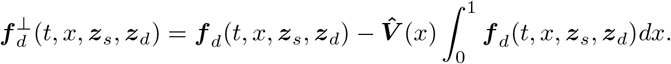

The following theorem is a direct consequence of the operator 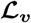 from Definition 1 being self-adjoint and negative semi-definite. This result will show that the infinite dimensional left over dynamics 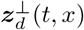 become order *ϵ* after a fast transient.

#### Theorem 2

*Consider the system defined by* (35) *Suppose that Assumption 1 holds and* 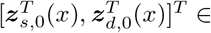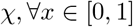, *where χ is defined in Assumption 1-I. Then, there exists ζ, L*_⊥_ > 0 *such that for all ϵ* > 0, *the solution* 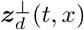 *of* (35c) *satisfies*

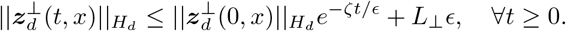

*Proof:*

Let 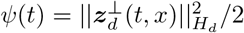 and *τ* = *t/ϵ* thus

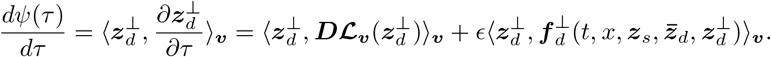

The following proof uses a similar logic as the the min-max theorem for matrices [49] to derive and upper bound for 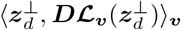. Let *λ_i,j_* and *ψ_i,j_* denote the *j*-th eigenvalue and eigenfunction, respectively, of 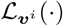 (i.e,. 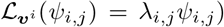). Recall that {*ψ_i,j_*} forms a complete orthonormal basis for 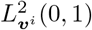 (Lemma 1-II). From the orthonormality of the eigenfunctions, linearity of 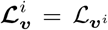, and the ordering of eigenvalues (Lemma 1-I), notice that:

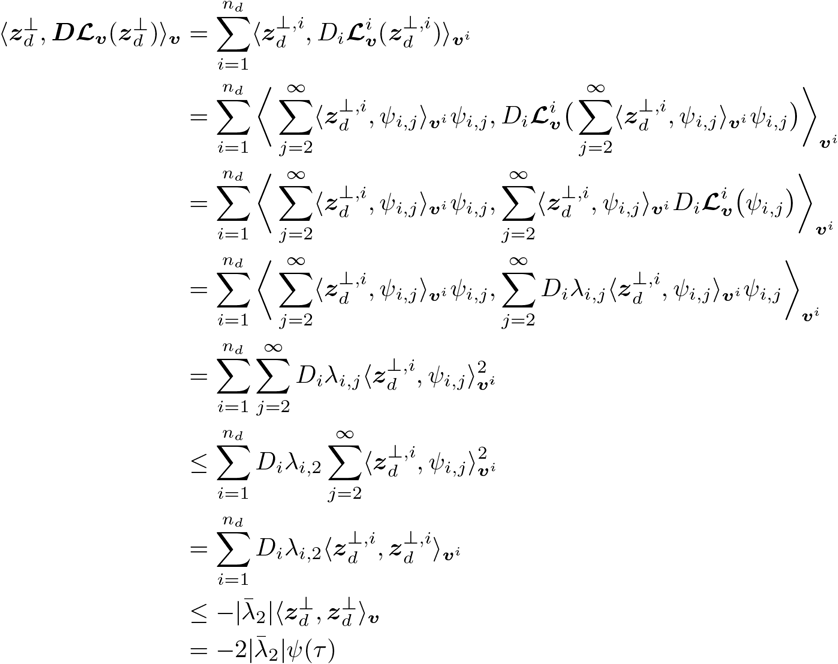

where 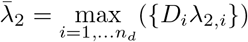. The fact that 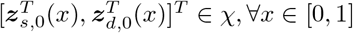 implies that 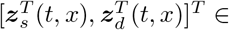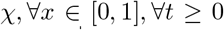 by Assumption 1-I. By the cauchy-Schwarz inequality, Assumption 1-II, and the fact that 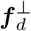 is a projection of *f_d_* (thus 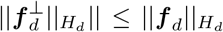) we have that 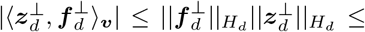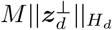. Let *ν* ∈ (0, 1) thus

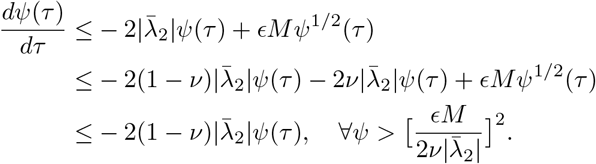

Therefore,

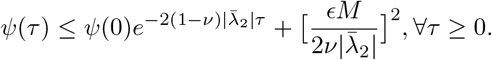

Finally,

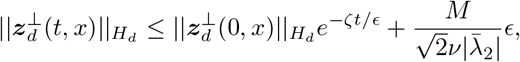

where 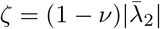 and 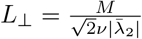.

### 2.3 Model Reduction of Enzymatic-like Reaction In the Limit of Fast diffusion

The spatial-temporal dynamics of E_i_, S, and c_i_ as described by the biochemical reactions (1) in the main text, are given in dimensionless form by

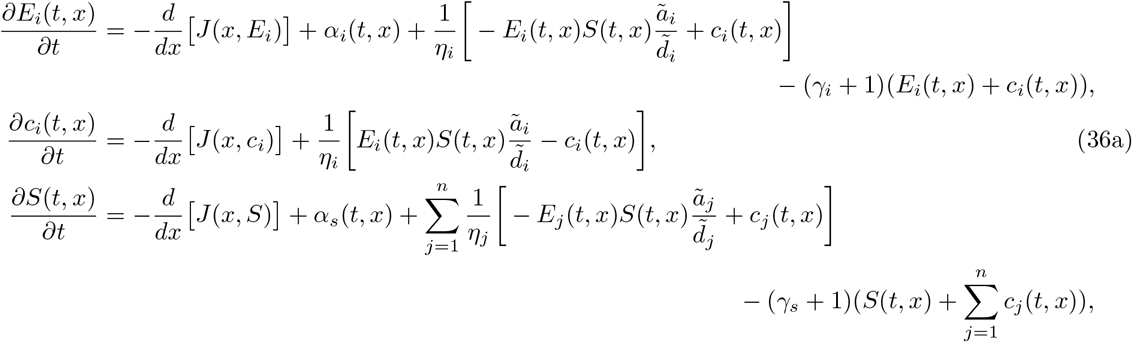

where the flux term and the boundary conditions are given in Table 2 for three cases of interest, 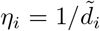, and 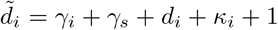. If applicable, 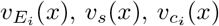 are the available volume profiles of E_i_, S, and c_i_, respectively, and let

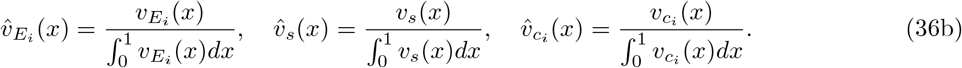

**Table 2:** The flux dynamics and the boundary conditions corresponding to (36) for Cases I-III. Here *v_E,i_*(*x*), *v_S_*(*x*), and *v_c,i_*(*x*), are the available volume profiles of E_i_, S, and c_i_, respectively, and 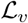 is as in Definitions 1. The parameters 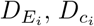, and *D_s_* are the enzyme, complex, and substrate diffusion coefficients, respectively, *ϵ* is a dimensionless parameter that captures the speed of diffusion (with respect to dilution). A species being spatially fixed translates to the flux being zero throughout the whole spatial domain. In Case II, we denote the location of the fixed species E_i_, as 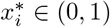 for *i* = 1, …, *n*. In Case III, we denote the location of the fixed species S, as 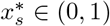.

| Case | I: All species diffuse | II: Substrate diffuses and enzymes fixed | III: Enzymes diffuses and substrate fixed |
| --- | --- | --- | --- |
| Dimensionless Flux | $-\frac{d}{dx} [J(x, E_i)] = \frac{1}{\epsilon} \chi_{E_i} \mathcal{L}_{v_{E_i}}(E_i)$<br>$-\frac{d}{dx} [J(x, S)] = \frac{1}{\epsilon} \mathcal{L}_{v_S}(S)$<br>$-\frac{d}{dx} [J(x, c_i)] = \frac{1}{\epsilon} \chi_{c_i} \mathcal{L}_{v_{c_i}}(c_i)$ | $-\frac{d}{dx} [J(x, E_i)] = 0$<br>$-\frac{d}{dx} [J(x, S)] = \frac{1}{\epsilon} \mathcal{L}_{v_S}(S)$<br>$-\frac{d}{dx} [J(x, c_i)] = 0$ | $-\frac{d}{dx} [J(x, E_i)] = \frac{1}{\epsilon} \mathcal{L}_{v_{E_i}}(E_i)$<br>$-\frac{d}{dx} [J(x, S)] = 0$<br>$-\frac{d}{dx} [J(x, c_i)] = 0$ |
| Boundary conditions | $J(0, E_i) = J(1, E_i) = 0$<br>$J(0, c_i) = J(1, c_i) = 0$<br>$J(0, S) = J(1, S) = 0$ | $J(0, S) = J(1, S) = 0$ | $J(0, E_i) = J(1, E_i) = 0$ |
| $\epsilon$ | $(\mu L^2)/D_s$ | $(\mu L^2)/D_s$ | $(\mu L^2)/D_{E_1}$ |
| Dimensionless diffusion | $\chi_{E_i} = D_{E_i}/D_s, \chi_{c_i} = D_{c_i}/D_s$ | | |
| Location of fixed species | N/A | $x_i^*$ | $x_s^*$ |

We denote 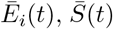, and 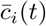 to be the space averaged enzyme, substrate, and complex concentrations, respectively (e.g., 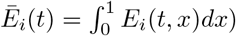. The dynamics governing these space averaged variables are derived by integrating (36) in space and applying the boundary conditions and are given by:

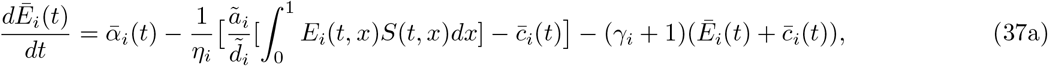

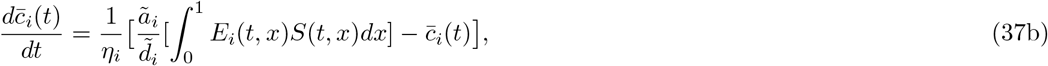

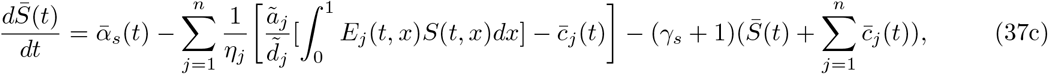

where overbars denote spatially averaged variables. We make the follow assumptions necessary to state our main result

#### Assumption 2

Consider (36), for *i* = 1, …, *n*. assume that

I. the functions *α_i_*(*t, x*) and *α_s_*(*t, x*) are smooth in each argument
II. there exists constant 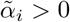 such that 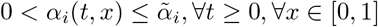
III. there exists constant 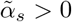 such that 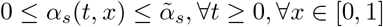
IV. the functions *v_E,i_*(*x*), *v_S_*(*x*), and 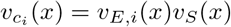 (as in Table 2) are smooth, strictly greater than zero, and bounded above by unity.

The following assumption makes precise what it means for the spatially fixed species E_i_ (in Case II) and S (in Case III) to be spatially localized at 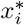 and 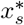 respectively.

#### Assumption 3

(Localization of spatially fixed species) Consider the system (36). Let 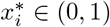 for Case II and 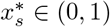 for Case III, be given by Table 2. Let *δ*\* = min(*x*_1_, …, *x_n_*, 1−*x*_1_, …, 1−*x_n_*) for Case II and 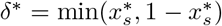 for Case III. We assume that for a given *δ* > 0 such that *δ* < *δ*\*, the functions *α_i_*(*t, x*) and *α_s_*(*t, x*) satisfy

- for Case II: *α_i_*(*t, x*) ≤ *δ*, for all 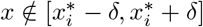, 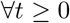
- for Case III: *α_s_*(*t, x*) ≤ *δ*, for all 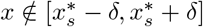, 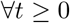,

Furthermore, we assume for Cases I-III, that there exists 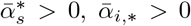 and 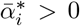 for *i* = 1, …, *n* independent of *δ*, such that 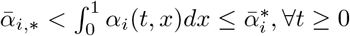 and 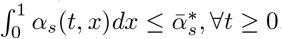.

The following definition will provide the candidate reduced model to approximates (37).

#### Definition 3

(Reduced space-averaged dynamics) Let 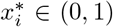 for Case II and 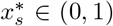 for Case III, be given by Table 2. For *i* = 1, …, *n*, consider the system

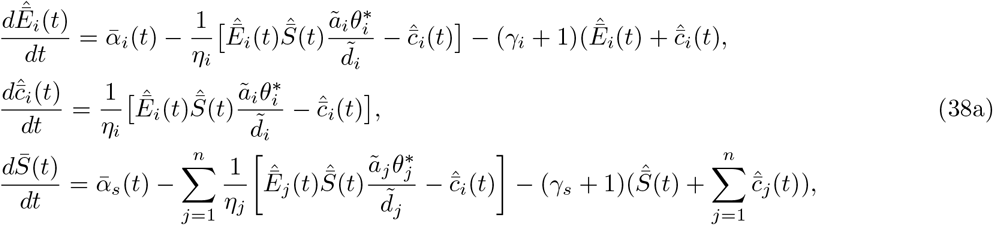

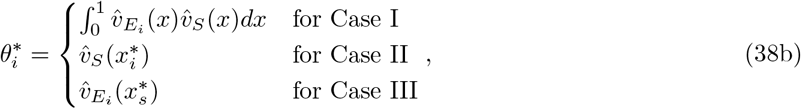

where 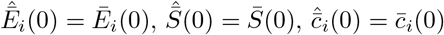, as given by (37),and 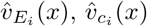, and 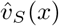 are given by (36b).

#### Theorem 3

*Consider the system* (36) *and let*

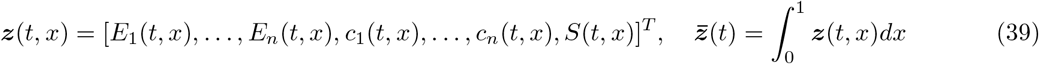

*Let* ϵ > 0, 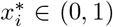 *and* 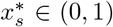 *be given for Case I-III by Table 2. Let* 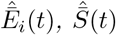 *and* 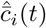 *be as in Definition 3 and let*

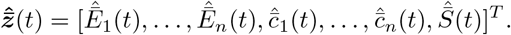

*Suppose that Assumptions 2 holds for Cases I-III. Then, there exists L*_1_, *ϵ*\* > 0, Ω_*z*_ ⊂ *L*^2^((0, 1), ℝ^2*n*+1^) *and* 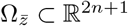 *such that for all* 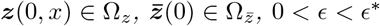, *the solutions* 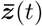 *and* 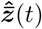 *satisfy*

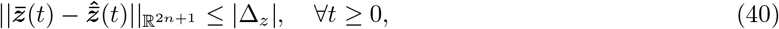

*where* |Δ_*z*_| = *L*_1_*ϵ for Case I. For Cases II-III, if in addition, Assumption 3 holds for all* 0 < *δ* < *δ*\*, *there exists L*_3_ > 0 *such that for all* 0 < *δ* < *δ*\*, there exists L_2_(*δ*) *and* Ω_*z,δ*_ ⊂ *L*^2^((0, 1), ℝ^2*n*+1^) *such that* (40) *holds for all* 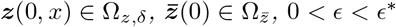 *with* |Δ_*z*_| = *L*_2_(*δ*)*ϵ* + *L*_3_*δ*.

#### Remark 4

The constant *L*_2_(*δ*), guaranteed to exist in Theorem 3 for Cases II-III, depends on *δ* Therefore, for a given *η* > 0, if one wishes to to have 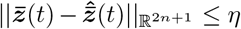, ∀*t* ≥ 0, then one would chose *δ* such that *L*_3_*δ* < η and then choose *ϵ* sufficiently small (depending on *δ*) such that *L*_2_(*δ*)*ϵ* + *L*_3_*δ* ≤ *η*.

#### Remark 5

The set Ω_*z*_ guaranteed to exist in Theorem 3, depends on *δ* in Cases II-III and this dependence is made precise in the proof.

#### Road map of proof

The rest of this section is dedicated towards proving Theorem 3. We first apply Theorem 2 to show that the spatial profile of a freely diffusing species converges to its available volume profile (e.g,. *v_E,i_*(*x*), *v_S_*(*x*), and 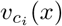) exponentially fast in the time scale associated with diffusion. If the localization assumption holds, then (37) has the form described by Definition 3, but with additional “disturbance” terms of order *E* and *δ* We proceed to demonstrate the system described in Definition 3 is contracting and apply the robustness property of contracting systems (Lemma 3) to show closeness between its solutions and those of (37).

The following result will define a positively invariant and bounded subset of R2*n*+1 such that solutions to (36) starting within this set at *t* = 0, remain within this set for all times and spatial values. To apply Theorem 2 to (36), the existence of such a positively invariant set is required by Assumption 1.

##### Claim 1

Consider the system given by (36), with Cases I-III specified by Table 2. Suppose Assumptions 2 holds. Let 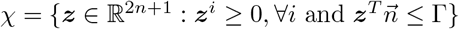, where

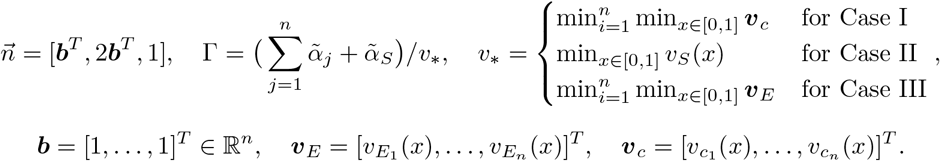

Let *z*(*t, x*) be given by (39), if *z*(0, *x*) ∈ *χ*, ∀*x* ∈ [0, 1], then *z*(*t, x*) ∈ *χ*, ∀*t* ≥ 0, ∀*x* ∈ [0, 1]. Thus *χ* defines a positively invariant set of (36).

*Proof:* We apply Theorem 1 in [50], which states that for a parabolic PDE system with sufficiently smooth coefficients, a closed convex subset of euclidean space is positively invariant if the vector field corresponding to the “reaction dynamics” never points outwards at the boundaries of the set. To apply this theorem we first make a coordinate transformation. The spatial differential operator (as in Definition 1) for a general diffusing species *y*(*t, x*) with available volume profile *v*(*x*), given by

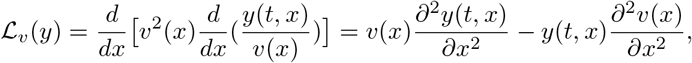

is not in the standard form stipulated by Equation 1.2 in [50]. Therefore, for Cases I-III, the following coordinate transformation is made

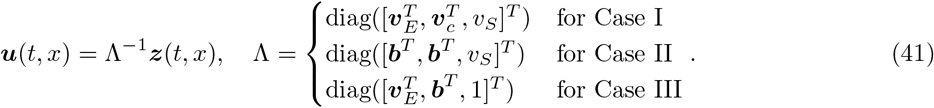

With the transformation *u*(*t, x*) = *y*(*t, x*)/*v*(*x*), the differential operator 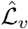 given by

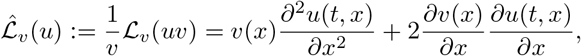

is in the postulated form. Let

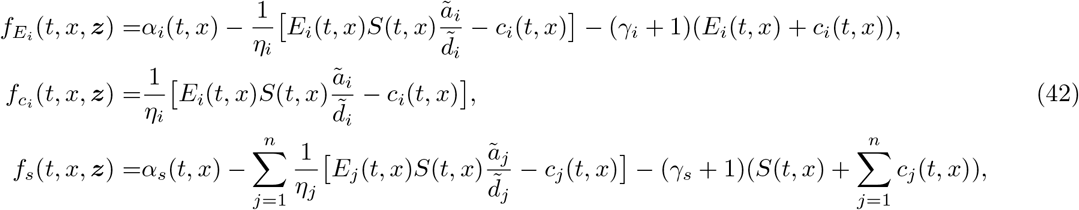

and 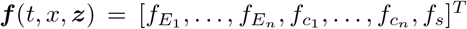 is the vector field of the reaction dynamics of (36). The vector field corresponding to the reaction dynamics of the transformed system is given by 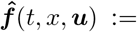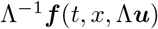. Let

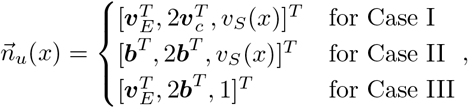

and 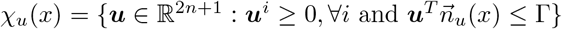. We now check that the vector field 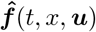 does not point outward at all the boundary points of *χ_u_*(*x*) ∀*x* ∈ [0, 1]. Checking that 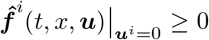, ∀*u* ∈ *χ_u_*(*x*), ∀*x* ∈ [0, 1] ∀*t* ≥ 0 is equivalent to checking 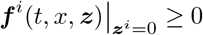, thus notice that for *i* = 1,…,*n*

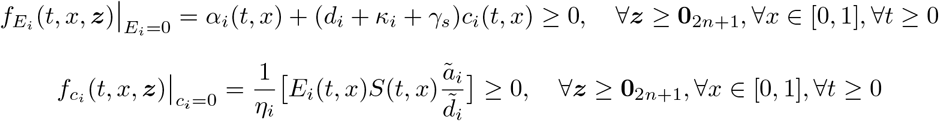

and

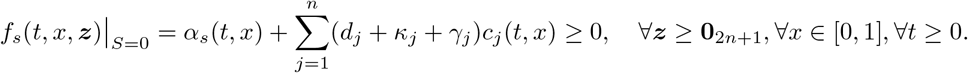

The set 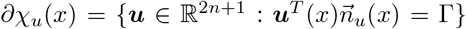, corresponds to the boundary points defined by planar surface 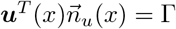 with normal vector 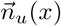, we need to check that for all boundary point 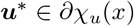 we have that 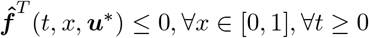:

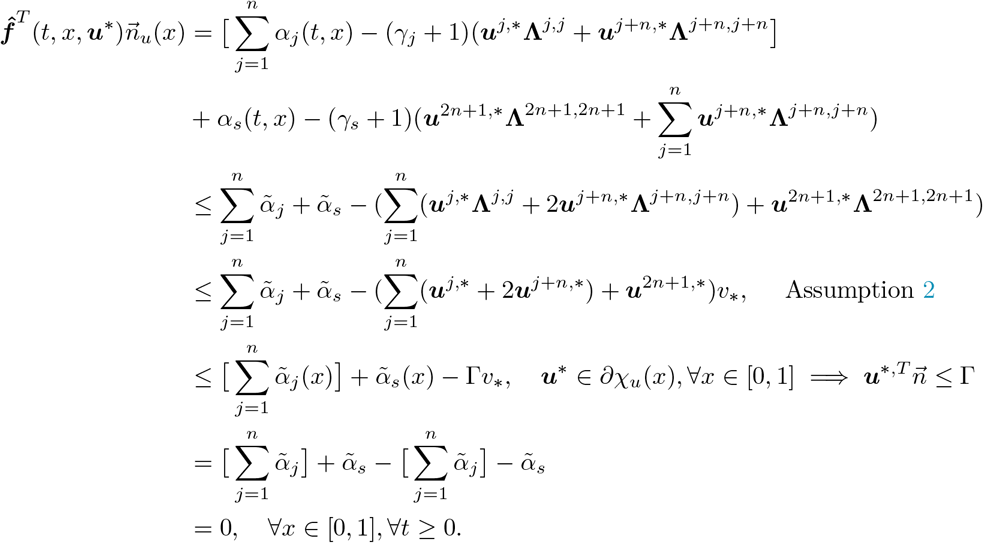

Thus, *χ_u_*(*x*) is a positively invariant set in the ***u***(*t, x*) coordinates. The corresponding invariant set in the ***z***(*t, x*) coordinates is given by *χ*.

##### Corollary 1

*The positivity of E_i_* (*t, x*), *S*(*t, x*), and *c_i_*(*t, x*) *imply the positivity of* 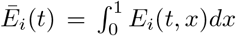, 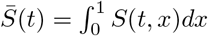, and 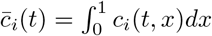.

##### Definition 4

Consider the systems given by (36) and (37) and let Cases I-III correspond to those in Table 2. We define

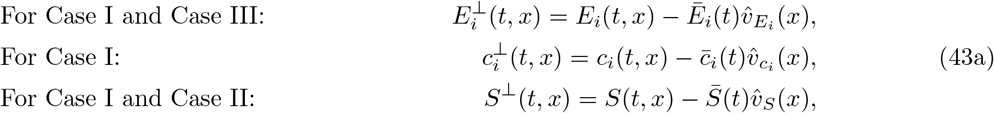

and

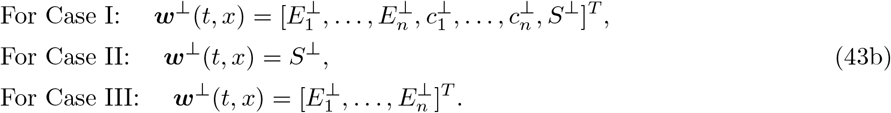

##### Lemma 4

*Consider the systems given by* (36) and let *ϵ* > 0 *be defined for Cases I-III by* Table 2. *Let **w***^⊥^(*t, x*) *be as in Definition 4 and suppose Assumption 2 holds. Let χ* ⊂ ℝ^2*n*+1^ *be as described in Claim 1 and let **z***(*t, x*) *be given by* (39) *Then there exists ζ*, *L*_⊥_ > 0 *such that for all **z***(0, *x*) ∈ *χ*, ∀*x* ∈ [0, 1] and ϵ > 0 ***w***^⊥^(*t, x*) *satisfies*

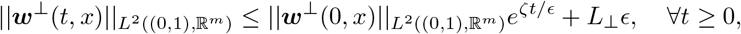

*where m* = 2*n* + 1 *in Case I, m* = 1 *in Case II, and m* = *n in Case III*.

*Proof:* This results follows directly from Theorem 2 where Assumption 1-I is satisfied by *χ* and Assumption 1-II is satisfied by the smoothness of the reaction dynamics in (36), the compactness of the sets *χ* and [0, 1], and the temporal boundedness (Assumption 2).

##### Remark 6

From the proof of Theorem 2, one can observe that *L*_⊥_ depends on the size of *χ*. Once Assumption 3 is made, the size of *χ* will depend on *δ* for Case II-III. Thus, *L*_⊥_ depends on *δ* for Cases II-III.

Next we define the space averaged total enzyme and substrate quantities and show that these are the same for (37) and (38). Furthermore, we will show the dynamics for these quantities are governed by uncoupled, linear, and contracting ODEs.

##### Definition 5

(Total space average enzyme and substrate concentrations) For (37), we define the total space averaged enzyme and substrate for *i* = 1. … *n* as 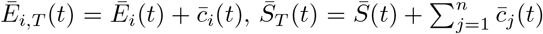, respectively, and similarly for (38), we define 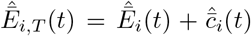 and 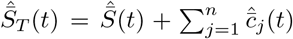, the dynamics of these quantities are given by

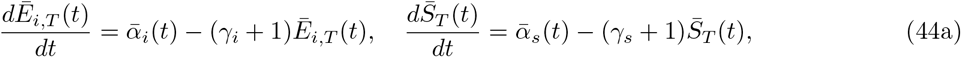

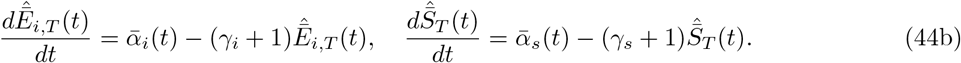

##### Remark 7

For *i* = 1, …, *n*, 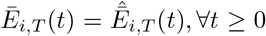 and 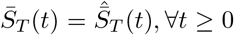 since from Definition 3, we have that 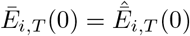 and 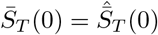.

##### Remark 8

From the linear and uncoupled structure of (44), it is clear that the dynamics for 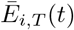 are contracting with contraction rate λ_i_ = *γ_i_*+1 for all *i* = 1,…, *n*. Similarly, the 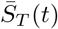 dynamics are contracting with contraction rate λ_*s*_ = *γ_s_* + 1.

##### Claim 2

Consider the systems (37), (38), and (44). Let 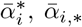, and 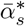 be as in Assumption 3. Assume that 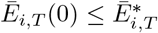 and that 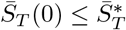. Then

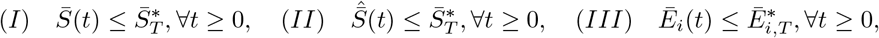

where 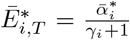 and 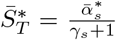. Furthermore, if we assume that 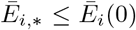, where 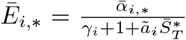, then we have that

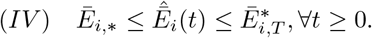

*Proof:* Let 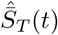 be given by (44), and let 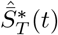 be given by

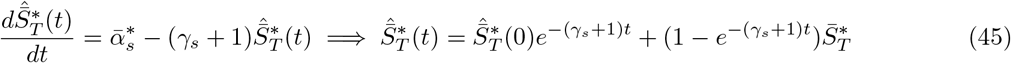

where 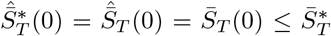, which implies that 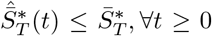. Let 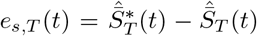 such that

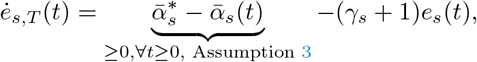

which implies that ℝ_+_ is a positively invariant set for the *e_s,T_* (*t*) dynamics. Since *e_s,T_* (0) = 0, this implies that *e_s,T_* (*t*) ≥ 0, ∀*t* ≥ 0 and thus 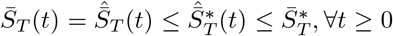 (first equality from Remark 7). From the positivity of 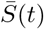 and 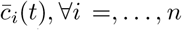 (Corollary 1), we have that 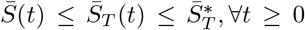, thus proving (I). Similarly, 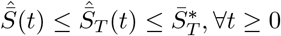, thus proving (II).

Let 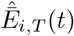 be given by (44) and 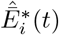 be given by

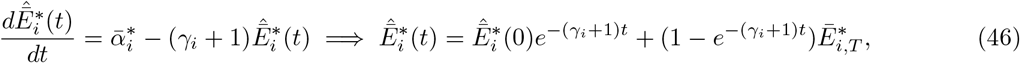

where 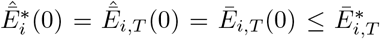, which implies that 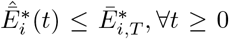. Let 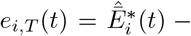 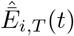 and thus

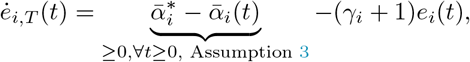

which implies that ℝ_+_ is a positively invariant set for the *e_i,T_* (*t*) dynamics. Since *e_i,T_* (0) = 0, this implies that *e_i,T_* (*t*) ≥ 0, ∀*t* ≥ 0 and thus 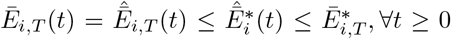 (first equality from Remark 7). From the positivity of 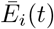 and 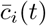 (Corollary 1), we have that 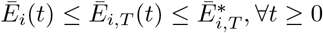, thus proving (III). By similar logic, we also have that 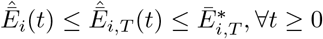, thus proving part of (IV).

Let 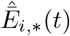 such that 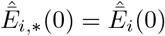 and

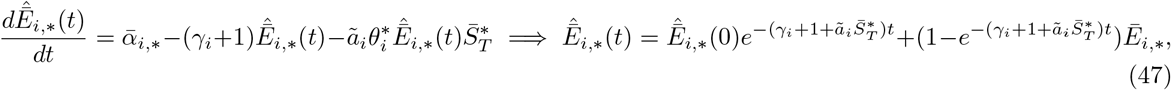

where 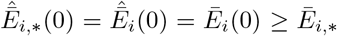, which implies that 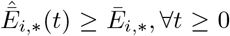. Let 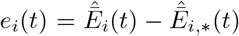 such that

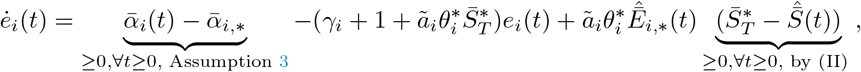

which implies that ℝ_+_ is a positively invariant set for the *e_i_*(*t*) dynamics. Since *e_i_*(0) = 0, this implies that *e_i_*(*t*) ≥ 0 ∀*t* ≥ 0 and thus 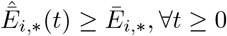. Thus proving (IV).

##### Remark 9

By Assumption 3, we have that 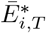 and 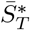 are independent of *δ* and thus these upper bounds for 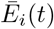 and 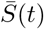, respectively, are also independent *δ*.

The following results demonstrates that Assumption 3 implies that the concentration of fixed species is localized at the region specified in Table 2. The parameter *δ* > 0, controls the amount of localization.

##### Proposition 1

Consider the systems given by (36). Let 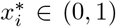 for Case II and 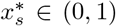 for Case III, be given by Table 2. Suppose that Assumption 3 holds for 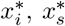, and a given *δ* > 0. Let *E_i,T_*(*t, x*) = *E_i_*(*t, x*) + *c_i_*(*t, x*) and 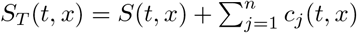. Then, for all 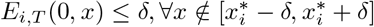 for Case II, and 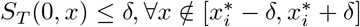 for Case III, we have that

1. Case II: 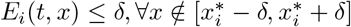, ∀*t* ≥ 0 ∀*i* = 1,…,*n*,
2. Case III: 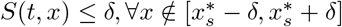, ∀*t* ≥ 0,

*Proof:* For Case II, *E_i,T_* (*t, x*) satisfies

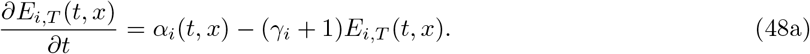

For 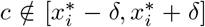 we have that 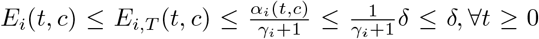. Similarly, for Case III, *S_T_* (*t, x*) satisfies

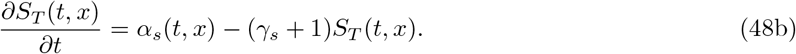

For 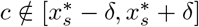 we have that 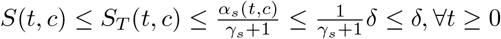.

The following claim will aid us in rewriting (37) in the form of the reduced dynamics given by (38) with additional “disturbance” terms of order *ϵ* and *δ* The claim is written to handle Case I-III.

##### Claim 3

For a given *y*_1_(*t, x*) ∈ *H* and *y*_2_(*t, x*) ∈ *H* where *H* = *L*_2_(0, 1), suppose that *y*_1_(*t, x*), *y*_2_(*t, x*) ≥ 0 ∀*t* ≥ 0 ∀*x* ∈ [0, 1] and that there exists 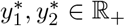 such that 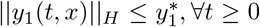 and 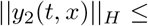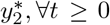, let 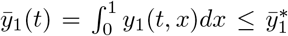 and 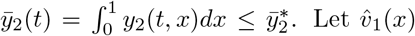. Let 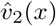 be smooth positive functions and denote 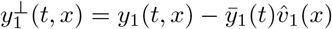 and 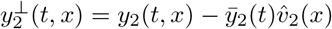.

1. There exists constant *k*_1_ *k*_2_ > 0 such that

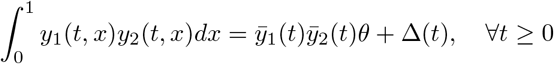

where 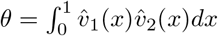 and 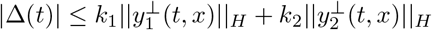.
2. Suppose that for a given *x*\* ∈ (0, 1) and *δ* > 0 such that [*x*\* − *δ*, *x*\* + *δ*] ⊂ [0, 1] we have that *y*_2_(*t, x*) ≥ *δ*, ∀*x* ∉ [*x*\* − *δ*, *x*\* + *δ*]. Furthermore assume that 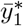 and 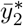 are independent of *δ* Then there exists constant *k*_3_(*δ*) *k*_4_ > 0 such that

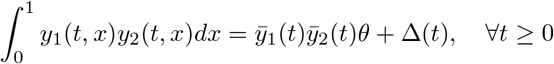

where 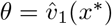 and 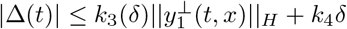.

*Proof:* To proof the first claim, notice that

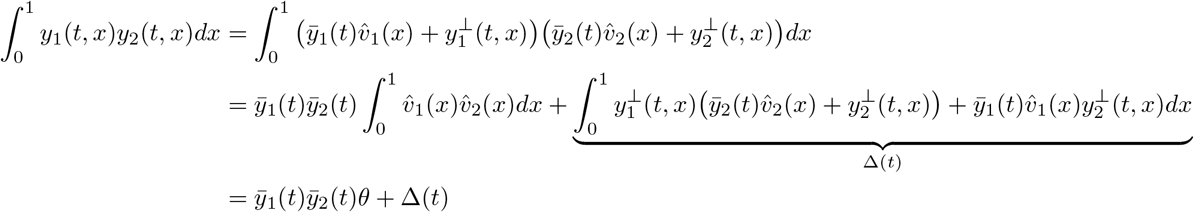

where 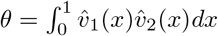. Let 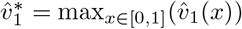 and leveraging the Cauchy-Schwarz inequality in *H*,

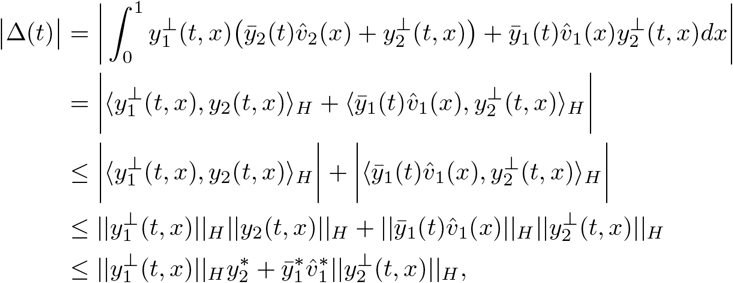

thus 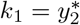 and 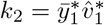. For the second part of the claim, notice that

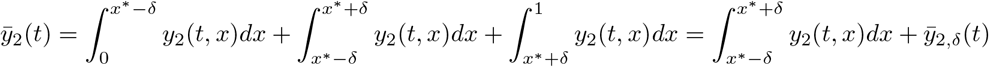

where

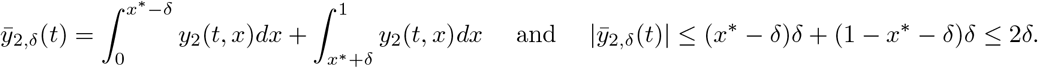

Next,

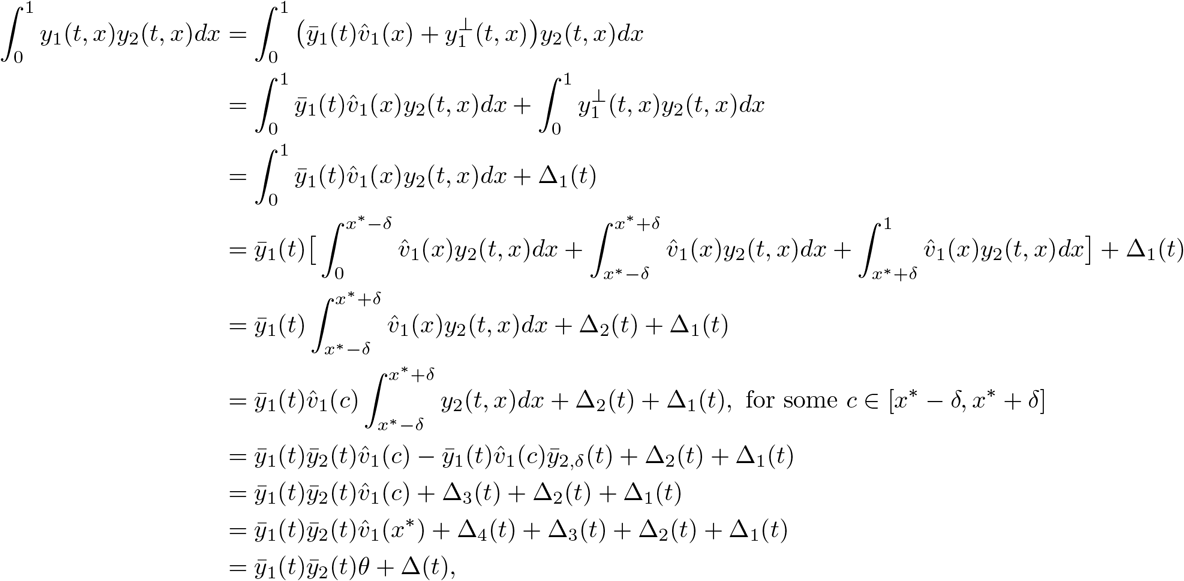

where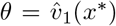 and 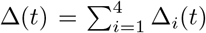, the existence of *c* is guaranteed by the mean-value theorem for integrals [51]. Notice that

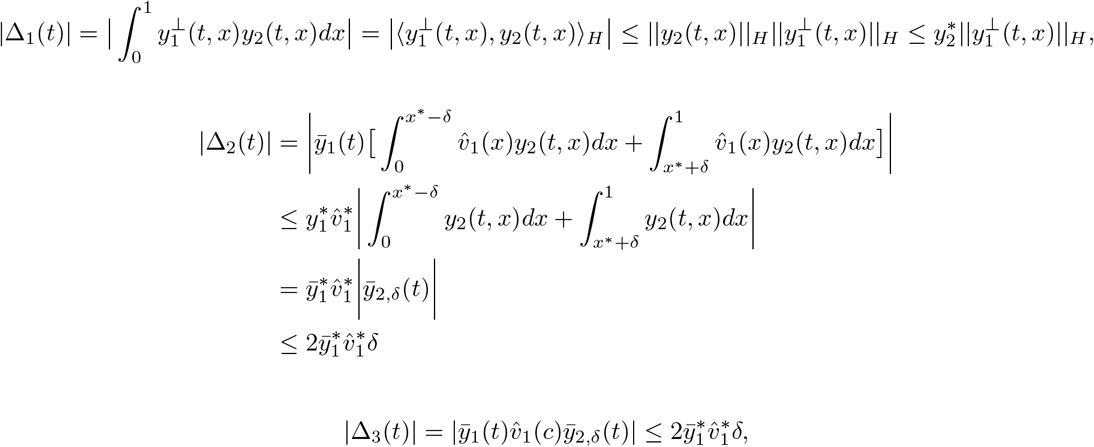

the following uses the smoothness 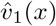 to guarantee uniform continuity i.e., the existence of *L_v_* > 0 such that for all *x*_1_, *x*_2_ ∈ (0, 1), we have that 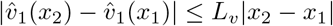, and hence

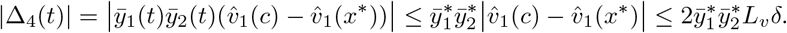

Finally,

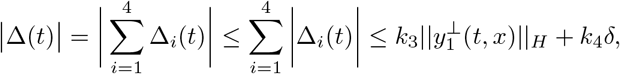

where 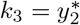 and 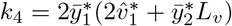.

##### Remark 10

In the proof of Claim 3-2, 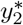 may depend on *δ* since we assume that 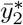 is independent of *δ* (one expects that 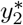 increases with decreasing *δ*), thus *k*_3_ may depend on *δ*.

##### Corollary 2.

*Consider the systems given by* (36). *The assumptions necessary to apply Claim 3 to*

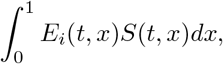

*are satisfied by Claim 1, Proposition 1, and Claim 2 (along with Remark 9). Furthermore, considering the results from Lemma 4, we are guaranteed he existence of L_i_* > 0 *for i* = 1, …, 5 *such that for all ϵ* > 0

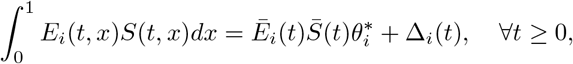

where 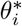 *is given by* (38b) *for Cases I-III (as in Table 2), and*

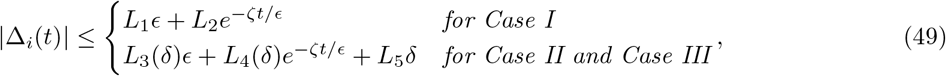

*where ζ is as in Lemma 4. The coefficients L*_3_ *and L*_4_ *may depend on δ by the discussion in Remark 6 and Remark 10*.

Let 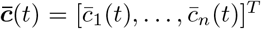 where 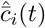 is given by (37) 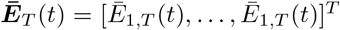 where 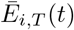 is given by (44), 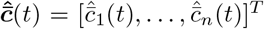, where 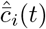 is given by (38), the 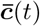 and 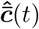 dynamics may be written as

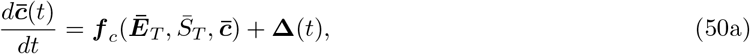

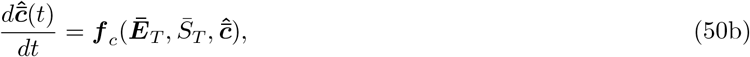

where **Δ**^*i*^ = Δ_*i*_(*t*) as in Corollary 2 and ***f**_c_*: ℝ*n* × ℝ × ℝ*n* → ℝ is given by

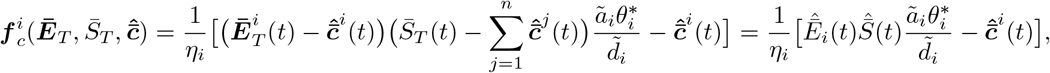

we used the fact that 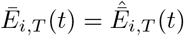 and 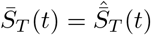 (Remark 7). By the form of (50), it is clear that the 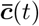 dynamics (50a) are in the form the 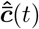 (50b) with additional “perturbation” terms of order *ϵ* and *δ*. The variables 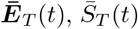, and 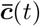 are enough to fully describe (37) and 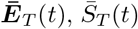, and 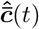 are enough to fully describe (38). Notice that

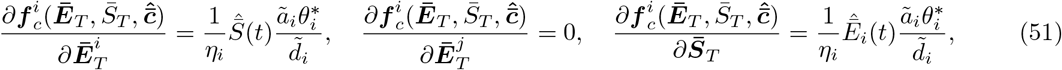

and by Claim 2 these terms are uniformly bounded in time and for *i* = 1,…, *n* Considering Lemma 2 with 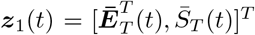 and 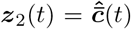, then condition (I) is satisfied by the discussion in Remark 8 and condition (III) is satisfied by (51), thus we can treat 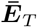 and 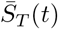 as time varying inputs to (50b) when showing that (II) is satisfied.

We now show that the dynamics (50b) are contracting and thus we can apply the robustness property of contracting systems (Lemma 3) to show that the solutions of (50b) and (50a) are close.

##### Lemma 5

*Consider the system* (36) *and let ϵ* > 0 *be defined for Case I-III by Table 2. Let* 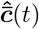 *be given by* (50b) *and* 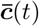 *be given by* (50a) *Suppose the conditions of Claim 1, Lemma 4, and Claim 2 hold. Then, there exists L*_*c*,1_ *ϵ*\* > 0, *such that for all ϵ* ≤ *ϵ*\* *the solutions* 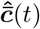 *and* 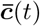 *satisfy*

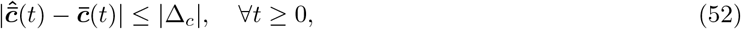

*where* |Δ_*c*_| = *L*_*c*,1_*ϵ for Case I. For Cases II-III, if in addition, the conditions of Proposition 1 hold for all* 0 ≤ *δ* ≤ *δ*\*, *there exists L*_*c*,3_ > 0 *such that for all* 0 < *δ* < *δ*\* *there exists L*_*c*,2_(*δ*) *such that* (52) *is satisfied with* |Δ_*c*_| = *L*_*c*,2_(*δ*)*ϵ* + *L*_*c*,3_*δ*.

*Proof:* Consider the metric

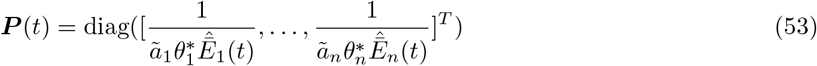

where 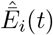 is given by (38). The total time derivative of ***P*** is given by

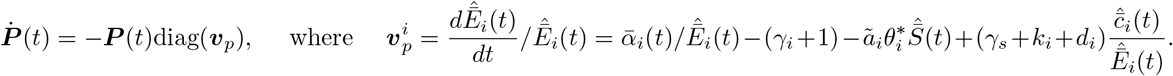

The Jacobian of (50b) is given by

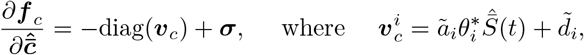

and ***σ*** is a rank one matrix given by

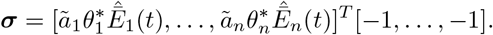

With the chosen metric ****P**** (*t*), we have that

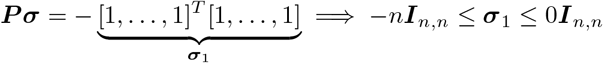

since the symmetric rank one matrix ****σ****_1_, has *n* − 1 zero eigenvalues and the nontrivial eigenvalue is *λ_σ_* = −*n* for eigenvector *v_n_* = [1,…, 1]^*T*^. Recalling the positivity of 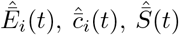 (Claim 1), and 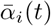, we have that

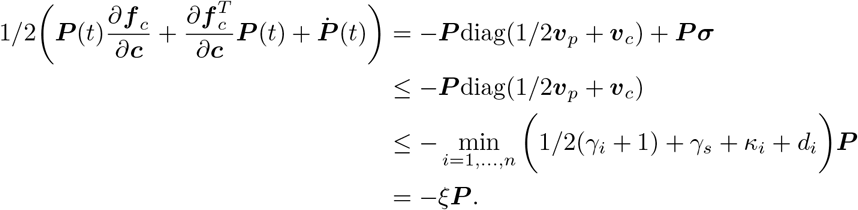

By Theorem 1, the system (50b) is contracting with contraction rate *ξ* = 1/2 +*γ_s_* + min_*i*=1,…,*n*_ (1/2*γ_i_* +*κ_i_* + *d_i_*). Assumptions 2, 3 imply that (49) holds for Δ_*i*_(*t*) in (50a). Therefore, for a given *δ* > 0, we apply the result from Lemma 3 to the nominal system (50b) and the perturbed system (50a). Let *ϵ*\* = *ζ*/(2*ξ*) and recalling that 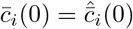, then by Lemma 3, there exists *l*_1_, *l*_2_(*δ*) *l*_3_ > 0 such that for all *ϵ* < *ϵ*\*

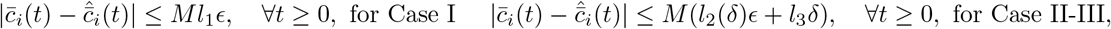

where *M* is a constant upper bound on the square root of the ratio of the biggest and smallest eigenvalues of ***P*** (*t*). Thus *L*_*c*,1_ = *Ml*_1_, *L*_*c*,2_(*δ*) = *Ml*_2_(*δ*), and *L*_*c*,3_ = *Ml*_3_. We now show that *M* exists. Let *r*(*t*) be the ratio of the biggest and smallest eigenvalues of ***P***(*t*). By Claim 2, there exists 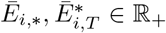 independent of *E* and *δ* such that 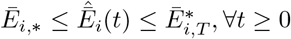 and thus

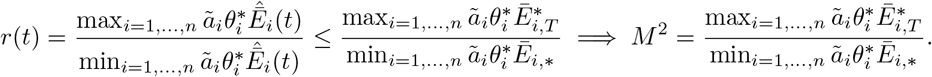

##### Corollary 3

*Recall Definition 5 and Remark 7, we have that*

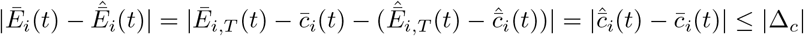

*and*

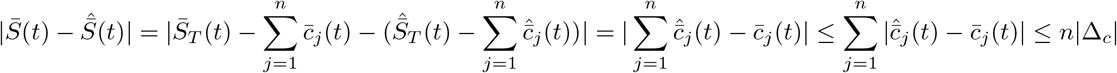

*where* |Δ_*c*_| *as in Lemma 5. Thus, the quantity* |Δ_*z*_| *as claimed to exist in Theorem 3, may be given by* 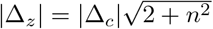.

##### Remark 11

We now comment on the sets Ω_*z*_ ∈ ℝ^*n*+1^ and 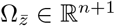 as claimed to exist in Theorem 3. Let *χ* be as in Claim 1.

- For Cases I-III we require that ***z***(0, *x*) *χ*, ∀*x* [0, 1] for Lemma 4 to hold. In Case I, Ω_*z*_ ∈ ℝ^*n*+1^ = *χ*. In Case II, for Proposition 1 to hold we also require that for *i* = 1… *n*, that 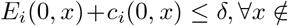 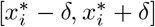 thus Ω_*z*_ is the intersection between the set that satisfies this conditions. and *χ*. In Case III, for Proposition 1 to hold we also require that 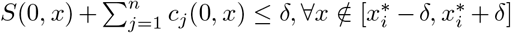 thus Ω_*z*_ is the intersection between the set that satisfies this condition and *χ*. As discussed in Remark 6, *χ* may depend on *δ*.
- For Claim 2 to hold, we assumed that 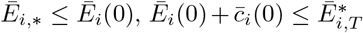, and that 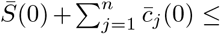 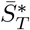 thus 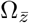 is the set that satisfies these conditions.

### 2.4 Fast Diffusion and Bingding Dynamics

The approximation result of Theorem 3, holds well if ***w***^⊥^ is small, where ***w***^⊥^ is given by Definition 4. This was guaranteed by Lemma 4 which is based on Theorem 2. The proof of Theorem 2 was based on the principle that the 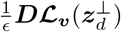 term dominates the 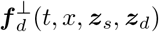 term in (35c) and thus all solutions converged to 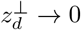, the quasi-steady state of 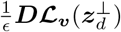. However, for (36), if *ϵ* and *η_i_* are of similar order of magnitude, the corresponding term in 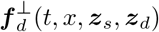 may be comparable to 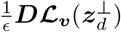 and cannot be neglected. The terms *ϵ* and *η_i_* being comparable corresponds to diffusion and the binding between E_i_ and S occurring at similar timescales, which often time occurs within the cell [2]. Here we show that when both diffusion and the binding dynamics dominate in (36), 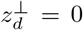 is still the quasi-steady state, that is, all freely diffusing species mirror their available volume profile.

When both diffusion and the binding dynamics dominate in (36) we have that

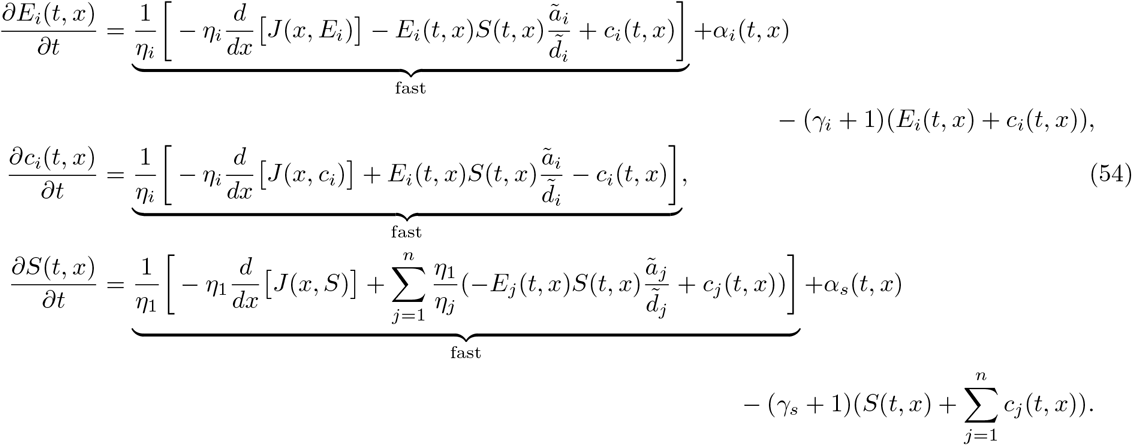

We compute the quasi-steady state of the “fast dynamics in (54) for Cases I-III, that is *E_i_*(*t, x*), *c_i_*(*t, x*), and *S*(*t, x*) such that

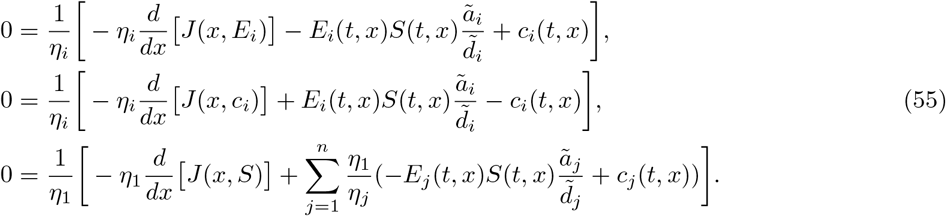

**Case I**: In this case, (55) is satisfied for

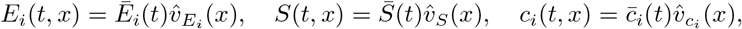

with the additional constraint that

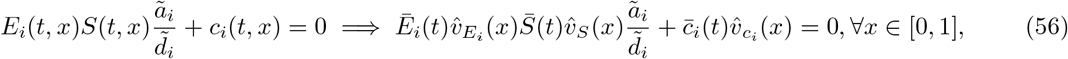

which in general (56) is a stringent condition since it is required to hold for all *x* ∈ [0, 1]. However, by the key fact that 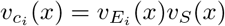 ((9) in the main text), (56) is satisfied for all *x* ∈ [0, 1] by

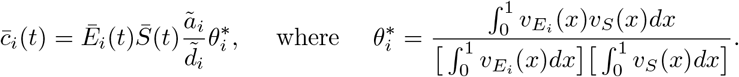

**Case II and III**: For this cases, (55) is satisfied for

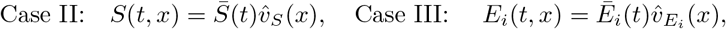

with the additional constraint that 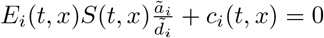 which implies that

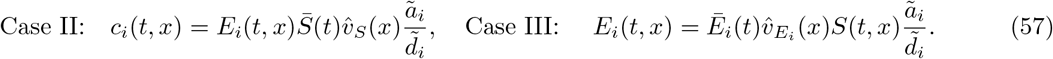

Thus, when both diffusion and the binding dynamics dominate, the quasi-steady states are still those that correspond to freely diffusing species converging to their available volume profile. We observed that for Case I, this was possible by the fact that 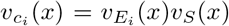. In a future study it should be shown that this quasi-steady state is a stable solution of the fast dynamics in (54).

### 2.5 Available Volume Profiles and Bounds on 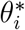

Following [13], we introduce a model of the available volume profiles of a freely diffusing species within the DNA mesh of the cell. Let *ρ*(*x*) be the local density of DNA length such that 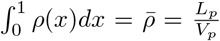 where *L_p_* is the total length of chromosome DNA, *V_p_* the volume where the DNA polymer is confined, and let 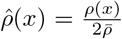. For a species diffusing inside the cell with radius of gyration *r*, we model the available volume profile *v*(*x*) as:

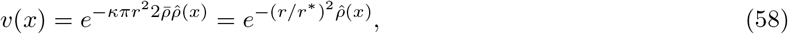

where *κ* is an empirical coefficient (as discussed in [13]) and 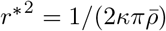. From the parameter values in [13], *r*\* ≈ 23 nm. As commented in SI Section 2.11, *r*\* can be estimated for a given context (e.g., growth conditions and strain) by analyzing the concentration profile inside the cell of a freely diffusing species with a known radius of gyration, which is possible via superresolution imaging [14].

As shown in Figure 8, we estimate the chromosome density as a step function.

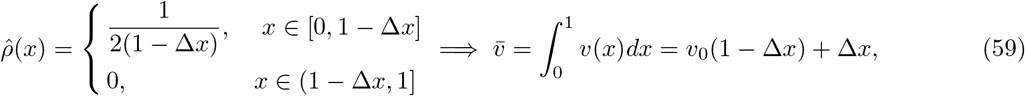

where 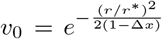 and Δ*x* is the distance between the end of the chromosome and the cell poles (see Figure 1 in the main text). Its clear now, that our choice to define 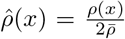 was motivated by the fact that when Δ_*x*_ = 1/2 (nucleoid evenly spread out between mid-cell and the halfway point between mid-cell and the cell poles), we have the convenient expressions 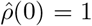 and 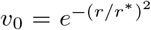. Notice that *v_0_* → 0 as (*r*/*r*\*)^2^ → ∞. Thus,

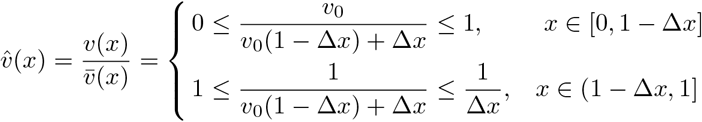

#### Bounds on 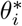 (13)

For the species E_i_ and S as described in the main text with radius of gyration *r_e,i_* and *r_s_* respectively. The available volume profiles are given by

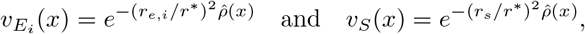

respectively, We summarize the bounds on 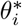 (13) in the main text, assuming *ρ*(*x*) is a step function as above. Let

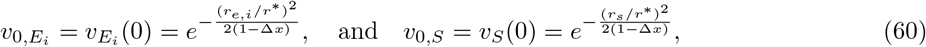

and Case I-III as in (13) in the main text, the bounds on 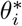 are given by

- For **Case I**, this idealization implies that

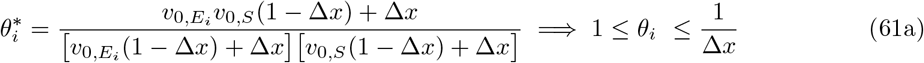 The upper limit of 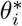 is 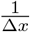 and is reached as 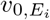 <u>and</u> *v*_0,*S*_ approach zero, which occurs as *r_e,i_*/*r*\* → ∞ and *r_s_*/*r*\* → ∞. The lower limit of 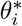 is unity and is achieved if 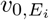 <u>or</u> *v_0, S_* approach one, which occurs if any of the two species is sufficiently small (*r_e,i_*/*r*\* ≪ 1 or *r_s_/r*\* ≪ 1). Since 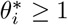 it implies that the binding betwen E_i_ and S is always equal to or greater than that predicted by a well-mixed model ((8) in the main text).
- For **Case II-III** let 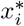 and 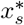 as in Assumption 3 in the main text and thus

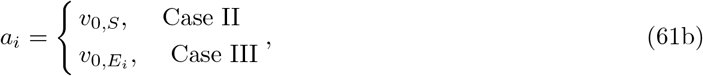

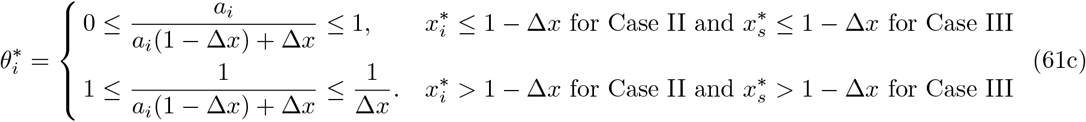 When 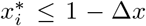 for Case II and 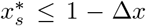 for Case III, we have that 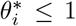, the lower limit 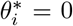is achieved when *r_e_*/*r*\* → ∞ for Case II (*r_s_/r*\* → ∞ for Case III), the upper limit 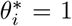 is achieved when *r_e_*/*r*\* → 0 for Case II (*r_s_/r*\* → 0 for Case III). When 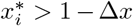 for Case II and 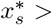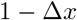 for Case III, we have that 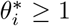, the lower limit 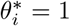 achieved when *r_e_*/*r*\* → 0 for Case is achieved when II (*r_s_/r*\* → ∞ for Case III), the upper limit 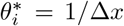 (*r_s_/r*\* → ∞ for Case III).

**Figure 8:**
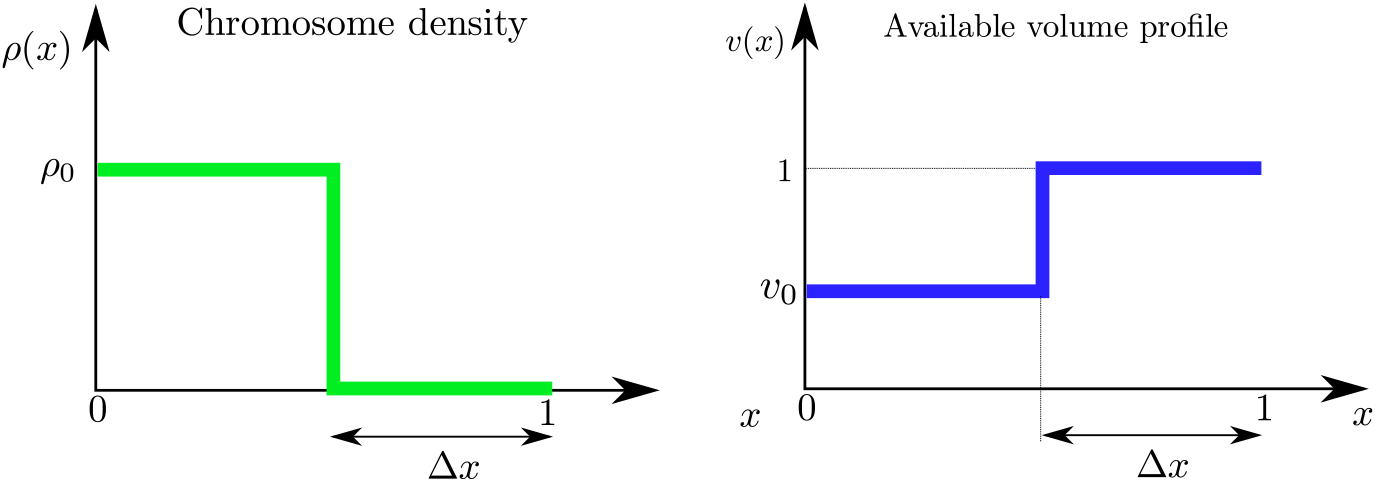
Idealization of the chromosome density which yields a simple estimate of the available volume profile. The chromosome density is approximated as a step function implying by (58) that the available volume profile is also a step function. Here Δ*x* is the distance between the end of the chromosome and the cell poles.

### 2.6 Protein Production: Transcription and Translation

We consider gene (*D*) being transcribed by RNAP (*S*) to form a DNA-RNAP complex (*c_s_*) to produce mRNA (*m*) which is translated by ribosomes (*R*) to form mRNA-ribosome complex (*c_m_*) which produces protein *P*. The mRNA’s degrade at rate *γ*. The RNAP, and ribosomes are produced at rates *α_s_*(*t, x*), *α_r_*(*t, x*), respectively. We assume all species dilute at rate *μ* the cells growth rate. The corresponding biochemical reactions are:

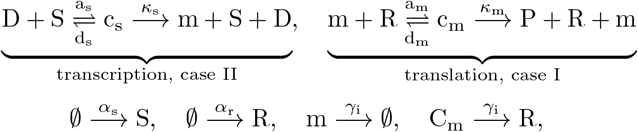

where *γ* is the mRNA degradation rate, *a_s_* and *d_s_* are the association and dissociation rate constants, respectively, between RNAP and the gene D, *κ_s_* is the catalytic rate of formation of mRNA m, *a_m_* and *d_m_* are the association and dissociation rate constants, respectively, between ribosomes and mRNA *κ_m_* is the catalytic rate of formation of protein P. We assume that the total concentration of D is conserved, so that *D_T_* (*x*) = *D*(*t, x*) + *c_s_*(*t, x*) and that *D_T_* (*x*) is localized at *x*= *x*\*.

#### Spatial-temporal Dynamics

The dynamics corresponding to these biochemical reactions are given by:

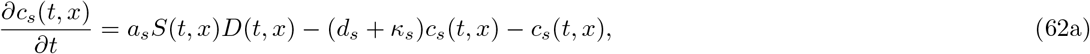

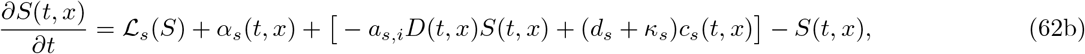

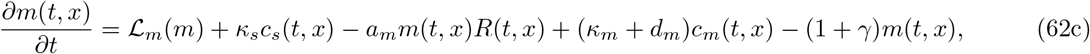

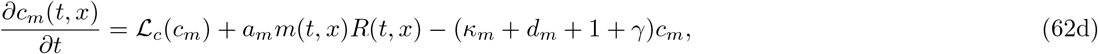

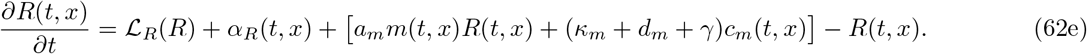

where the spatial variable has been normalized by *L* (cell-length) and the time variable has been normalized by 1/*μ* the time scale associated with dilution. The flux dynamics and boundary conditions are given by,

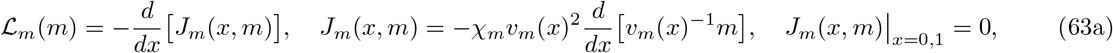

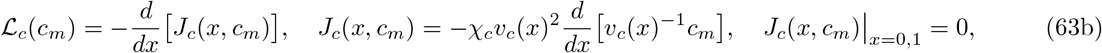

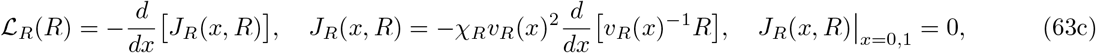

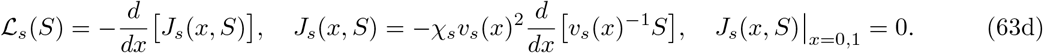

where *v_m_*(*x*), *v_c_*(*x*), *v_R_*(*x*), and *v_s_*(*x*) are the available volume profiles for the mRNA, mRNA-ribosome complex, ribosome, and RNAP, respectively and *χ_m_* = *D_m_*/(*L*^2^*μ*), *χ_c_* = *D_c_*/(*L*^2^*μ*), *χ_R_* = *D_R_*/(*L*^2^*μ*) and *χ_s_* = *D_s_*/(*L*^2^*μ*), are the dimensionless diffiusion coefficients for the mRNA, mRNA-ribosome complex, ribosome, and RNAP, respectively. The space averaged protein concentration 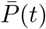 is given by

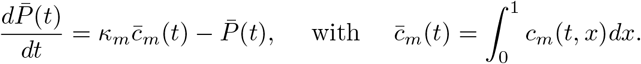

#### Values for dimensionless parameters

We set all production rates with respect to that of RNAP such that 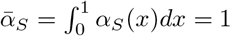. All time scales relative to *μ* = 0.5 1/hr, consistent with the experiments [52] The total number of RNAP (*N*_RNAP_) ranges between 2,000 −10,000 we took it to be 5,000 [14]. The total number of ribosomes (*N*_ribo_) was taken to be 10,000 and since both RNAP and ribosomes and RNAP are stable, it implies 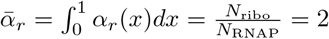.mRNA degradation is about 10 times faster than dilution [2], therefore, *γ* = 10.The rate of transcription (translation) is about 80 (40) times faster than dilution [2], thus we choose *κ_s_*= 80 and *κ_m_* = 40. We assumed that the DNA is on a high copy plasmid (≈ 500 copies) and thus 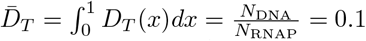.

The association and dissociation rate constants are varied as shown below to show that all of our results hold dispiite fast binding and unbinding but we maintain the ratio *d_s_*/*a_s_* = *d_m_*/*a_m_* = 1.

The length of the cell is about 3*μ*m and thus *L* = 1.5*μ*m [13]. The diffusion coefficient of RNAP is taken to be *D_s_* = 0.22*μm*^2^/*s* [53] and thus *χ_s_* = 704. The diffusion coefficient of free ribosomes is taken to be *D_r_* = 0.4*μm*^2^/*s* [13] and thus *χ_r_* = 1280. In [14], the diffusion coefficient of polysomes is 0.05 ± 0.02*μm*^2^/*s* and thus we take the diffusion coefficient of a free mRNA to be the upper bound 0.07*μm*^2^/*s* and thus *χ_m_* = *χ_c_* = 224.

For the following, the spatial profiles for the production rates are given proportional to their functional form since the constant that fully specifies them is such that the production rate per-cell satisfies the above values.

#### Additional simulation details for Figure 5-A in the main text

*D_T_*(*x*) ∝ *e*^−20*x*^ when DNA near mid cell and *D_T_*(*x*) ∝ *e*^20(*x*−1)^ when DNA at cell poles. The RNAP production was kept roughly spatially constant *α_s_*(*x*) ∝ *e*^−.001*x*^. The binding and unbinding coefficients for DNA-RNAP were *a_s_* = 1000 and *d_s_* = 1000. We set *a_m_* = *d_m_* = 0 such that mRNA did not bind to ribosomes and thus the free amount of mRNA is equivalent to the total mRNA.

#### Additional simulation details for Figure 5-B in the main text

*D_T_*(*x*) ∝ *e*^−.001*x*^ is chosen to be roughly constant. The RNAP production was kept roughly spatially constant *α_s_*(*x*) ∝ *e*^−.001*x*^. The ribosome production was kept roughly spatially constant *α_r_*(*x*) ∝ *e*^−.001*x*>^. The RNAP radius of gyration was taken to be *r_s_*/*r*\* = 0.001 such that its excluded volume effects were negligible. The binding and unbinding coefficients for DNA-RNAP were *a_s_* = 1000 and *d_s_*= 1000. The binding and unbinding coefficients for ribosome-mRNA were *a_m_* = 10 and *d_m_* = 10.

**Figure 9:**
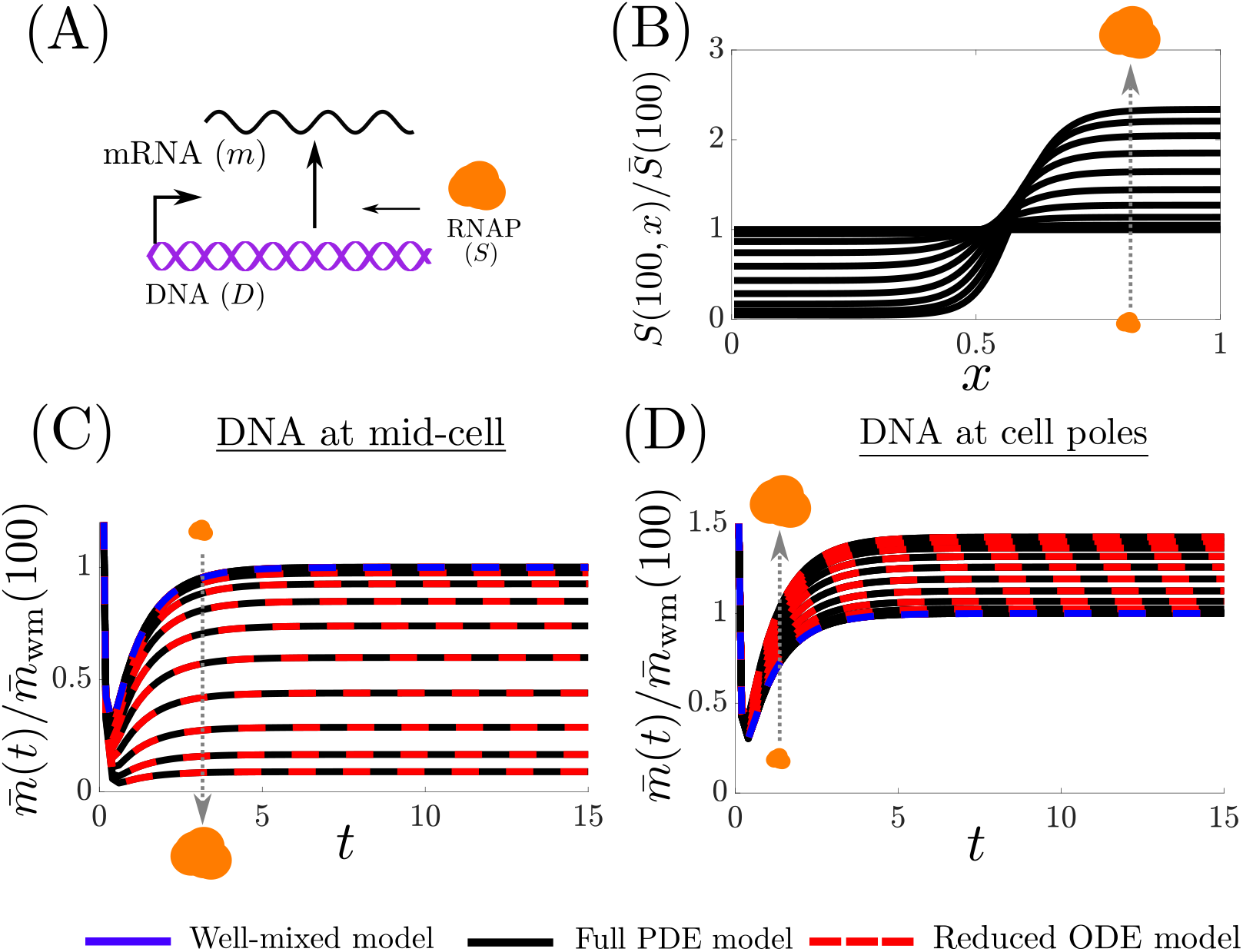
RNAP steady state spatial profiles and space averaged mRNA transients. For the following we refer to the well-mixed model as (15) with (16) given by *θ_s_* = 1 and *θ_r_* = 1. Here time is nondimensionalized with the time scale associated with dilution. (A) DNA transcribed by RNAP (S) to form mRNA (m) (B) The steady state RNAP spatial profile predicted by (62), normalized by spatial averaged value. From the results on the main text this should mirror the normalized available volume profile, which it does (Remark 1). Note as the size of RNAP increases, it is further excluded from the chromosome. (C) The temporal space-average concentration of mRNA when the DNA is localized mid-cell for several sizes of RNAP for the well-mixed model, reduced ODE model (15) and PDE (62). (D) The temporal space-average concentration of mRNA when the DNA is localized near the cell poles for several sizes of RNAP for the well-mixed model, reduced ODE model (15) and PDE (62). The simulation set up and parameters are identical to those of Figure 5-A.

### 2.7 Multiple Ribosomes on a Single Strand of mRNA

The biochemical reactions that models a polysome with *N_r_* bound ribosomes are given by

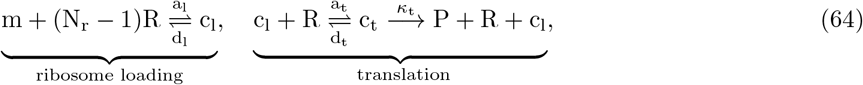

where the first and second reaction model the loading and translation steps, respectively. The translation dynamics corresponding to (64) are given by

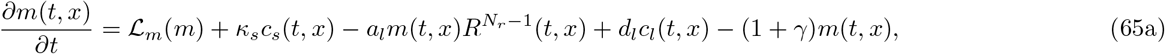

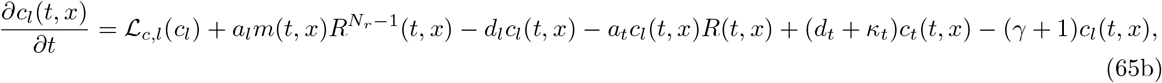

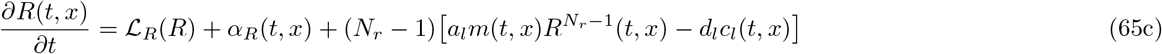

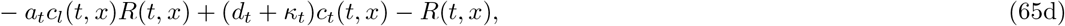

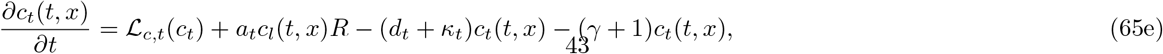

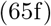

where *c_s_*(*t, x*) is given by (62), 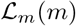 and 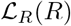 are given by (63) and

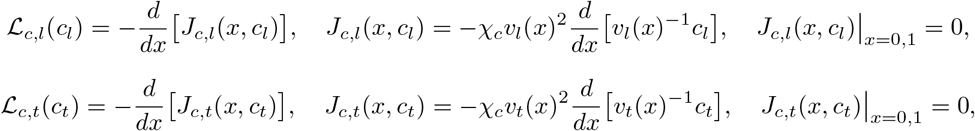

where 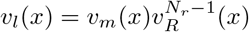 and *v_t_*(*x*) = *v_l_*(*x*)*v_R_*(*x*). Integrating (65) in space yields:

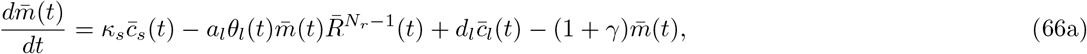

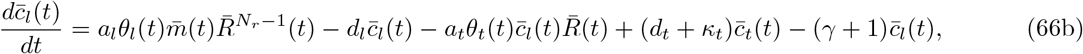

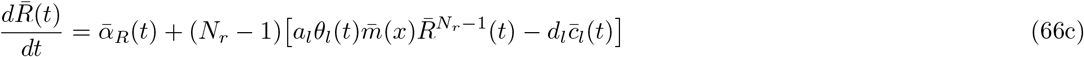

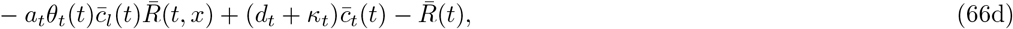

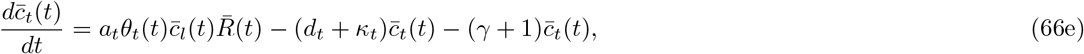

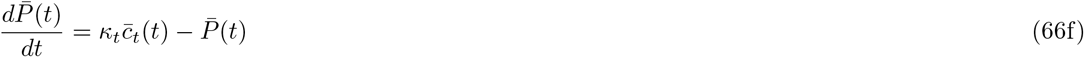

where

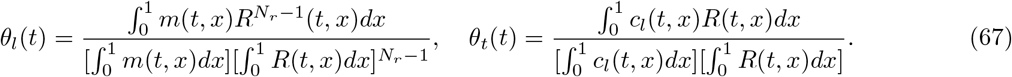

**Figure 10:**
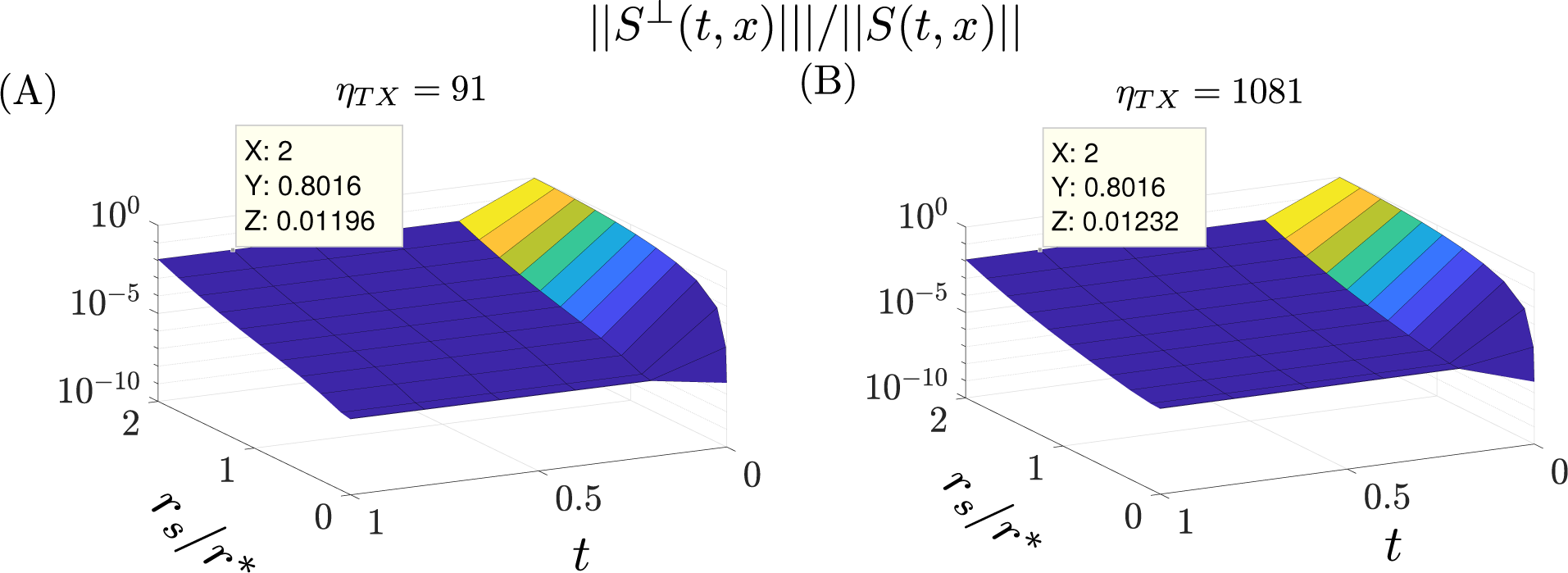
The error in the RNAP approximation for several binding and unbinding speeds between DNA and RNAP. Let *S*(*t, x*) be as in (62) and 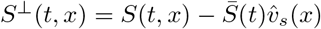 be the a measure of the error in our approximation, where 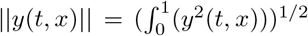, and *η*_TX_ = 1/(*κ_s_* + *d_s_* + 1). The values of *η*_TX_ are varied by modifying *d_s_* while maintaining *a_s_*/*d_s_* = 1. Here time is nondimensionalized with the time scale associated with dilution. (A) The relative error in time for several values of *r_s_* for *η*_TX_ = 91 =⇒ *d_s_* = 10. (B) The relative error in time for several values of *r_s_* for *η*_TX_ = 1081 =⇒ *d_s_* = 1000. For both values of *η*_TX_ the error is high at *t* = 0 since the initial RNAP spatial profile is chosen to be a constant but quickly decays to less than 2%. The rest of the simulation set up and parameters are identical to those of Figure 5-A.

**Figure 11:**
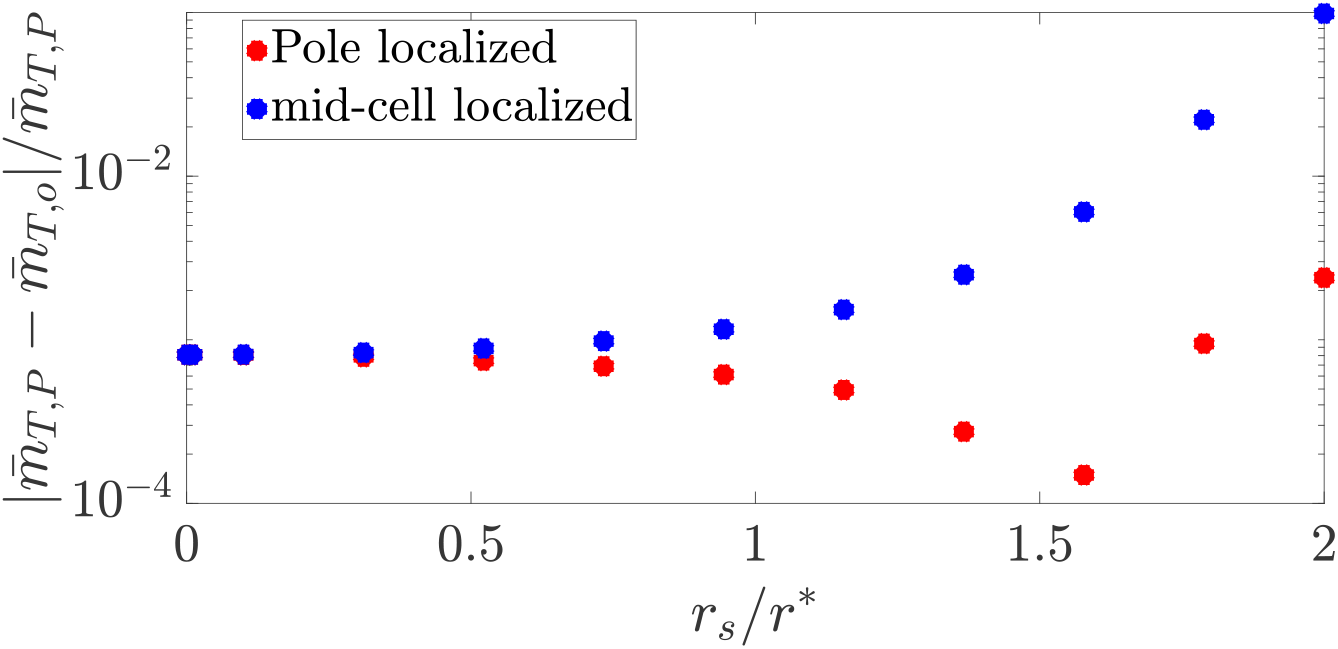
The relative error between the space averaged PDE model and the reduced ODE model from the data in Figure 5-a in the main text. The relative error in the steady state space averaged mRNA for the full PDE model 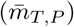 and the reduced model 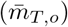 from the data in Figure 5-a in the main text. When the DNA is pole localized the relative error is less than 1% and when the DNA is localized near mid-cell the error is less than 10%.

**Figure 12:**
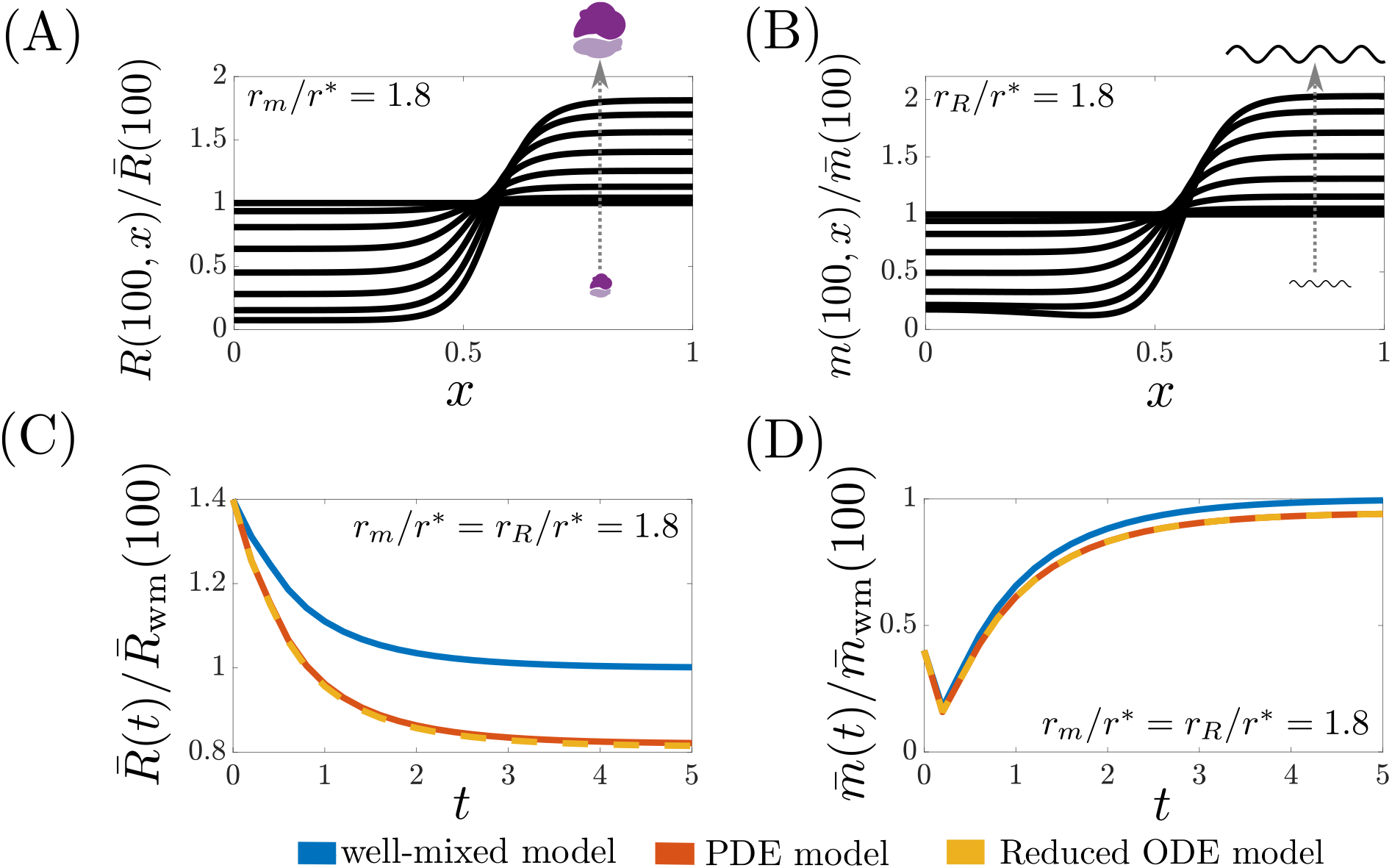
Ribosome and mRNA steady state spatial profiles and space averaged transients. For the following we refer to the well-mixed model as (15) with (16) given by *θ_s_* = 1 and *θ_r_* = 1. Here time is nondimensionalized with the time scale associated with dilution. (A) The steady state ribosome spatial profile predicted by (62), normalized by spatial averaged value. From the results on the main text this should mirror the normalized available volume profile, which it does. Note as the size of the ribosome increases, it is further excluded from the chromosome. (B) The steady state mRNA spatial profile predicted by (62), normalized by spatial averaged value. From the results on the main text this should mirror the normalized available volume profile, which it does. Note as the size of the mRNA increases, it is further excluded from the chromosome. (C) The temporal space averaged concentration of ribosomes normalized by the steady state of the well-mixed model for the reduced ODE model (15) and the PDE model (62) when *r_m_*/*r*\* = *r_R_*/*r*\* = 1.8. (D)The temporal space averaged concentration of mRNA normalized by the steady state of the well-mixed model for the reduced ODE model (15) and the PDE model (62) when *r_m_*/*r*\* = *r_R_*/*r*\* = 1.8. The simulation set up and parameters are identical to those of Figure 5-B.

**Figure 13:**
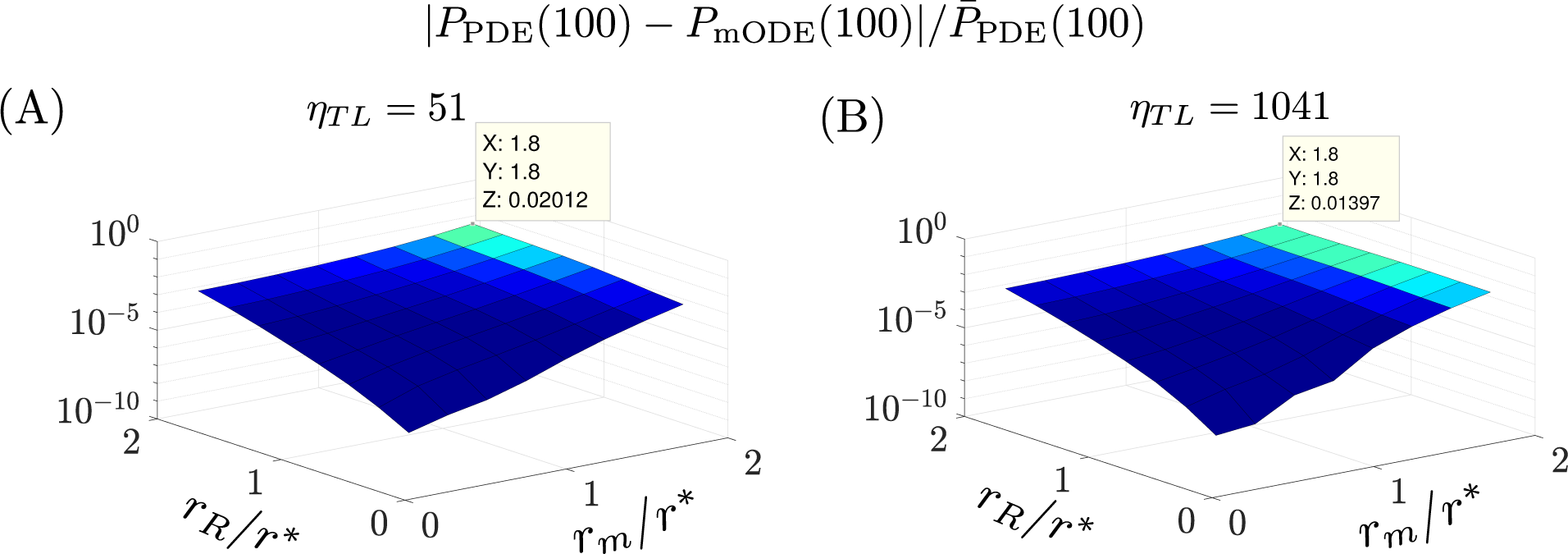
PDE and reduced ODE model agree well in protein production example. Let *P*_PDE_(100) be the steady state protein space averaged concentration predicted by (62) and *P*_ODE_(100) be the steady state protein space averaged concentration predicted by (15). Let *η*_TL_ = 1/(*κ_m_* + *d_m_* + 1 + *γ_m_*). The values of *η*_TL_ are varied by modifying *d_m_* while maintaining *a_m_*/*d_m_* = 1. (A) The relative error for several values of *r_m_* and *r_s_* for *η*_TL_ = 51 =⇒ *d_m_* = 10. (B) The relative error for several values of *r_m_* and *r_s_* for *η*_TX_ = 1041 =⇒ *d_m_* = 1000. For both cases the relative error is less than 2.1% The rest of the simulation set up and parameters are identical to those of Figure 5-B.

**Figure 14:**
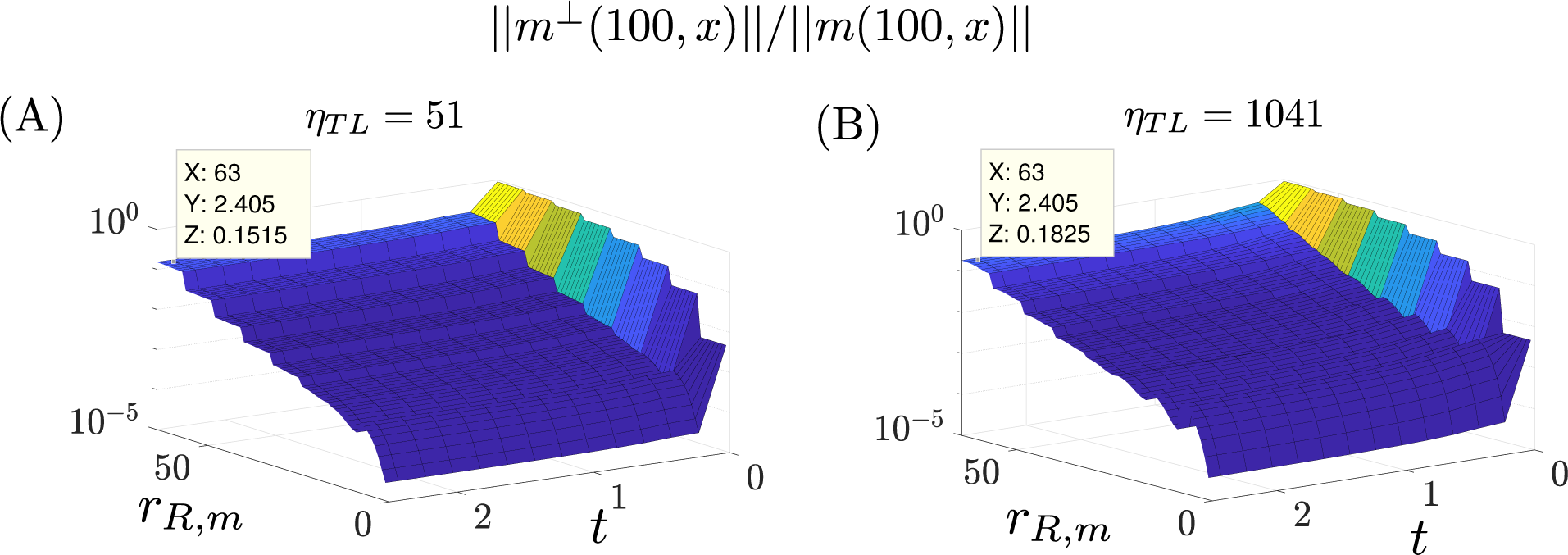
The error in the mRNA approximation for several binding and unbinding speeds between mRNA and ribosome. Let *m*(*t, x*) be as in (62) and 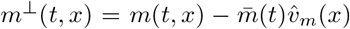 be the a measure of the error in our approximation, where 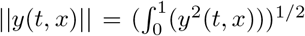, and *η*_TL_ = 1/(*κ_m_* + *d_m_* + 1 + *γ_m_*). The values of *η*_TL_ are varied by modifying *d_m_* while maintaining *a_m_*/*d_m_* = 1. Here *r_R_*, *m* is a sequence corresponding to the mRNA and ribosome pairs from Figure 5-B. Here time nondimensionalized with the time scale associated with dilution. (A) The relative error in time for several values of *r_R_*, *m* for *η*_TL_ = 51 =⇒ *d_s_* = 10. (B) The relative error in time for several values of *r_R_*, *m* for *η*_TL_ = 1041 =⇒ *d_s_* = 1000. For both values of *η*_TL_ the error is high at *t*= 0 since the initial mRNA spatial profile is chosen to be a constant but quickly decays to less than 20%. The rest of the simulation set up and parameters are identical to those of Figure 5-B.

**Figure 15:**
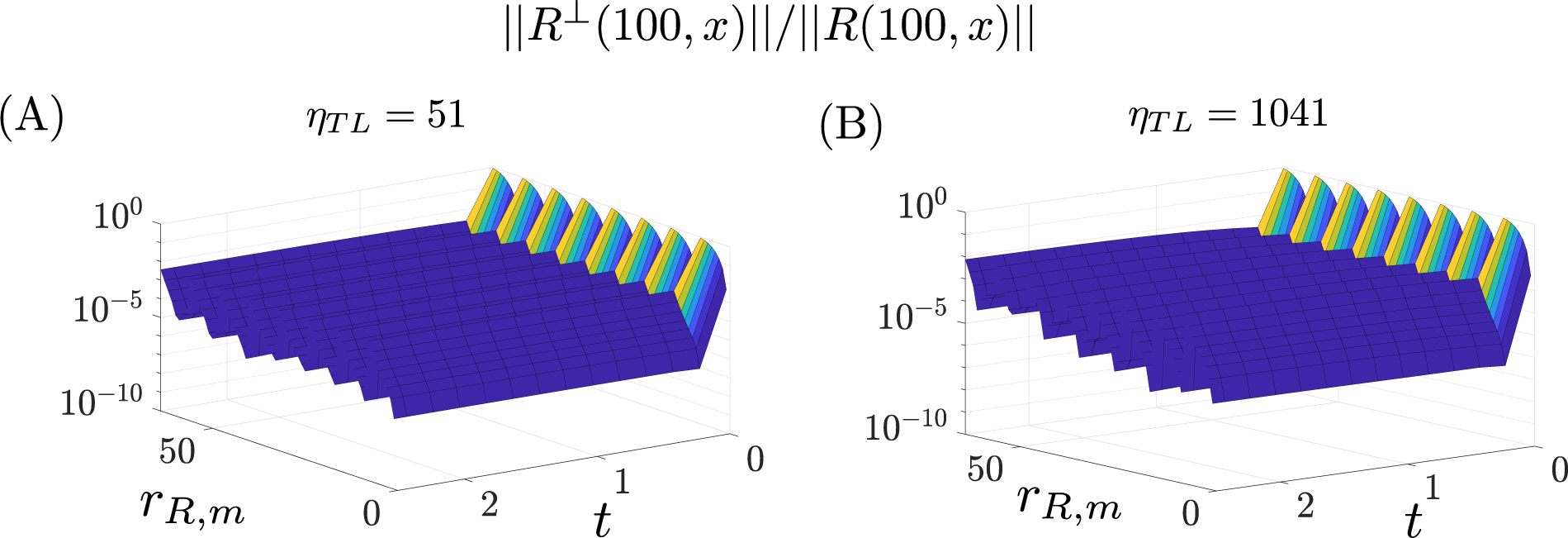
The error in the ribosome approximation for several binding and unbinding speeds between mRNA and ribosome. Let *m*(*t, x*) be as in (62) and 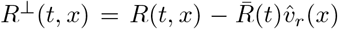 be the a measure of the error in our approximation, where 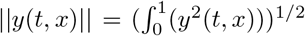, and *η*_TL_ = 1/(*κ_m_* + *d_m_* + + *γ_m_*). The values of *η*_TL_ are varied by modifying *d_m_* while maintaining *a_m_*/*d_m_* = 1. Here *r_R_*, *m* is a sequence corresponding to the mRNA and ribosome pairs from Figure 5-B. Here time nondimensionalized with the time scale associated with dilution. (A) The relative error in time for several values of *r_R_*, *m* for *η*_TL_ = 51 =⇒*d_s_* = 10. (B) The relative error in time for several values of *r_R_*, *m* for *η*_TL_ = 1041 =⇒ *d_s_* = 1000. For both values of *η*_TL_ the error is high at *t* = 0 since the initial mRNA spatial profile is chosen to be a constant but quickly decays to less than 1%. The rest of the simulation set up and parameters are identical to those of Figure 5-B.

The production rate of 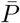 denoted by 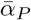 is given by 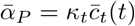. From our analysis in Section 2.2, we expect that 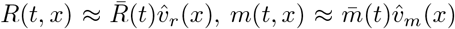, and 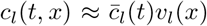 (this is verified computationally in Figure 16), and thus we can estimate 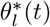 and 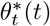 by the constants

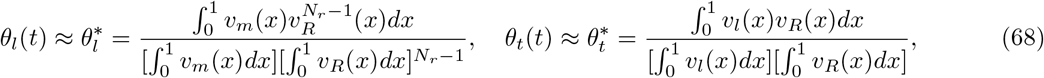

this is verified via simulation in Figure 16)-D.

Let 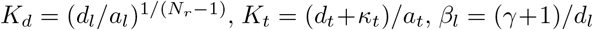, and *β_t_* = (*γ*+1)/(*κ_t_*+*d_t_*), if *β_l_, β_t_, β_t_R/K_t_* ≪ 1 (dilution and mRNA degradation is much slower the rate of ribosome unbinding and *K_t_* is sufficiently large), then a simple expression for the steady state protein is given by

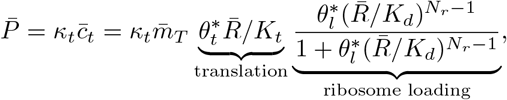

where 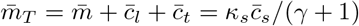 is the total mRNA.

**Figure 16:**
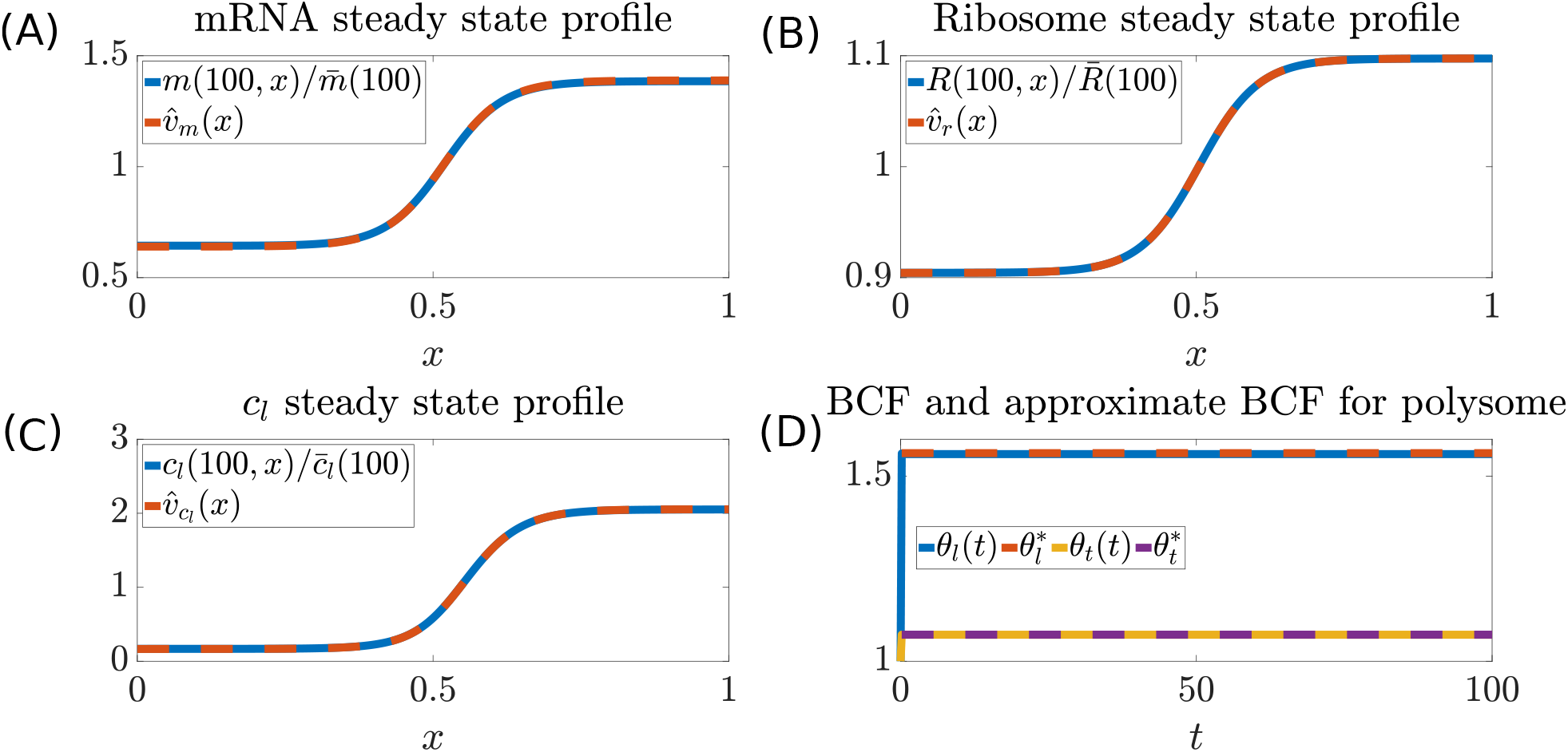
BCF for Polysome with 10 Ribosomes. (A) The steady state mRNA spatial concentration profile (*m*(100, *x*)) predicted by (65) normalized by the space averaged concentration along with the normalized mRNA available volume profile 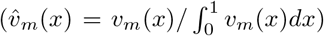. (B) The steady state ribosome spatial concentration profile (*r*(100, *x*)) predicted by (65) normalized by the space averaged concentration along with the normalized mRNA available volume profile 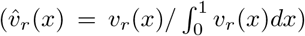. (C) The steady state polysome (loaded with 9 ribosomes) spatial concentration profile (*c_l_*(100, *x*)) predicted by (65) normalized by the space averaged concentration along with the normalized mRNA available volume profile 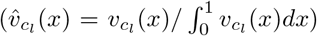. (D) The BCF’s *θ_l_*(*t*) and *θ_t_*(*t*) given by (67) and their constant approximation 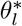 and 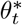 given by (68). For these simulations, we do not model transcription directly but instead set *κ_s_c_s_*(*t, x*) = 1, such that at steady state *m_T_*(*x*) = *m*(*x*) + *c_l_*(*x*) + *c_t_*(*x*) = *κ_s_c_s_*(*t, x*)/(*γ* + 1) = 0.09 for *γ* = 10. The used parameter values are *N_r_* = 10, *χ_m_* = *χ_c_* = 224 *χ_r_* = 1280, *a_t_* = 10, *d_t_* = 10, *κ_t_* = 40, 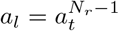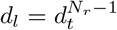, *α_r_*(*t, x*) = 1, *r_m_/r*^*^ = 0.88 and *r_R_/r*^*^ = 0.44.

### 2.8 Transcription Factor Regulation

Intracellular signaling to control gene expression is often done via transcription factors (TFs). In this section we model a general transcription factor architecture where the repressor P_r_ dimerizes to form c_1_ (e.g., TetR dimerizes before targeting gene [32]) and then blocks the transcription of gene *D* that produces protein P. The biochemical reactions corresponding to this process are:

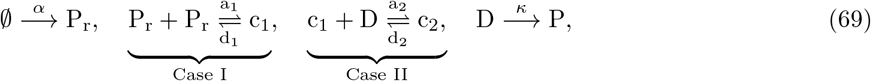

where *α* is the production rate of P_r_, *a*_1_ (*d*_1_) is the association (dissociation) constant to form the c_1_ complex, *a*_2_ (*d*_2_) is the association (dissociation) constant to form the the c_2_ complex, and *κ* is the catalytic rate to produce protein P. Since the repressor P_r_, freely diffuses, the dimerization reaction belongs to Case I. The gene D is spatially fixed and it is repressed by the freely diffusing c_1_, thus this interaction falls under Case II.

**Figure 17:**
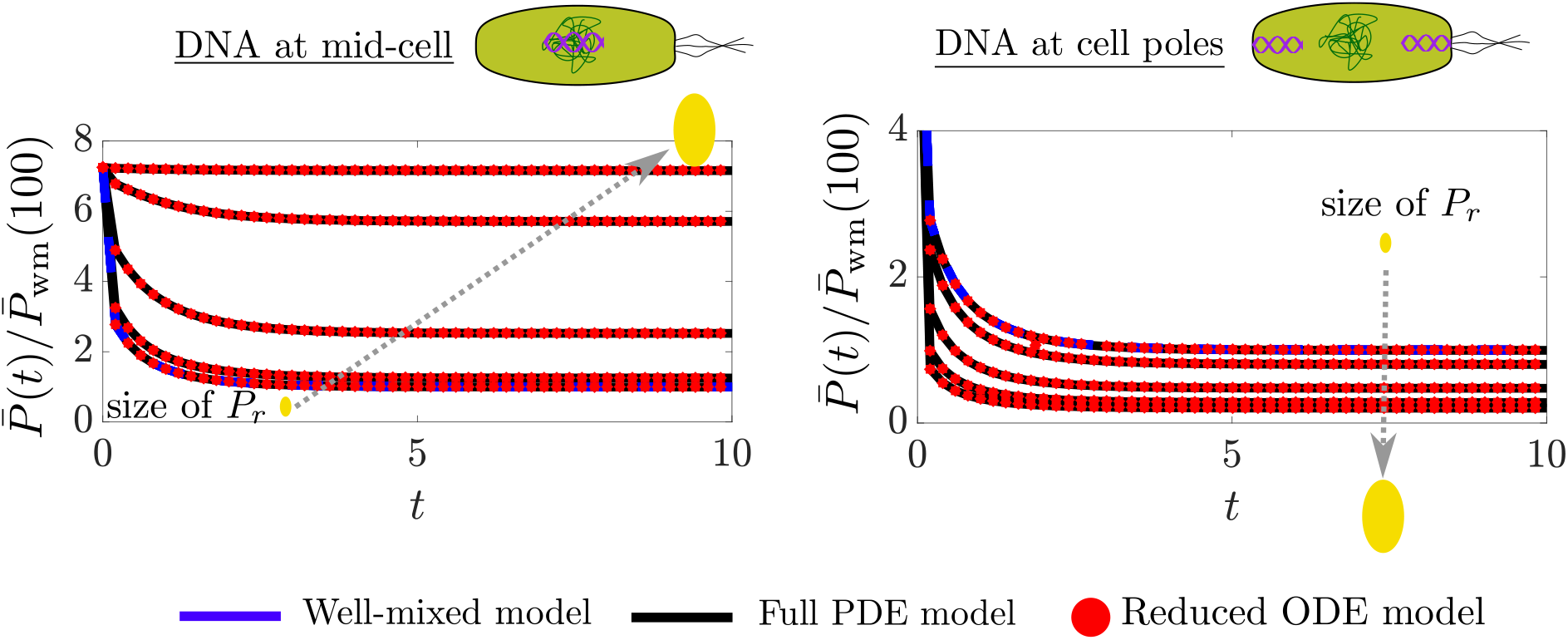
Transcription factor regulation transient response. The transient response corresponding to Figure 6 in the main text when 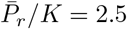

We assume that the total concentration of *D* is conserved, so that *D_T_*(*x*) = *D*(*t, x*) + *c*_2_(*t, x*). The reaction diffusion equations corresponding to (69) are

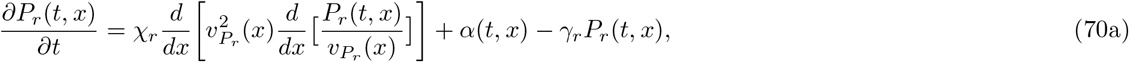

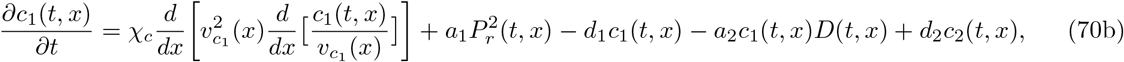

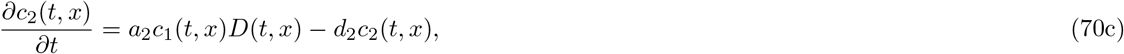

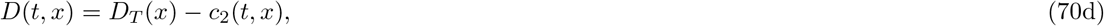

where 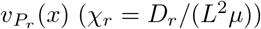 and 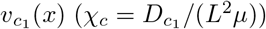 are the available volume profiles (dimensionless diffusion coefficients) of P_r_ and c_1_, respectively, and from (9), 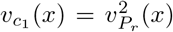. The boundary conditions corresponding to (70) are

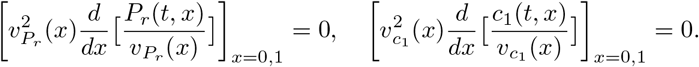

#### Values for dimensionless parameters

The growth rate we used to nondimensionalize the time scales was *μ* = 0.5 1/hr, consistent with the experiments [52]. The length of the cell is about 3*μ*m and thus *L* = 1.5*μ*m [13]. The diffusion coefficient of the transcription factor is taken to be *D_r_* = *D_c_*1 = 0.4*μm*^2^/*s* (that of LacI) [54] and thus *χ_r_* = *χ_c_* = 1280. The transcription factor was assumed to be stale thus *γ_r_* = *μ* The total concentration of D given by 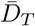 was used to nondimensionalize the other concentration variables such that 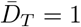.

#### Additional simulation details for Figure 6 in the main text

*D_T_*(*x*) ∝ *e*^−20*x*^ when DNA near mid cell and *D_T_*(*x*) ∝ *e*^20(*x*−1)^ when DNA at cell poles. The transcription factor production was kept roughly spatially constant *α_s_*(*x*) ∝ *e*^−.001*x*^. The binding and unbinding coefficients were chosen to be *a*_1_ = *a*_2_ = 1000 and *d*_1_ = *d*_2_ = 1000 such that the dissociations constants 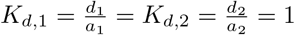.

#### Approxmating BCF from known parameter values

From (9) and (2) in the main text, we observe that the effective radius of gyration of a dimer complex is 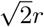, where *r* is the radius of gyration of the individual species. In [55] it was estimated that the radius of gyration for the Tet repressor dimer is 3.1 nm and thus we estimate the radius of gyration of the monomer as 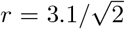. From the expression for *θ*\* given by (24), Figure 4-B, and 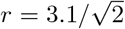, we have that *θ*\* 0.99 and *θ*\* ≈ 1.01, when the target DNA is near mid-cell and the cell poles, respectively. Thus, for TetR, the binding strength between the repressor and the DNA varies by about 1% with respect to a well-mixed model in this parameter range. In [56] it was estimated that the radius of gyration for the Lac repressor tetramer is *r* = 5.3 nm. Assuming that the tetramer is made up of two dimers, then the radius of gyration of each individual dimer is given by 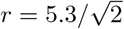. From the expression for *θ*\* given by (24), Figure 4-B, and 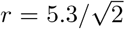, we have that *θ*\* ≈ 0.97 and *θ*\* ≈ 1.03, when the target DNA is near mid-cell and the cell poles, respectively. Thus, for the Lac repressor the binding strength between the transcription factor and the DNA varies by about 3% with respect to a well-mixed model in this parameter range.

**Figure 18:**
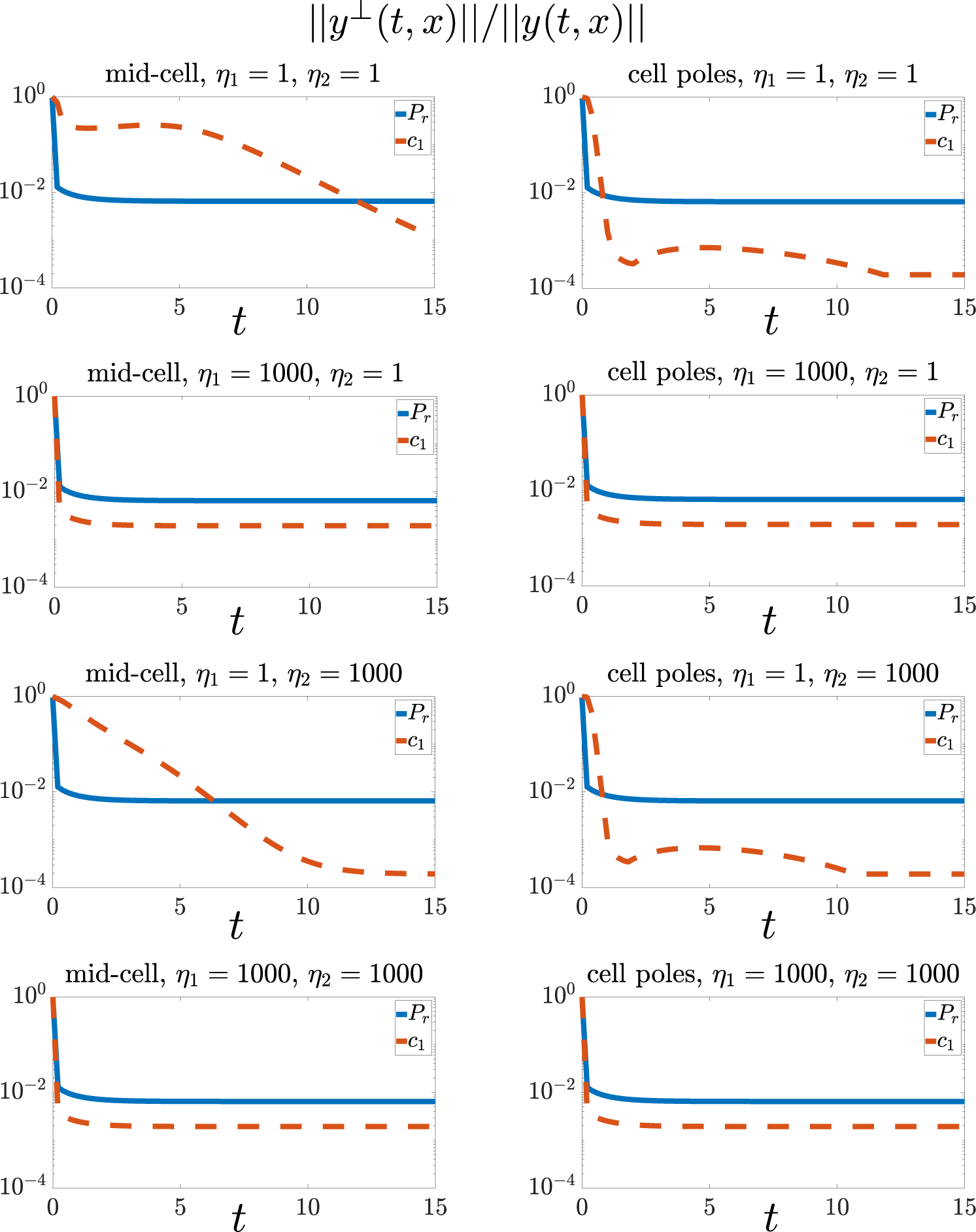
Infinite dimensional dynamics decay in time independently of binding/unbinding speed. Let *y*(*t, x*) be as in (70) (represents *P_r_*(*t, x*) or *c*_1_(*t, x*)) and let 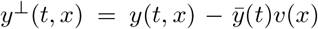 be the a measure of the error in our approximation, where *v*(*x*) is the available volume profile of the species. Let 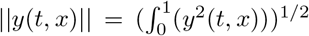, and *η*_2_ = *d*_2_/*μ* and recall that in the simulations we keep *d*_1_/*a*_1_ = *d*_2_/*a*_2_ = 1. We show ‖*y*^⊥^‖/‖*y*‖, the relative error for several values of *η*_1_ and *η*_2_ over dimensionless time (with respect to *μ*) both when the DNA is near mid-cell and the cell poles. The other simulation parameters are identical to those used to generate Figure 6 in the main text. For each time point shown, we took the max relative error with respect to the sizes of P_r_ used to generate Figure 6. The error is high at *t* = 0 since the initial spatial profiles were chosen to be a constant but note that they quickly decay.

**Figure 19:**
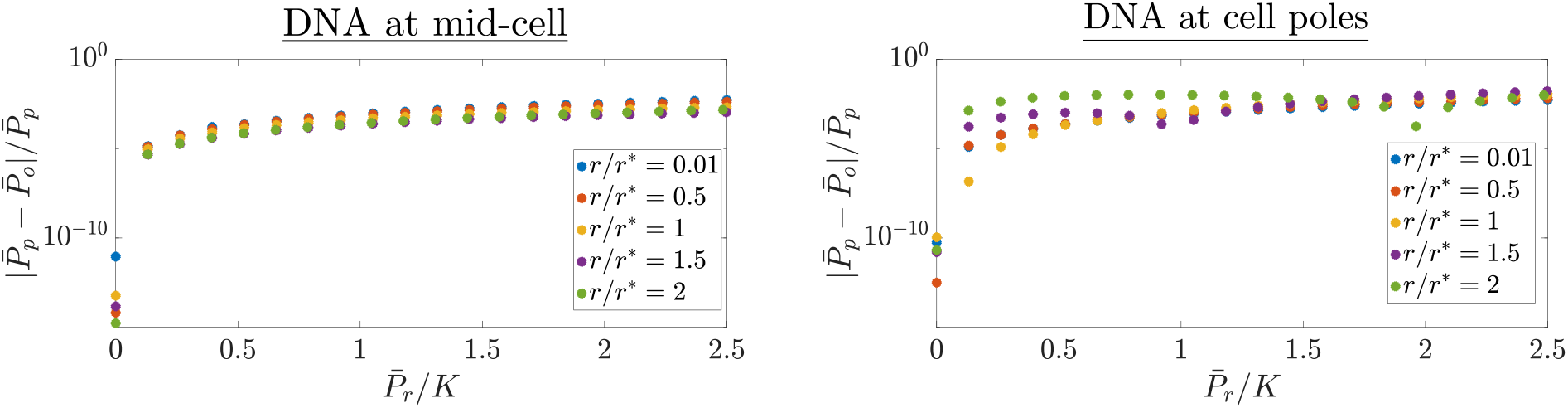
The relative error between the space averaged PDE model and the reduced ODE model from the data in Figure 6 in the main text. The relative error in the steady state space averaged protein for the full PDE model 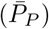 and the reduced model 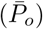 from the data in Figure 6 in the main text. The errors are less than 1.7% both when the DNA is localized at mid-cell and at the cell poles.

While we could not find an exact value for the radius of gyration of the dcas9-gRNA complex, in [57] it was shown that the size of the Cas9-gRNA complex is roughly 10 nm. If we assume this value to be the radius of gyration, from Figure 4-B, we have that the BCF for this complex is *θ*\* = 0.9 and *θ*\* = 1.1 when the target DNA is near mid-cell and the cell poles, respectively. Thus, the binding strength between the Cas9-gRNA complex and the DNA varies s by about 10% with respect to a well-mixed model in this parameter range.

### 2.9 Oscillator

Now we consider the repressor activator clock genetic circuit [35]. This circuit produces sustained oscillations if tuned within an appropriate parameter range [36, 1]. The circuit consists of two proteins P_a_ and P_r_. Protein P_a_, is an activator which dimerizes to form P_a,2_ and then binds to its own gene D_a_ to form complex c_a,1_ to initiate transcription. The dimer P_a,2_ also finds to the gene D_r_, which transcribes P_r_ to form complex c_a,2_ and initiates transcription. Protein P_r_, dimerizes to form P_r,2_ and then represses P_a_ by binding to D_a_ to form complex c_r_. The biochemical equations corresponding to this circuit are:

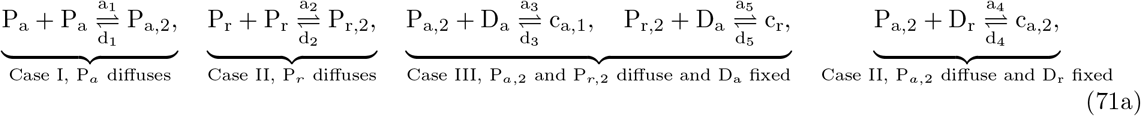

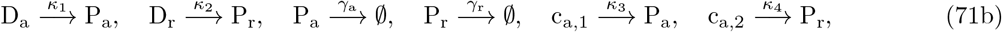

where *a_i_*(*d_i_*) for *i* = 1, …, 5 are association (dissociation) rate constants, *γ_a_* (*γ_r_*) is the degradation rate of P_a_ (P_r_) *κ*_1_ (*κ*_2_) is the basal rate at which gene D_a_ (D_r_) is transcribed, and *κ*_3_ (*κ*_4_) is the rate at which the DNA-transcription-factor complexes are transcribed for D_a_ (D_r_). We assume that the total concentration of D_a_ is conserved, so that *D_a,T_* (*x*) = *D_a_*(*t, x*) + *c*_*a*,1_(*t, x*) + *c_r_*(*t, x*). Similarly, we assume that the total concentration of D_r_ is conserved, so that *D_r,T_* (*x*) = *D_r_*(*t, x*) + *c*_*a*,2_(*t, x*). The spatiotemporal dynamics describing (71) are given by

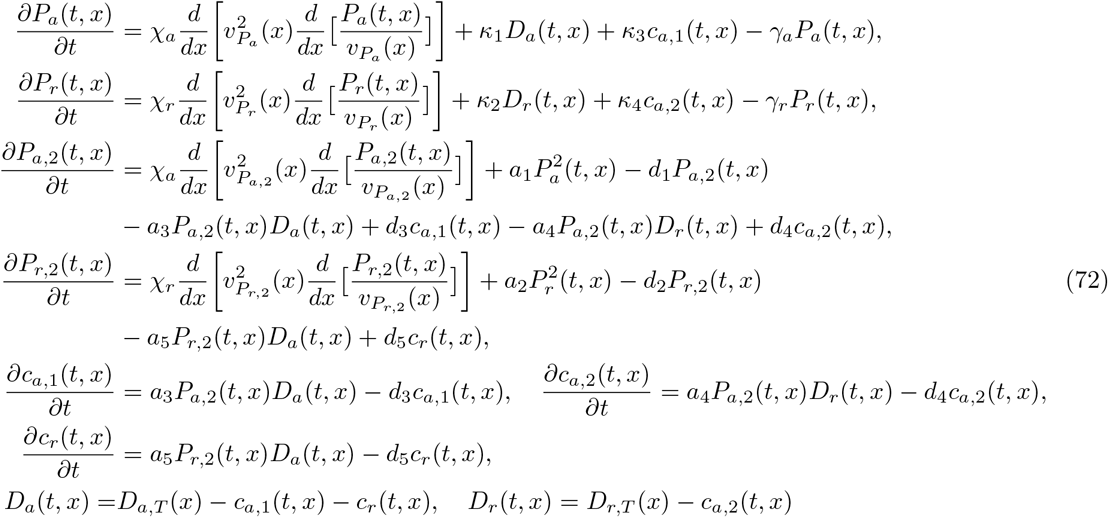

where 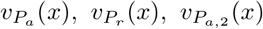, and 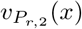 are the available volume profiles of P_a_, P_r_, P_a,2_, and P_r,2_, respectively, *χ_a_* = *D_a_*/(*L*^2^*μ*) is the dimensionless diffusion coefficient of P_a_ and P_a,2_, *χ_r_* = *D_r_*/(*L*^2^*μ*) is the dimensionless diffusion coefficient of P_r_ and P_r,2_. From (9), 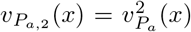 and 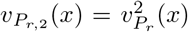. The boundary conditions corresponding to (72) are

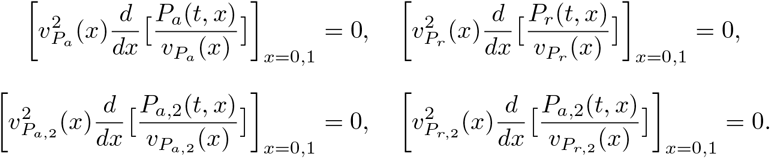

#### Parameters for Figure 7 in the main text

The growth rate we used to nondimensionalize the time scales was *μ* = 0.5 1/hr, consistent with the experiments [52]. The length of the cell is about 3*μ*m and thus *L* = 1.5*μ*m [13]. The diffusion coefficient of the transcription factor is taken to be *D_a_* = *D_r_* = 0.4*μm*^2^/*s* (that of LacI) [54] and thus *χ_a_* = *χ_r_* = 1280. The following dimensionless parameters were chosen such that the well-mixed model displayed sustained oscillations: *a*_1_ = 220, *d*_1_ = 1000, *a*_2_ = 1000, *d*_2_ = 1000, *a*_3_ = 1000, *d*_3_ = 1000, *a*_4_ = 1000, *d*_4_ = 1000, *a*_5_ = 1000, *d*_5_ = 1000, *κ*_3_ = 250, *κ*_1_ = .04 *κ*_4_ = 30, *κ*_2_ = .004 *γ_a_* = 1 *γ_r_* = 0.5. Furthermore, we choose *d_i_* and *a_i_* for *i* = 1, …, 5 large, to demonstrate our results hold even for large binding and unbinding rates. The total concentration of D_a_ which is the same as D_r_ since we assume they are on the same plasmid, is given by 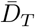 and it was used to nondimensionalize the other concentration variables such that 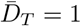. The total DNA spatial profile was chosen as *D_T_*(*x*) ∝ *e*^50(*x*−1)^ to model DNA at cell poles.

### 2.10 Numerical Method Convergence Rate

For the simulation in Figure 5-A when the DNA is localize at the cell-poles and *r_s_/r*\* = 1, we varied the number of spatial nodes used to discretized the spatial domain to demonstrate the convergence rate of our numerical scheme. Let *N* be the number of points used to discretize the spatial domain and let 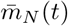 as the space averaged mRNA concentration for that given discretization. We considered when *N* = 1024 to be the true solution *m*_1024_(*t*) and thus define the following relative error

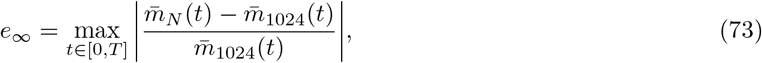

where we took its max value over the time interval of the simulation *t* ∈ [0, *T*] where *T* = 100. The results from this numerical experiment are shown in Figure 22. The convergence rate of our numerical scheme is 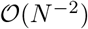 as expected for a second order finite difference numerical scheme.

**Figure 20:**
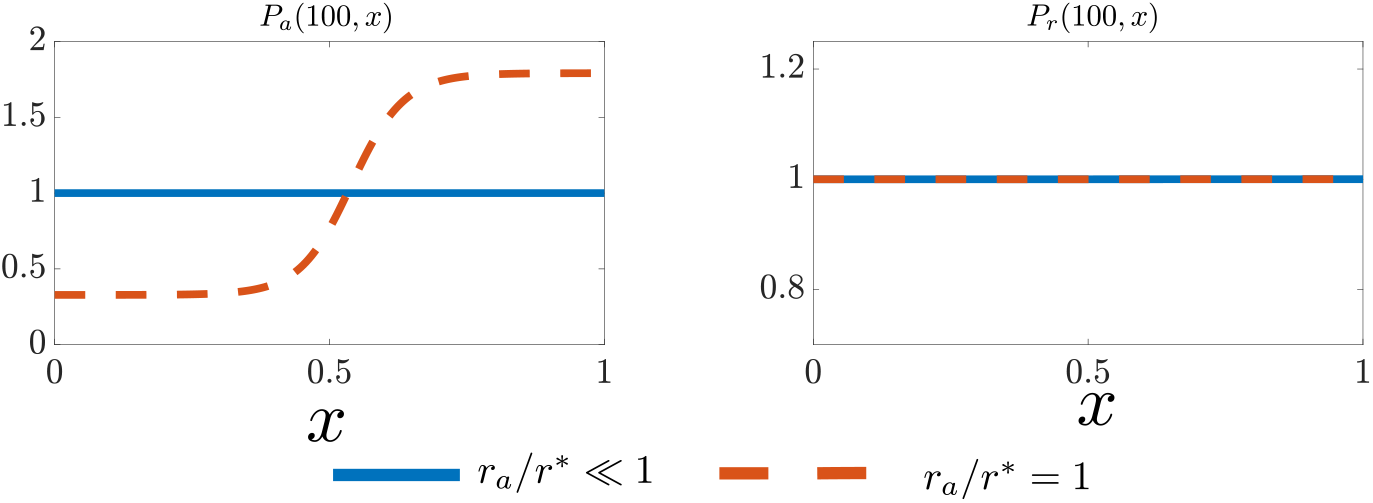
The activator is excluded from the chromosome as its size increases. The steady state spatial profiles normalized by the average values for P_a_ and P_r_, that is 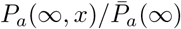 and 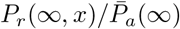 for the results of Figure 7 in the main text. As the size of P_a_ increases it is excluded from the chromosome. The repressor remains homogeneously distributed throughout the cell. The parameter values and simulation details are identical to those of Figure 7 in the main text.

**Figure 21:**
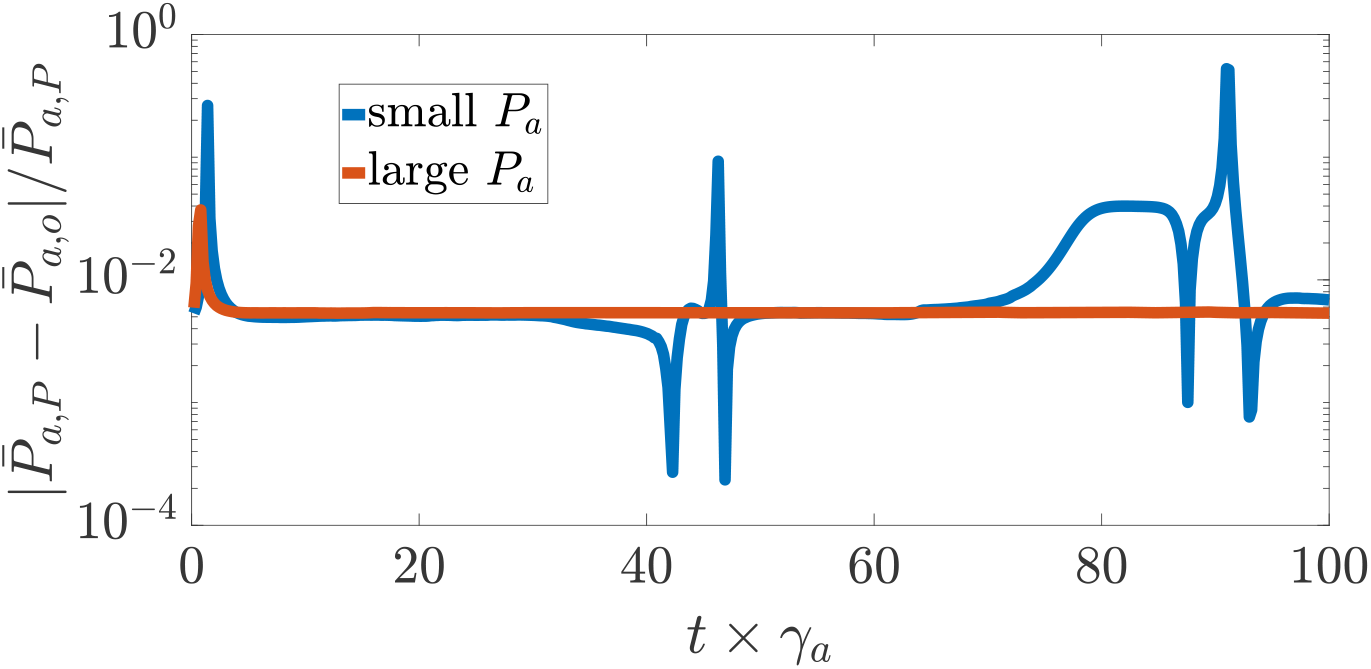
The relative error between the space averaged PDE model and the reduced ODE model from the data in Figure 7 in the main text. The relative error in the steady state space averaged activator protein concentration for the full PDE model 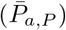 and the reduced model 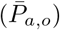 from the data in Figure 7 in the main text. Note that for the case when P_a_ is small, large relative errors occur near when 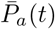 reaches a minimum during each period of oscillation. Otherwise all errors are less than 1%.

### 2.11 Estimating *r*\* from Concentration Profiles and Estimating the BFC

#### Estimate *r*^*^

As discussed in Remark 1 in the main text, we expect the concentration profile of a freely diffusing species to mirror that of the normalized available volume profile. That is, for a freely diffusing species y, with concentration *y*(*t, x*), and available volume profile *v*(*x*), we expect

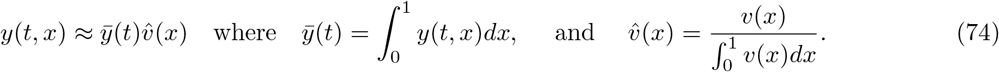

Suppose the radius of gyration of y denoted by *r* is known and as discussed in the main text, we have that 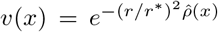. Approximating 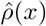 as a step function as in (59), we have that *v*(1) 1 and 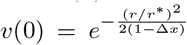, where Δ*x* is the distance between the end of the chromosome and the cell poles (see Figure 1 in the main text). Let let *y*^in^ and *y*^out^ denote the the average concentration inside and outside the nucleotide, respectively, which are given by

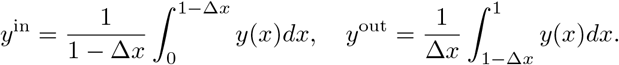

**Figure 22:**
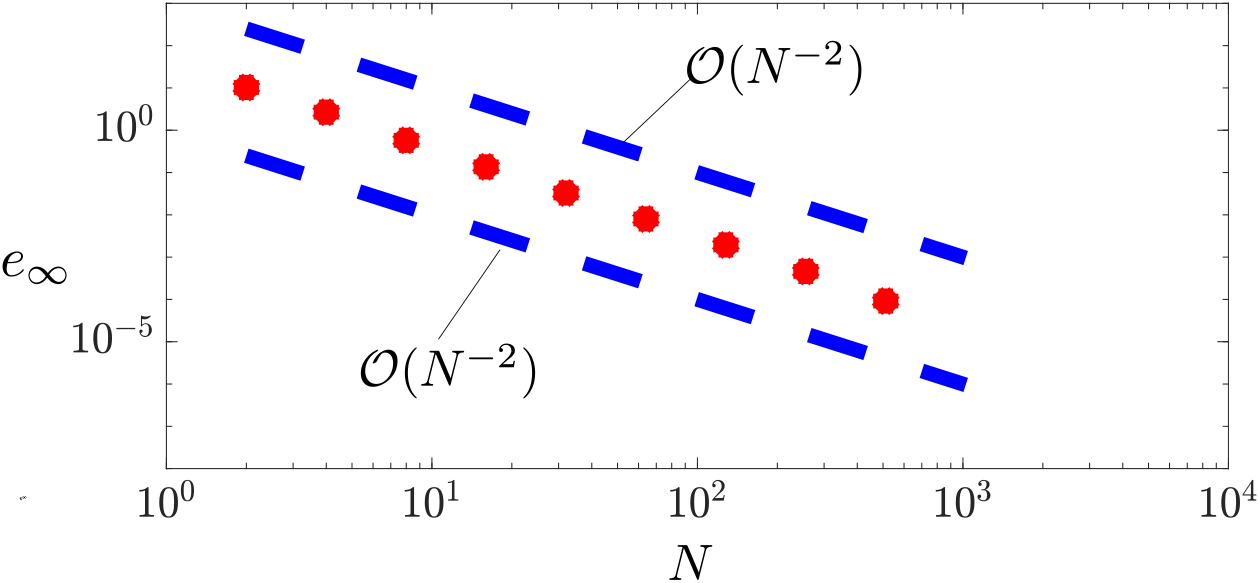
Convergence rate of numerical scheme used to simulate PDEs in this work. For the simulation in Figure 5-A when the DNA is localize at the cell-poles and *r_s_/r*^*^ = 1, we varied the number of spatial nodes used to discretize the spatial domain to demonstrate the convergence rate of our numerical scheme. Let *N* be the number of points used to discretize the spatial domain and let 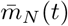 as the space average mRNA concentration for that given discretization. We considered when *N* = 1024 to be the true solution *m*_1024_(*t*) and the relative error *e*_∞_ is given by (73). The relative error is given by the red markers and the blue dashed lines serve as references for 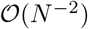 convergence rates.

Let *ψ_y_* = *y*^out^/*y*^in^ and from (74), we have that

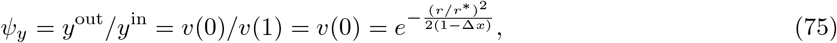

then *r*\* can be estimated assuming one knows Δ*x* that is, how far the dense nucleoid region extends beyond mid-cell. A similar calculation was done in [13], using the fact that the free ribosome concentration is 10% higher at the cell poles than mid-cell. To estimate *ψ* it is sufficient to know the average concentration of a species inside and outside nucleoid region.

#### Estimate the BCF

The BCF provides a measure to determine the extent to which spatial effects modulate the biomolecular dynamics. Therefore, an experimental method to estimate the BCF is desirable. We propose a method that only requires knowing Δ*x* and the concentration of freely diffusing species inside and outside the nucleoid.

Suppose that for Case I and Case III, the concentration of E_i_ is measured inside and outside the nucleoid and denoted by 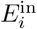 and 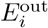, respectively. Similarly, for Case I and Case II, we assume that *S*^in^ and *S*^out^ is measured. If a fluorescence imaging method is used to measure these quantities (as in [14]), then we emphasize that the free E_i_ and S must be measured, not when they are in complex form (c_i_). Let 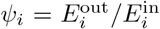 and *ψ_s_* = *S*^out^/*S*^in^. By (75), 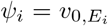 and *ψ_s_* = *v*_0,*S*_ where 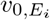 and *v*_0,*S*_ are as in (60). Thus, using (61) we can estimate the BCF for Cases I-III by

- Case I:

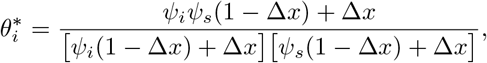
- Case II and Case III:

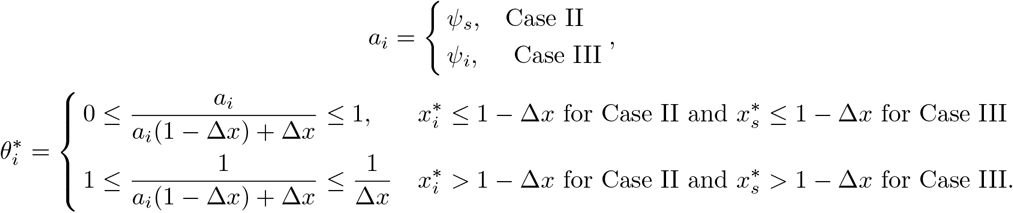

As *r_e,i_/r*\* → (*r_s_/r*\*→ ∞) we have that *ψ_i_* → 0 (*ψ_s_* → 0), thus *ψ_i_* and *ψ_s_* are a measure of the excluded volume effects on E_i_ and S, respectively. Physically, this is expected because when E_i_ is severely expelled from the nucleoid by available volume effects, we have that 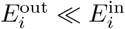 and similarly for S.

### 2.12 Experimental Setups to Verify the Role of Spatial Effects Predicted by Model

A potential experiment to test our hypothesis that genes near the poles our transcribed more effectively than gene near mid-cell, is to measure the rate of transcription (via Quantitative PCR) of a gene under the control of the T7 promoter. This promoter is solely transcribed by the T7 RNAP which specifically targets the promoter, thus this system can be considered orthogonal to the endogenous transcription machinery [58]. By appending random base pairs (BPs) to the sequence of T7 RNAP that do not effect its functionally, we can control its size and thus how much excluded volume effects it experiences. We can then measure the transcription rate of the gene when it is localized in the cell-poles and mid-cell. The results of this experiment should look similar to Figure 5-A in the main text, as the size of the T7 RNAP increases the mid-cell (pole) gene has lower (higher) transcription rate. For the mid-cell localized gene, this can be repeated in parts of the chromosome which are known to be dense to amplify these effects.

To experientially validate our analytical prediction that protein steady state levels will increase with mRNA size, we propose expressing a florescence protein from a plasmid with an appended sequence of base pairs added downstream of the stop codon. The appended sequence should have a low affinity to recruit ribosome such that the amount of ribosomes sequestered by the mRNA are the same as without the appended sequence. Assuming this appended sequence does not affect the lifetime of the mRNA, then it should yield the same functional protein which can be used to quantify the mRNA excluded volume effects. This appended sequence of base pairs will allow us to control the size of the mRNA without increasing its ribosome usage. From our theory, for longer appended sequences, more protein expression is expected.

To validate the hypothesis that a transcriptional repressor regulates genes near the poles more effectively than gene near mid-cell, we propose a genetic circuit on a plasmid expressing a repressor that targets a gene expressing a florescence protein. The transcription factor chosen should be large enough or dimerize to have considerate excluded volume effects. The target DNA expressing protein should be placed on several axial locations in the cell (under the same promoter) achieved by using backbones with different localization profiles and/or different chromosomal integration sites. We should observe that the effective disassociation constant of the repression curve increases as the target genes location is closer to the mid-cell. The disassociation constant is proportional to the amount of repressor necessary to cause the genes expression to decrease by half.

### 2.13 Cell Division: Time Varying Cell Length and Chromosome Profile

As the cell divides it partitions molecular species amongst daughter cells, this along with changes in the cell length cause dilution effects on intracellular concentrations. Furthermore, early in the cell division cycle, the chromosome density is highest mid-cell, but as the cell divides the peak chromosome density tends towards the cell-poles [14, 13] (to distribute genes evenly among daughter cells). From the results in the main text, we expect this temporal changes in the chromosome density will effect the BCF since species are repelled away from regions with high chromosome density via excluded volume effects. In this section, we provide the modeling framework to account for dilution effects and temporal fluctuations in the chromosome density.

#### Dilution effects on spaced average concentrations

Here we demonstrate how cell division and a time varying cell length effects space average concentrations. Let 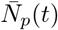 be the total molecular count in a cell population of a molecule of interest (i.e., ribosomes) as the cell expands and divides. To model dilution from cell division, assume that 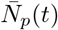 is identically distribute among *N*_cells_(*t*) number of cells such that *N*_cells_(*t*) = *N*_cells_(0)*e^μt^* where *μ* is the cell growth rate and each cell has a volume given by *V_c_*(*t*) = 2*πR*^2^*L*(*t*), where *R* is the cell radius and 2*L*(*t*) is the cell length. The total population volume is then given by *V*(*t*) = *N_c_*(*t*)*V_c_*(*t*) and letting 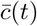 be the number of molecules per total volume (concentration), this quantity is given by 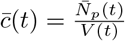, note that this is identical to the concentration per cell volume (since we assume all cells in the population have identical averaged concentrations). This implies that

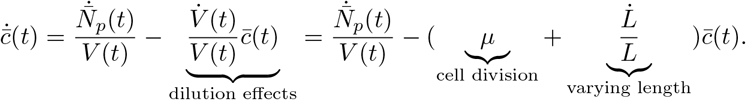

#### Dilution effects on local concentrations

Let *N*(*t, x*) be the number of molecules per unit length of a cell such that that 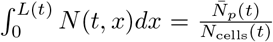 The temporal evolution of *N*(*t, x*) in the presence of dilution and a moving boundary, which introduces an advective term [59] (to account for the extra diffusion as the cell length varies), is given by

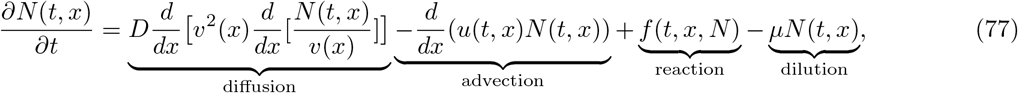

with boundary conditions

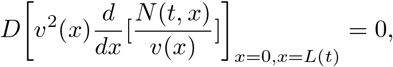

where 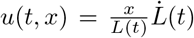 is the velocity of a material point induced by the increase in cell length. Notice that our current finite difference method with a stationary mesh cannot be applied directly to (77), thus we propose the following spatial coordinate transformation 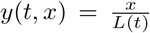 (and to be consistent with the nondimensionalization from the main text), which renders a stationary domain. Let *c*(*t, y*) ≔ *N*(*t, yL*(*t*)) and thus 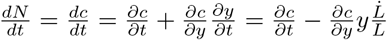 finally

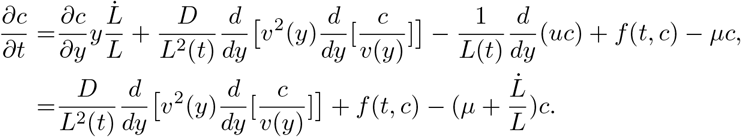

Notice that the effective dilution coefficient is now given by 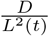, which as expected increases as cell length increases and 2 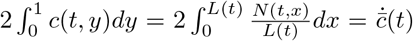, thus the space averaged under this coordinate system provides us the concentration per cell volume. The boundary conditions are

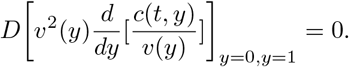

#### Time varying chromosome density

We now model the chromosome density varying in time 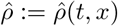 as the cell divides. This implies that the available volume profiles will also depend on time since 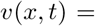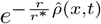, and thus

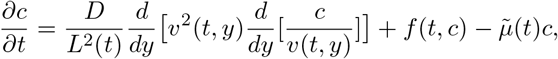

where 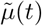 is the effective dilution rate given by

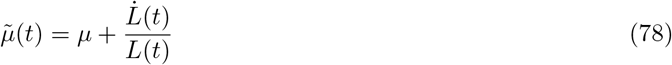

The quantities *L*(*t*) and *v*(*t, x*) will vary with a time scale related to cell growth, for example, let 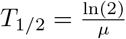 be the cell doubling time, then one possibility is

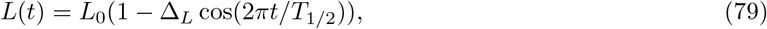

for this choice of *L*(*t*), the effective dilution rate (78) is graphically shown in Figure 23. In [13] it was shown the cell length late in the cell division cycle was 4.4*μm* (compare to its nominal length 3*μm*), thus for our simulations we take Δ_*L*_ = 0.2.

In [14] it was experimentally shown how the chromosome density varies with time and a model for the density in early 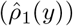 and late 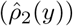 in the cell division process were provided in [13]

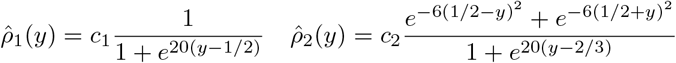

where *c*_1_ and *c*_2_ are chosen such that 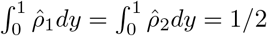. To capture the transition between 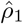 and 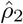 as the cell divides we propose the following model

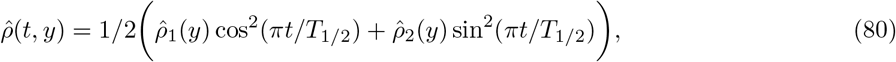

notice that 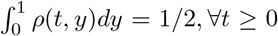. The model for the cell length and the chromosome density is shown in Figure 24. The model for the chromosome density is consistent with experimental observations where late in the division phase the chromosome is in the form of two lobes, where each lobe of DNA will correspond to a daughter cell.

**Figure 23:**
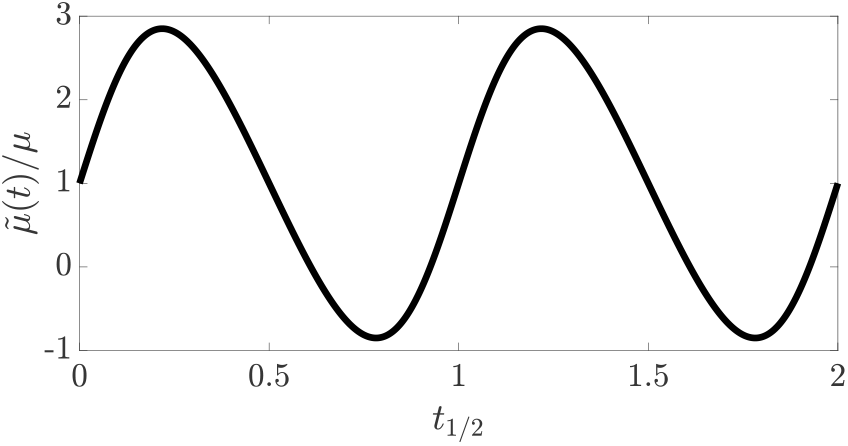
Varying cell length modulates dilution rate. The effective dilution rate (78) is given for *L*(*t*) given by (79) and Δ_*L*_ = 0.2, where *t*_1/2_ is time normalized by the cell doubling time (*t*_1/2_ = *t/T*_1/2_).

**Figure 24:**
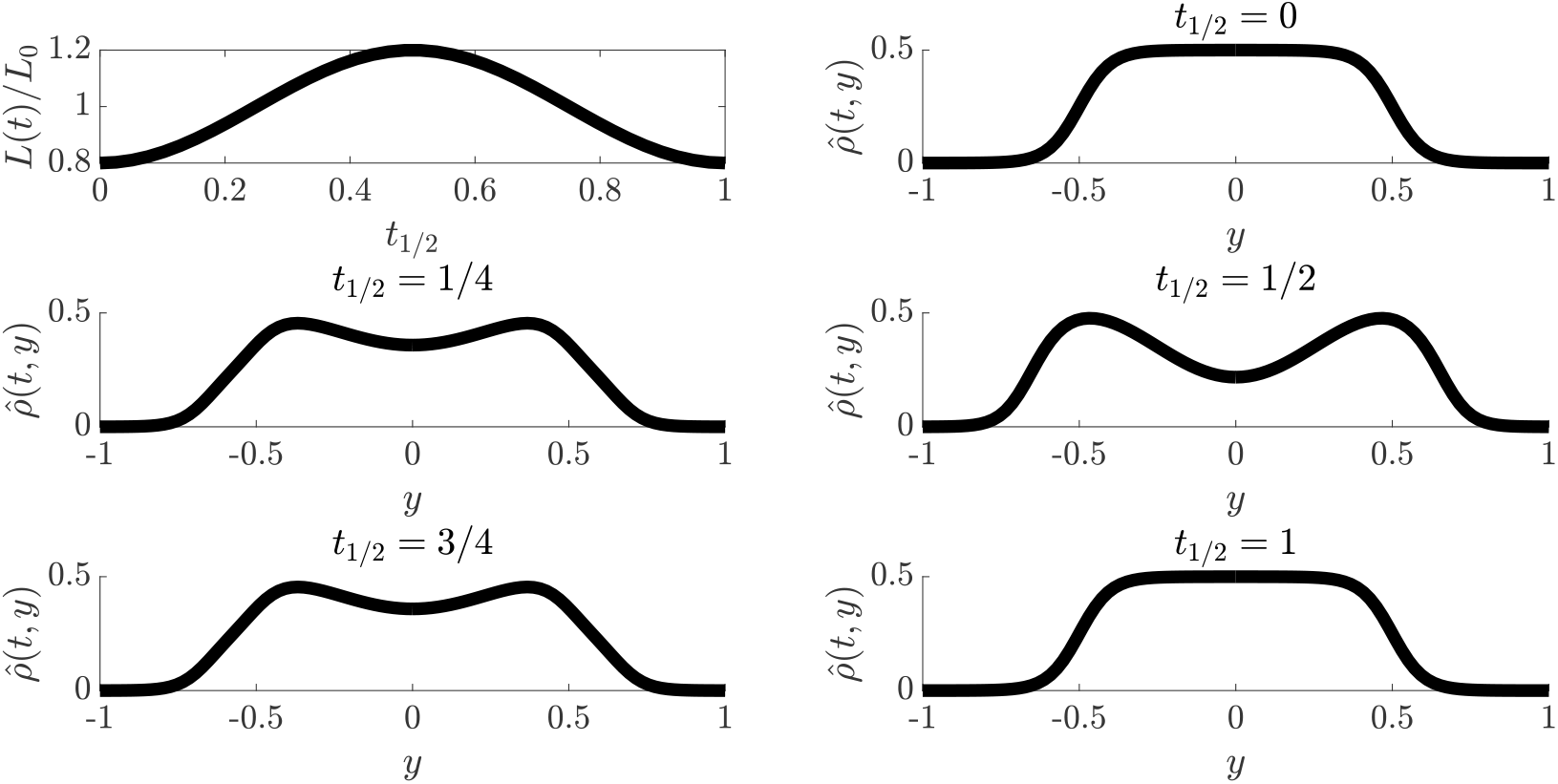
The cell length and chromosome density varies over time. The normalized cell length *L*(*t*)/*L*_0_ (79) with Δ_*L*_ = 0.2 and the chromosome density 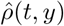 (80) shown over one cell division cycle, where *t*_1/2_ is time normalized by the cell doubling time. The model for the chromosome density is consistent with experimental observations [14, 13] where late in the division phase the chromosome is in the form of two lobes, where each lobe of DNA will correspond to a daughter cell.

#### Time scale separation

When the scale associated with diffusion is much fast than dilution *D/L*_0_ ⨠ *μ* (and any other time scale associated with the reaction dynamics), we can treat *L*(*t*) and *v*(*t, y*) as constant in time when performing model reduction as in Section 1.3 in the main text, thus we expect (similar to the results of the main text)

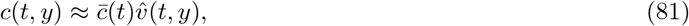

where 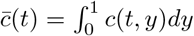 and 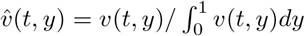. So all of our previous analysis still holds except that the BCF will vary slowly (slow with respect to the time scale of diffusion) as the cell divides.

**Example:** We verify via simulations the prediction that (81) holds and that the BCF can be treated as a slowly (with respect to diffusion) varying parameter. Consider the simple bimolecular reaction:

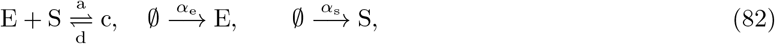

with dynamics given by

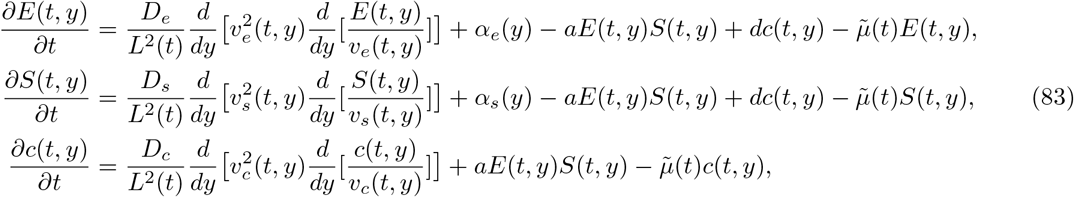

where *α_e_*(*y*) and *α_s_*(*y*) are the production rates of E and S, respectively. The space averaged dynamics 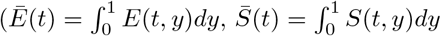 and 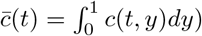 are given by

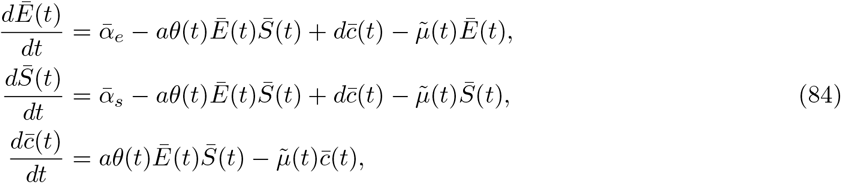

where the BCF is given by

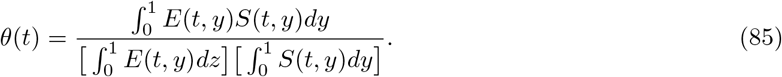

We first focus on the *E*(*t, y*) dynamics when *a* = 0 and *d* = 0

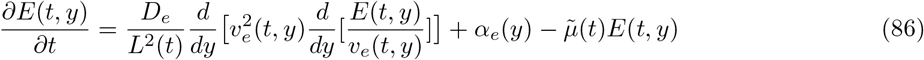

to show the effects of having time varying dilution 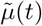 and available volume profiles on the expression level. Figure 25 shows how the space averaged concentration 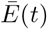 is modulated by the time varying dilution 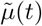. We observe that the concentration reaches a periodic steady state centered at unity where the oscillations have a period that coincides with the doubling time. Furthermore, in Figure 26, we verify that

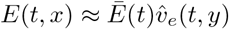

where 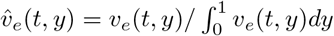 as expected (since diffusion much faster than dilution *D_e_*/(*L*_0_*μ*) ⨠ 1). Thus, even with a time varying *v_e_*(*t, y*), the enzyme will be expelled from the chromosome to areas of higher available volume.

**Figure 25:**
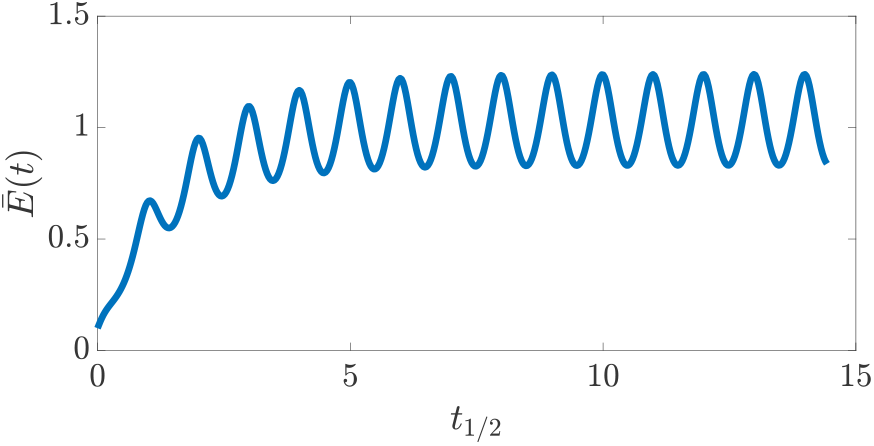
Enzyme expression as cell length varies. The space averaged enzyme expression (86) there is no binding/unbinding with S (*a* = 0 and *d* = 0). The oscillations arise due to changes in the cell length during cell division. The simulation parameters are: *r_e_/r*\* = 2 *μ* = 1 *D_e_/*(*L*_0_*μ*) = 13 10^3^ *α_e_*(*y*) = 1, Δ_*l*_ = 0.2.

Next, we demonstrate how the binding dynamics are effected by having a time varying available volume profile, therefore *a* ≠ 0 in (83). Similar to “Case 1” in the main text, we consider the case when *D_e_, D_s_, D_c_* ≠ 0 (all species freely diffuse), where we expect the BCF to be approximated by

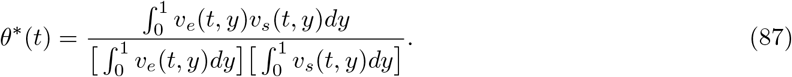

**Figure 26:**
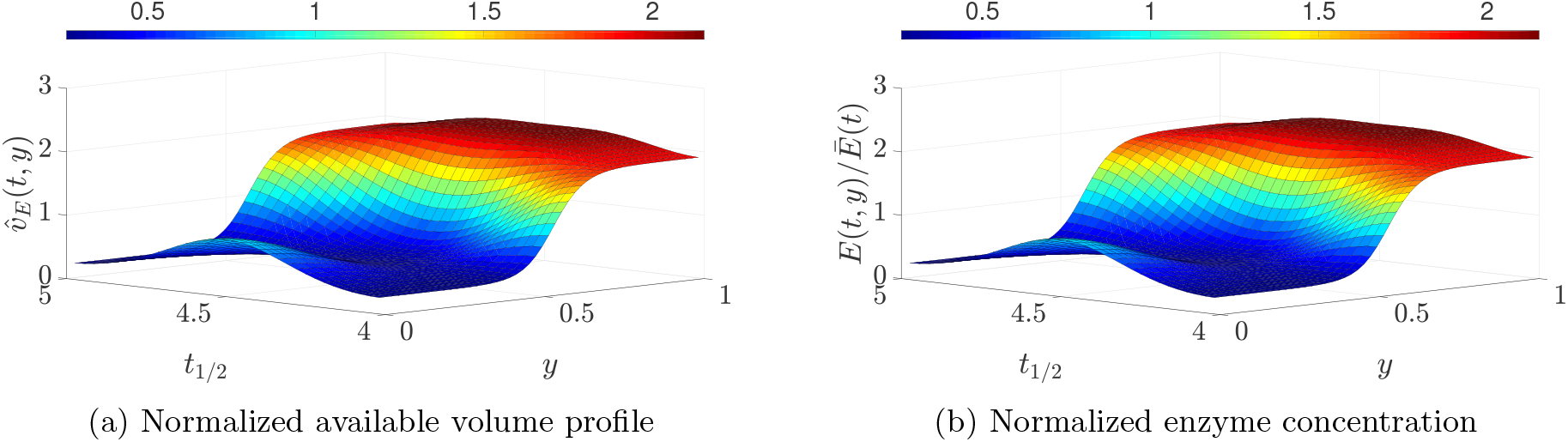
The normalized enzyme spatial profile matches that of its normalized available volume profile. The enzyme concentration spatial profile normalized by its space average shown over one cell division cycle after four cell division cycles (“steady state”) matches its available volume as expected (86). The simulation parameters are: *r_e_/r*\* = 2 *μ* = 1 *D_e_*/(*L*_0_*μ*) = 13 × 10^3^ *α_e_*(*y*) = 1, Δ_*l*_ = 0.2.

This is verified in Figure 27, where *θ*(*t*) given by (85) and *θ*\*(*t*) given by (87) are shown after three doubling times and are shown to be in good agreement. The BCF varies periodically in time (with the period consistent with the doubling time) and oscillates near a nominal value of 1.5 with amplitude 0.04.

Next we look at the the case when S and c are spatially fixed (*D_s_* = *D_c_* = 0) and localized near *y*\*, which is similar to “Case 2” in the main text. For this scenario we expect the BCF to be approximated by

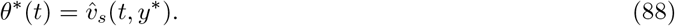

The results are shown in Figure 28 when *y*\* = 0 and *y*\* = 1 after three doubling times, there is good agreement between the BCF and its approximation. When *y*\* = 0 (S localized near mid-cell), the BCF is less than unity and oscillates near a nominal value of 0.55 with amplitude 0.3. When *y*\* = 1 (S localized near the cell poles), the BCF is greater than unity and oscillates near a nominal value of 2 with amplitude 0.1.

These results suggest that the BCF for a species localized near mid-cell will vary significantly as the cell density varies during cell division. This is expected because as shown in Figure /?, the chromosome density is initially high near mid-cell but decreases by half as the cell divides, thus no longer excluding from that region.

**Figure 27:**
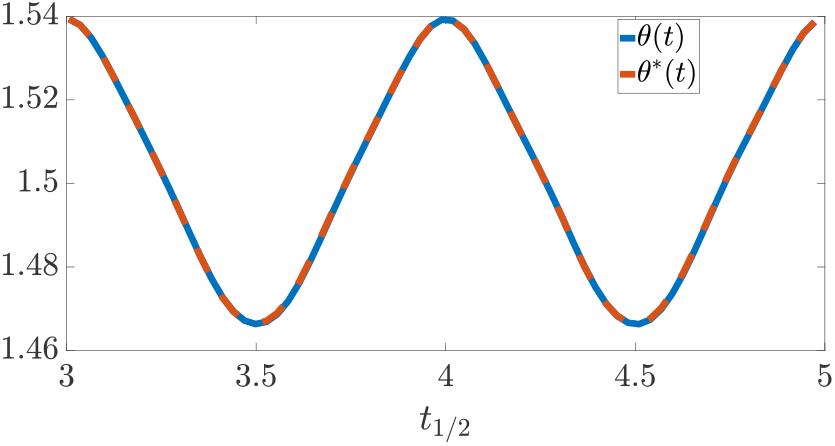
The BCF for the case when all species freely diffuse. The binding correction factor *θ*(*t*) (85) and its approximation *θ*\* (87) over two cell division cycles. The BCF oscillates around a nominal value of 1.5 with amplitude 0.04 and period consistent with the doubling time. The simulation parameters are *r_e_/r*\* = *r_s_/r*\* = 2, 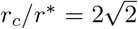, *μ* = 1, *D_e_/L*_0_ = *D_s_/L*_0_ = *D_c_/L*_0_ = 13 × 10^3^ *α_e_*(*y*) = *α_s_*(*y*) = 1, Δ_*l*_ = 0.2, *d* = 100, *a* = 100.

### 2.14 Exclusion Effects from Plasmid DNA Density

The genome of *E. coli* MG1655 has 4.6 Mbp [60]. Comparatively, a single plasmid can have .01 Mbp a copy number as high as 500-700 (e.g. pUC19). Therefore, the total plasmid and chromosome basepair count may be comparable in applications with high copy number plasmids. In these applications, it may be necessary to account how plasmid DNA repels freely diffusing species and “excludes” them. To do so we modify our model of the DNA density 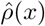 as shown in Figure 29 to account for plasmid DNA. For the DNA density profiles from Figure 29, we calculate the approximate BCF *θ*\* (13), these are shown in Figure 30. For Case 1 where the reactants freely diffuse. We observe that the BCF decreases as the plasmid DNA density increases (as shown in Figure 29). This occurs because as the plasmid DNA increases the overall density profile becomes more uniform. Note that when the plasmid DNA is sufficiently high to render an almost uniform DNA density profile, the BCF is unity as expected. For Case 2, where one reactant freely diffuses and the other is fixed at *x*\*. As the plasmid density increases (as shown in Figure 29) we observe that the BCF decreases at the cell poles (as expected since species are excluded from dense plasmid DNA mesh) and increases at region near *x*\* ≈ 0.65 (where there is minimal overlap between chromosome and plasmid DNA). When the plasmid DNA density is similar to that of the chromosome rending a uniform DNA distribution, we observe that the BCF is unity everywhere (as expected since there are no exclusion effects).

**Figure 28:**
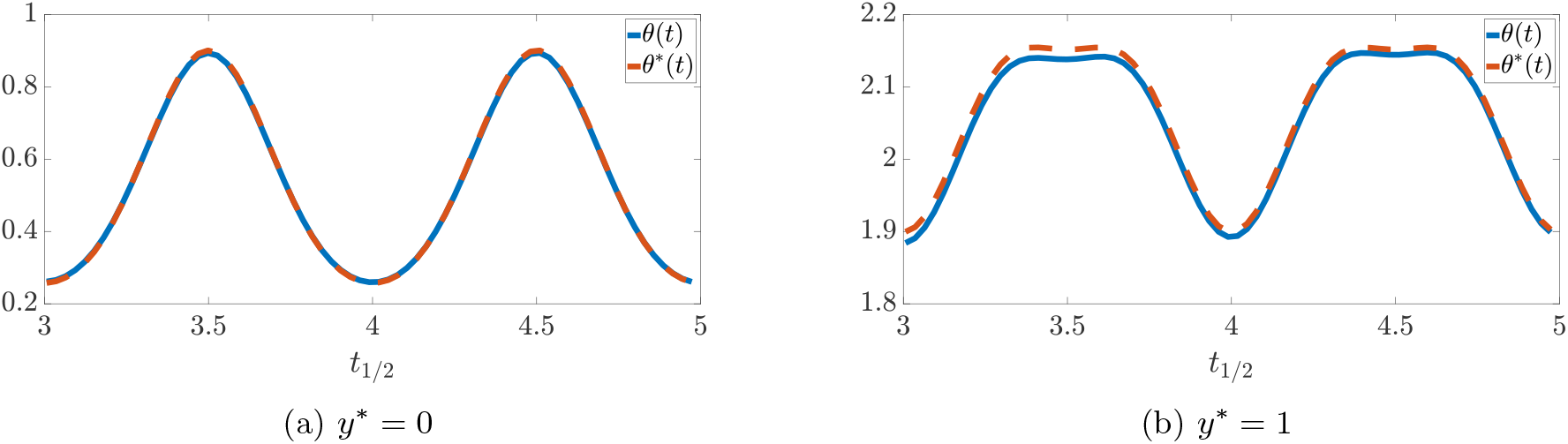
The BCF for the case when S is spatially fixed at *y*\*. The binding correction factor *θ*(*t*) (85) and its approximation *θ*\* (88) over two cell division cycles. The BCF oscillates with a period consistent with the doubling time. When *y*\* = 0 (S localized near mid-cell), the BCF is less than unity and oscillates near a nominal value of 0.55 with amplitude 0.3. When *y*\* = 1 (S localized near the cell poles), the BCF is greater than unity and oscillates near a nominal value of 2 with amplitude 0.1. This is shown for a molecule localized near mid-cell *y*\* = 0 and near the cell poles *y*\* = 1. The simulation parameters are *r_e_/r*\* = *r_s_/r*\* = 2, 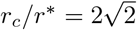 *μ* = 1, *D_e_/L*_0_ = *D_s_/L*_0_ = *D_c_/L*_0_ = 13 × 10^3^ *α_e_*(*y*) = *α_s_*(*y*) = 1, Δ_*l*_ = 0.2, *d* = 100, *a* = 100.

**Figure 29:**
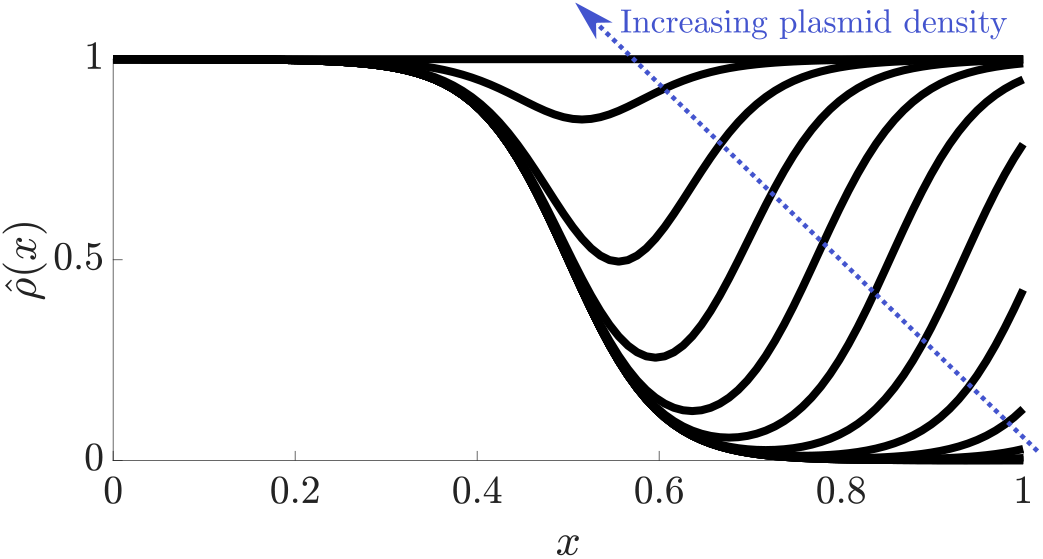
DNA density with plasmid contributions. The DNA density now includes contributions from plasmid DNA. We show several profiles with increasing plasmid density. For this results we had 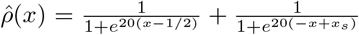 where *x_s_* ∈ [1/2,3/2] is the parameter we varied to get difference plasmid densities (*x_s_* = 3/2 lowest plasmid density and *x_s_* = 1/2 highest plasmid density).

**Figure 30:**
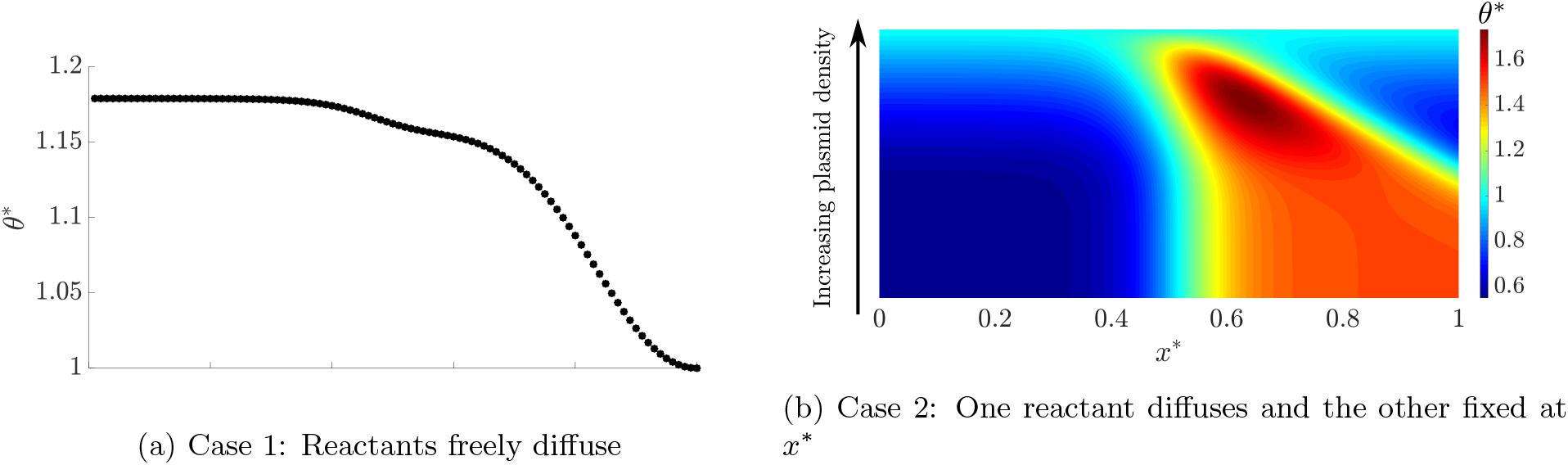
The approximate BCF *θ*\* when plasmid DNA is accounted for. (a) Case 1 where the reactants freely diffuse. We observe that the BCF decreases as the plasmid DNA density increases (as shown in Figure 29). This occurs because as the plasmid DNA increases the overall density profile becomes more uniform. Note that when the plasmid DNA is sufficiently high to render an almost uniform DNA density profile, the BCF is unity as expected. (b) Case 2 where one reactant freely diffuses and the other is fixed at *x*\*. As the plasmid density increases (as shown in Figure 29) we observe that the BCF decreases at the cell poles (as expected since species are excluded from dense plasmid DNA mesh) and increases at region near *x*\* ≈ 0.65 (where there is minimal overlap between chromosome and plasmid DNA). When the plasmid DNA density is similar to that of the chromosome rending a uniform DNA distribution, we observe that the BCF is unity everywhere (as expected since there are no exclusion effects). The simulation parameters are *r/r*\* = 1 and 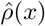 as in Figure 29.

